# Endowing Universal CAR T-cell with Immune-Evasive Properties using TALEN-Gene Editing

**DOI:** 10.1101/2021.12.06.471451

**Authors:** Sumin Jo, Shipra Das, Alan Williams, Anne-Sophie Chretien, Thomas Pagliardini, Aude Le Roy, Jorge Postigo Fernandez, Diane Le Clerre, Billal Jahangiri, Isabelle Chion-Sotinel, Agnes Gouble, Mathilde Dusséaux, Roman Galetto, Aymeric Duclert, Emanuela Marcenaro, Raynier Devillier, Daniel Olive, Philippe Duchateau, Laurent Poirot, Julien Valton

**Author notes:** Co-first Authors (Sumin Jo, Shipra Das, Alan Williams).

## Abstract

Universal CAR T-cell therapies are poised to revolutionize cancer treatment and to improve patient outcomes. However, realizing these advantages in an allogeneic setting requires universal CAR T-cells that can kill target tumor cells, avoid depletion by the host immune system, and proliferate without attacking host tissues. Here, we describe the development of a novel immune-evasive CAR T-cells scaffold that evades NK cell and alloresponsive T-cell attacks and imparts efficient antitumor activity *in vitro* and *in vivo*. This scaffold could enable the broad use of universal CAR T-cells in allogeneic settings and holds great promise for future powerful clinical applications.

## Introduction

Universal Chimeric Antigen Receptor (CAR) expressing T-cells have great potential to democratize and improve the treatment of cancer patients worldwide. Reasons for such potential are multiple but all stem from the source of the biological material used to produce them. Because universal CAR T-cells are engineered out of third-party healthy donor T-cells, donors can be carefully selected for potency and cells can be manufactured, formulated, and controlled thoroughly before being adoptively transferred to multiple patients in an allogeneic setting.

To realize their full potential in an allogeneic setting, universal CAR T-cells must not induce two detrimental and potentially toxic phenomena: the Graft versus Host (GvH) reaction and the Host versus Graft (HvG) reaction. The GvH reaction can be readily addressed by the transient or constitutive inactivation of T cell receptor αβ (TCRαβ) expression in CAR T-cells^1,2,3,4^. In contrast, preventing depletion of CAR T-cells due to the HvG reaction is less straightforward. The preconditioning regimen that is commonly used to lymphodeplete patients prior CAR T-cell transfer (cyclophosphamide and fludarabine)^5,6^ can delay the HvG reaction and create a first window of opportunity for CAR T-cell engraftment. However, because this lymphodepletion is transient, it may not fully prevent the HvG reaction. In addition to human leukocyte antigen (HLA) matching between CAR T-cell donors and recipients^7,8^, two main engineering strategies have been thoroughly assessed for their ability to further inhibit or delay the HvG reaction. The first strategy relies on developing drug-resistant CAR T-cells wherein TCRαβ and genes that modulate sensitivity to lymphodepleting drugs (CD52 and dCK which are responsible for alemtuzumab binding and fludarabine metabolism^2,9^, respectively) are inactivated. This strategy, which is designed to allow for CAR T-cell engraftment and proliferation under prolonged lymphodepletion of the host, showed encouraging antitumor potency in clinical trials when alemtuzumab is used as part of the lymphodepletion regimen ^10,11^. The second strategy relies on the genetic inactivation of beta-2 microglobulin (B2M) in CAR T-cells. The inactivation of B2M prevents the expression of the HLA Class-I surface marker that is responsible, in part, for the host T-cell-mediated HvG reaction^12–14^. TCRαβ(-) HLA-ABC(-) CAR T-cells can be efficiently generated via several gene editing methods, are hypoimmunogenic with respect to alloresponsive T-cells^13–16^, and are currently being evaluated in a phase I clinical trial.

This second approach, often presented as the next generation of universal CAR T-cell therapy, offers the potential advantage of extending the engraftment of CAR T-cells without relying on prolonged lymphodepletion which increases the risk of opportunistic infections and reduces any potential benefits provided by endogenous immune effectors. However, while highly attractive, this strategy is likely to be impaired by the presence of host NK cells, which recognize and readily deplete HLA-ABC(-) T-cell through the missing-self response^12^. Clinical studies investigating the antitumor potential of HLA-ABC(-) CAR T-cells will be informative but it is difficult to predict whether NK cells will recover in number and fitness to mediate a missing-self response following lymphodepletion. Therefore, realizing the full potential of universal HLA-ABC(-) CAR T-cells will require new engineering strategies that can enable them to evade host NK cell attacks.

Here, we report the development of an immune-evasive universal CAR T-cell scaffold named ΔTRAC_CAR_ΔB2M_HLAE_, which incorporates disruptive insertions of a CAR and HLAE, a non-polymorphic NK inhibitor^17^ into the TRAC and B2M loci, respectively. Using a combination of multiplex TAL Effector Nucleases (TALEN) and recombinant adeno-associated virus 6 (AAV6) treatments, we show that ΔTRAC_CAR_ΔB2M_HLAE_ can be efficiently produced, displays antitumor activity, and resists primary allo-responsive T-cells and primary NK cells sourced from healthy donors and from acute myeloid leukemia (AML) patients. Taken together, these findings support further development of ΔTRAC_CAR_ΔB2M_HLAE_ as a novel immuno-evasive CAR T-cell scaffold that is compatible with adoptive cell transfer in allogeneic settings.

## Results

### Efficient production of ΔTRAC_CAR_ΔB2M_HLAE_ by disruptive insertions of CAR and HLA-E at the TRAC and B2M loci, respectively

Multiple approaches could be used to inhibit the cytolytic function of NK cells toward HLA-ABC(-) T-cells^18–20^. To develop a robust and straightforward approach that is compatible with clinical applications, we chose to use the B2M gene to re-express the engineered NK inhibitor HLA-E, via a disruptive gene insertion approach (i.e., a knock-out by knock-in). Based on the approach of Crew *et al*^17^, we derived an AAV6 promoter-less DNA repair matrix that is specific for the B2M locus; we named this matrix HLAE_m_. The HLAE_m_ matrix includes a nonameric peptide derived from HLA-G (VMAPRTLIL^17,21,22^), codon-optimized B2M and HLA-E domains covalently linked together by GS linkers (HLA-E^17,23^, Fig. 1a), and 300-bp left and right homology arms that are specific for exon 1 of the B2M locus. HLAE_m_ was designed to be used in combination with a B2M-specific TALEN that would inactivate the endogenous B2M gene and repurpose its regulatory elements and reading frame to express the engineered HLA-E. In the following, the B2M-specific TALEN and HLAE_m_ will be used in combination with the TRAC TALEN and the AAV6 promoter-less CAR matrix (CAR_m_, Fig.1a), described earlier to mediate the disruptive insertion of a CAR construct at the TRAC locus^24^.

**Figure 1.**
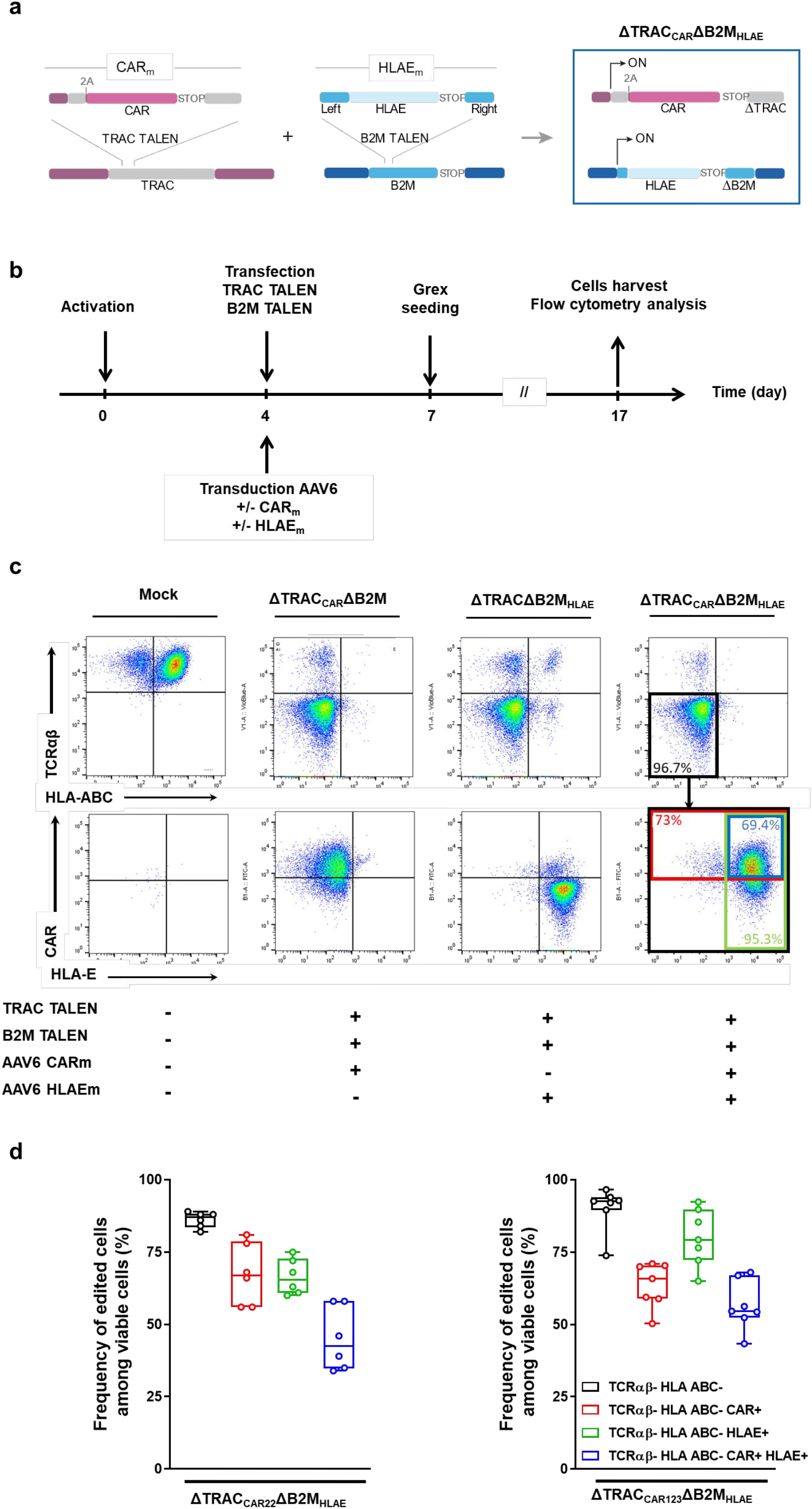
TALEN-mediated multiplex editing enables efficient CAR and HLA-E expression in TCRαβ/B2M double knock-out T-cells. **a** Schematic showing the editing strategy at the TRAC and B2M loci to generate ΔTRAC_CAR_ΔB2M_HLAE_ T-cells from wild type T-cells. **b** Experimental design for multiplex editing of TRAC and B2M loci using TALEN and adeno-associated viral (AAV6) particles and analysis of the resulting engineered T-cells. **c** Representative flow cytometry analysis of mock-transfected T-cells and T-cells engineered with different combinations of TALEN and AAV6 particles (CAR_123m_ and HLAE_m_) **c,** top panel, flow cytometry plots showing the frequency of TCRαβ and HLA-ABC expression in viable engineered T-cells. **c,** bottom panel, flow cytometry plots showing the frequency of CAR and HLA-E expression detected among TCRαβ(-)/HLA-ABC(-) viable engineered T-cells. The black bold box depicts the parent population. **d** Box plot showing the frequency of different subpopulations detected in engineered ΔTRAC_CAR22_ΔB2M_HLAE_ T-cells (left) and ΔTRAC_CAR123_ΔB2M_HLAE_ T-cells (right). The gating strategy used to obtain the frequency of each subpopulation is illustrated by the black, red, green and blue bold boxes in **1c**, right flow cytometry panel). In each box plot, the central mark indicates the median, the bottom and top edges of the box indicate the interquartile range (IQR), and the whiskers represent the maximum and minimum data point. Each point represents one experiment performed with a given donor (n = 6 for ΔTRAC_CAR22_ΔB2M_HLAE_ and n = 7 for ΔTRAC_CAR123_ΔB2M_HLAE_).

To assess the efficiency of TALEN-mediated CAR_m_ and HLAE_m_ insertion at their respective loci, we simultaneously transfected T-cells with mRNA encoding the TRAC and B2M TALEN and transduced them with CAR_m_ and HLAE_m_ using the protocol described in Fig. 1b^24^. For this proof of concept, two different CAR_m_-encoding CAR Tool constructs specific for antigens that are expressed in hematological malignancies (tool CAR_123_ and tool CAR_22_, which are specific for CD123 and CD22, respectively) were used to assess the robustness of our engineering strategy. Control samples transduced with each single repair matrix after TALEN transfection were included to accurately identify gene-edited cell subpopulations using flow cytometry. TALEN efficiently co-inactivated the TRAC and B2M genes with up to 96% of TCRαβ(-) HLA-ABC(-) T-cells obtained in the presence of both repair matrices (Fig. 1c, top panel, black box, Fig. 1d black box plots, median approx. 90%). CAR and HLA-E expression were observed in TCRαβ(-) HLA-ABC(-) T-cells (Fig. 1c, red or green boxes, respectively, Fig. 1d, red or green box plot, respectively) indicating successful disruptive insertion of both transgenes at TRAC and B2M loci. The efficiency of double targeted insertion reached as much as 68% with the two different CAR constructs (Fig. 1c blue box and 1d, blue box plots, median approx. 50%). Therefore, this TALEN/AAV6-mediated editing strategy enables efficient expression of CAR and HLA-E by TCRαβ(-) HLA-ABC(-) cells after a single transfection/transduction step. For the sake of clarity, we refer to this engineered T-cell scaffold as ΔTRAC_CAR_ΔB2M_HLAE_ in the following sections. Of note, similar levels of CAR expression and TRAC and B2M inactivations were reached using a CAR construct vectorized by recombinant lentivirus particle (rlv, Supplementary Figure 1, TLA sample 8).

### Specificity of TALEN- and AAV6-mediated engineering of ΔTRAC_CAR_ΔB2M_HLAE_

To investigate the specificity of TALEN cleavage and AAV6 matrix insertion, we performed an Oligo Capture Assay (OCA)^24^ and Targeted Locus Amplification (TLA)^25^ analysis of engineered T-cells. These methods enable unbiased identification of off-site TALEN activity and of the integration sites of CAR_m_ and HLAE_m_. We also quantified translocations between B2M and TRAC loci by qPCR. OCA analysis of TRAC and B2M TALEN co-treated T-cells identified candidate off-target sites that were then validated or invalidated using high-throughput DNA sequencing of amplicon-specific PCRs (Supplementary Figure 1a, Supplementary Table 1). High-throughput DNA sequencing of the 22 top-scoring candidate off-target sites showed insertions/deletions (indels) frequencies falling under the threshold of relevant detection (threshold = 0.16, see material and methods section), indicating that the TRAC and B2M TALEN co-treatment does not promote significant off-site targeting. As expected^2,26^, simultaneous transfection of T-cell by both TALEN, promoted translocation between the TRAC and B2M loci in up to 4% of cells (T2-TCR centromeric, Supplementary Figure 1b and Supplementary Tables 3-4). When the CARm and HLAE_m_ AAV6 matrices were added to the TALEN treatment, translocations were detected at a level similar to that observed in mock-treated T-cells. Because our qPCR method could not amplify translocation events containing the HLAE_m_ or CAR_m_ matrices, we cannot draw conclusions about the presence of translocation events integrating all or part of the AAV6 payload.

TLA analysis of engineered T-cells with the double insertion of CAR_m_ and HLAE_m_ at the TRAC and B2M loci showed that the transgenes were precisely inserted at their proper locations (CAR_m_ at chr14 and HLAE_m_ at chr15, Supplementary Figure 1d and 1f, Sample 7, red boxes), as reported earlier with similar editing strategies^24,27,28^. We also observed rare CAR_m_ and HLAE_m_ integrations at chr15 and ch14, respectively (CAR_m_ at chr15 and HLAE_m_ at chr14, Supplementary Figure 1d and 1f, Sample 7, blue box). This suggests that a small number of integrations arose from homology-independent insertion of AAV6 matrices, as documented in former reports^24,27,28^. Notably, such homology-independent insertion events were found to occur at markedly lower rates than those observed in control experiments performed with a single AAV6 matrix and an unpaired TALEN (CAR_m_ with B2M TALEN or HLAE_m_ and TRAC TALEN, compare the blue boxes in Sample 2 or 4 from Supplementary Figure 1c and 1f to Sample 7 from supplementary Figure 1d). Similar conclusions were drawn when comparing the number of homology-independent insertions obtained using AAV6 matrices with those obtained using CAR construct control vectorized by lentivirus particles along with B2M and TRAC TALEN co-transfection (Supplementary Figure 1e, sample 8, blue boxes).

### ΔTRAC_CAR_ΔB2M_HLAE_ cells display antitumor activity in vivo and in vitro

To investigate the impact of B2M depletion and HLA-E expression on CAR T-cell antitumor activity^29^, we first determined whether CAR T-cells engineered with these changes showed cytotoxic activity against leukemia cell lines *in vitro*. Engineered CAR T-cells specific for CD22 or CD123 show similar antitumor activity toward two leukemia cell line models (Fig. 2a and 2b) regardless of the number of edited features (ΔTRAC_CAR_, ΔTRAC_CAR_ΔB2M, and ΔTRAC_CAR_ΔB2M_HLAE_). We then evaluated the antitumor activity of TRAC_CAR123_ΔB2M_HLAE_ T-cells *in vivo* using MOLM13 tumors xenografted in immunodeficient NSG mice (Fig. 2c). A single administration of ΔTRAC_CAR123_ΔB2M_HLAE_ T-cells led to significantly extended survival in leukemia-bearing mice (Fig. 2d). Consistent with our *in vitro* results in the absence of host T-cells and NK cells, we observed no significant difference in overall survival between mice treated with ΔTRAC_CAR123_ T-cells and mice treated with ΔTRAC_CAR123_ΔB2M T-cells or between those treated with ΔTRAC_CAR123_ΔB2M T-cells and ΔTRAC_CAR123_ΔB2M_HLAE_ T-cells. Altogether, these *in vitro* and *in vivo* results indicate that neither the inactivation of B2M nor the disruptive targeted insertion of HLA-E at the B2M locus affect the antitumor activity of engineered CAR T-cells. This conclusion was reached with two different tool CAR constructs indicating that our engineering process could be used with other CAR constructs including those used in clinic.

**Figure 2.**
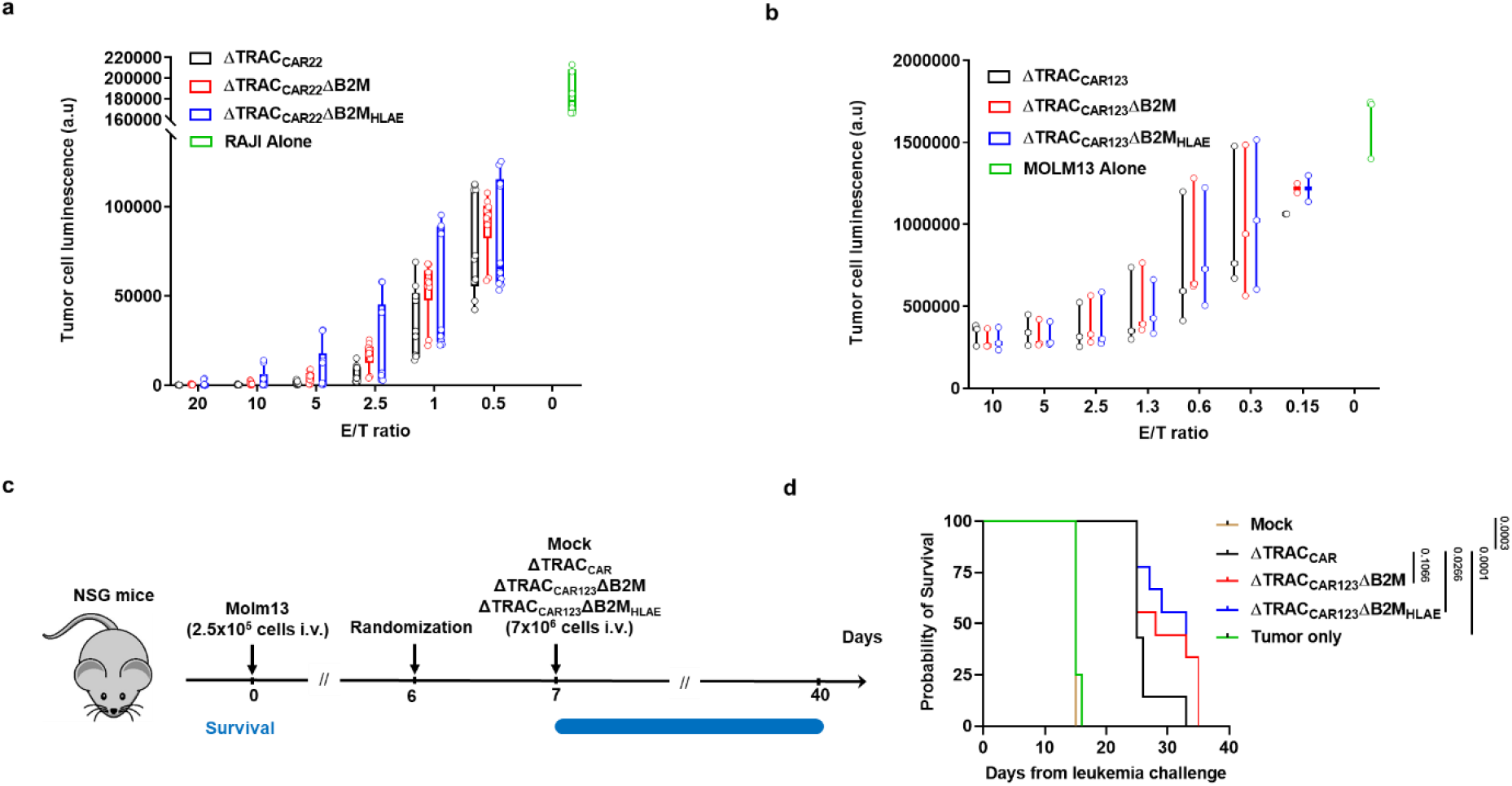
The disruptive targeted insertion of HLA-E at the B2M locus does not affect the antitumor activity of ΔTRAC_CAR_ΔB2M_HLAE_. **a, b** Antitumor activity of engineered T-cells in vitro. T-cells edited as indicated and expressing a CAR specific for either CD22 (a) or CD123 (b) were cultivated *in vitro*, with luciferase expressing RAJI and MOLM13 leukemia cell lines, respectively, using variable effector to target (E/T) ratios. The luciferase signal obtained after co-culture is plotted as a function of E/T ratio. On each box plot, the central mark indicates the median, the bottom and top edges of the box indicate the interquartile range (IQR), and the whiskers represent the maximum and minimum data point. Each point represents one experiment performed with a given T-cell donor (n=10 and n=3 for CAR T-cell specific for CD22 and CD123, respectively). **c** Schematic showing the experimental design to investigate the antitumor activity of ΔTRAC_CAR123_ΔB2M_HLAE_ in xenograft mouse model. Immunodeficient NSG mice were adoptively transferred with MOLM13-Luc-GFP leukemia cells (2.5×10^5^ cells/mouse in 100 μL of PBS via i.v.). On day 7, leukemia-bearing mice were adoptively transferred (i.v.) to randomized mice with mock-transduced T-cells (n = 7), ΔTRAC_CAR123_T-cells (n = 7), ΔTRAC_CAR123_ΔB2M T-cells (n = 9), ΔTRAC_CAR123_ΔB2M_HLAE_ T-cells (n = 9) or no T-cells (n = 8) (7×10^6^ viable CAR^+^ cells/mouse in 100 μL of PBS via i.v. injection). Mice were monitored and followed for overall survival. **d** Kaplan-Meier plot obtained for different mouse cohorts. Log-rank (Mantel-Cox) tests were used for statistical analysis. **e** Evolution of average radiance as a function of time for relevant mouse cohorts. The dotted lines correspond to the average radiance recorded for individual mice. Similar results were obtained in a second independent *in vivo* experiment (not shown). *p*-values are indicated on the figures. Source data are provided as a Source Data file.

### Targeted knock out of the B2M locus enables engineered T-cells to resist alloresponsive T-cell attacks

We next verified that the depletion of HLA-ABC from the surface of engineered T-cells prevented their elimination by alloresponsive T-cells. Alloresponsive T-cells primed for 3 weeks to recognize and attack T-cells from 3 different donors were used to perform mixed lymphocyte reactions (MLR) with TRAC and B2M TALEN-engineered T-cells (Supplementary Figure 2a). T-cells were intentionally engineered to obtain a mixture of engineered T-cell subpopulations (roughly 50% HLA-ABC(-) and 50% HLA-ABC(+)) and to assess their relative sensitivity to alloresponsive T-cell attack within each sample. The HLA-ABC(-) subpopulation of engineered T-cells (Supplementary Figure 2b) was significantly enriched by up to 23-fold over the HLA-ABC(+) subpopulation in the presence of alloresponsive T-cells in MLR assays (Supplementary Figure 2c). Consistent with former reports^c13 16^, these results indicate that the HLA-ABC(-) subpopulation is resistant to allogeneic T-cell-mediated cytolytic attack and confirms the hypoimmunogenic properties of HLA-ABC(-) cellular scaffold.

### Targeted insertion of HLA-E at the B2M locus enables engineered ΔTRAC_CAR_ΔB2M_HLAE_ to resist NK cell attacks in vitro

To assess whether HLA-E expression can prevent the depletion of HLA-ABC deficient T-cells by NK cells, we compared the sensitivity of ΔTRAC_CAR22_ΔB2M and ΔTRAC_CAR22_ΔB2M_HLAE_ cells to the cytotoxic activity of healthy donor NK cells *in vitro* (Fig. 3a). First, as a proof of principle, we intentionally engineered T-cells in suboptimal conditions to obtain balanced engineered CAR T-cell subpopulations (HLA-ABC(-)/(+) and HLA-E(-)/(+)). This allowed us to assess the relative sensitivity of these subpopulations to NK cell attack. After 3 days in culture, HLA-ABC(-) HLA-E(-) T-cells were selectively and markedly depleted by NK cells in both ΔTRAC_CAR22_ΔB2M T-cells and ΔTRAC_CAR22_ΔB2M_HLAE_ T-cells, consistent with the missing self-mediated activation of NK cells (Fig. 3b). Such depletion was correlated with a significant enrichment of the HLA-ABC(-) HLA-E(+) T-cell subpopulation of ΔTRAC_CAR22_ΔB2M_HLAE_ T-cells, as demonstrated by a 4-fold increase in the HLA-E(+) to HLA-E(-) ratio among HLA-ABC(-) T-cells compared to untreated control (Fig. 3c, left panel). This result indicates that the expression of HLA-E at the surface of T-cell successfully inhibited the cytolytic activity of NK cells. HLA-E(+)T-cells were also enriched among HLA-ABC(-) T-cells when the CD123 tool CAR was substituted for a CD22 tool CAR (about a 3-fold increase, ΔTRAC_CAR123_ΔB2M and ΔTRAC_CAR123_ΔB2M_HLAE_, Fig. 3c right panel), confirming the protective role of HLA-E against NK cell activity and demonstrating the transposable nature of this feature. It is to be noted that the values of the HLA-E(+)/HLA-E(-) ratio is more dispersed in the data set for CD123 CAR, possibly due to a larger data set (8 points vs. 4 points) where the variability in the cytolytic activity of NK cells from donor to donor is more likely to be observed.

**Figure 3.**
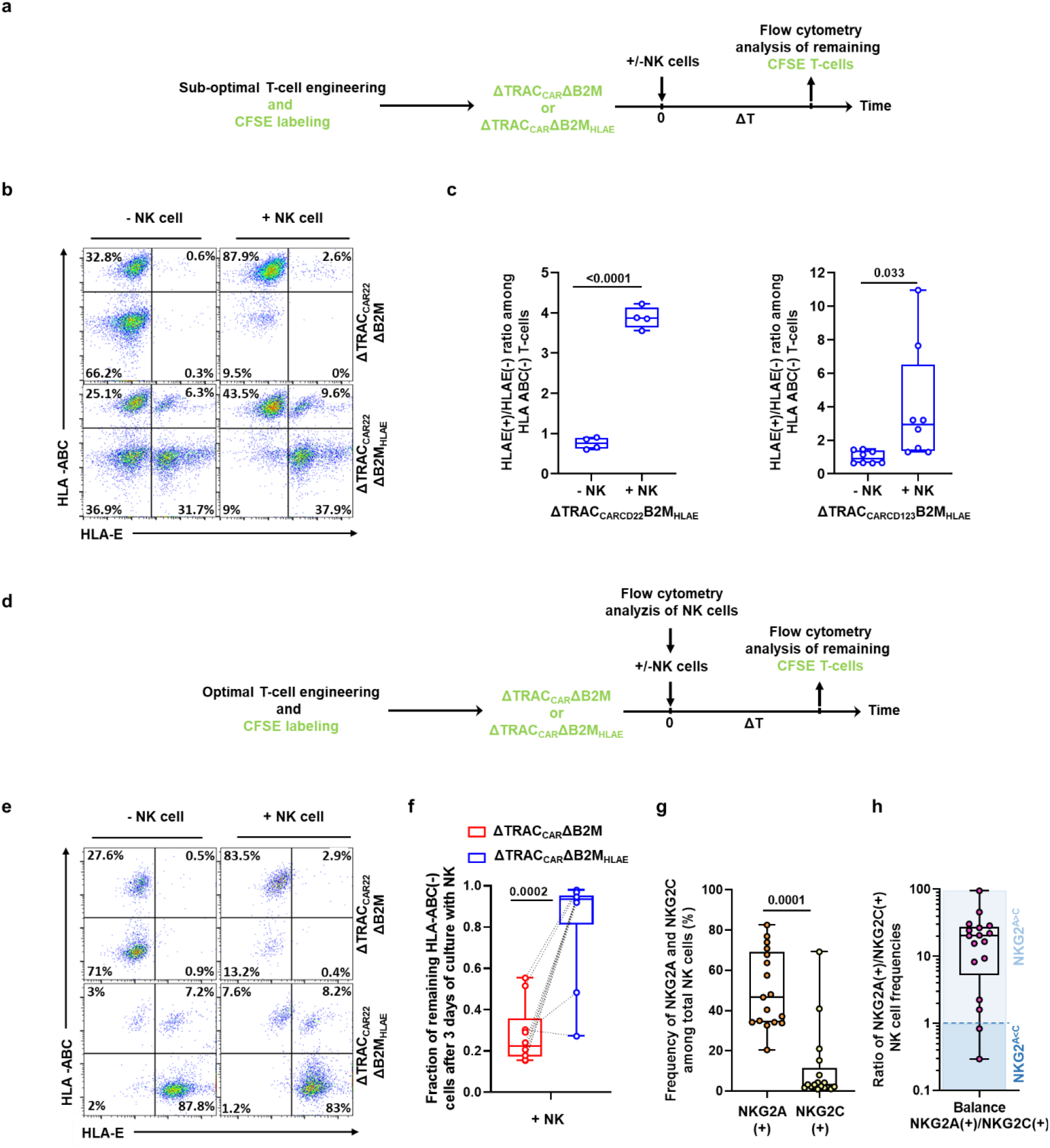
Targeted insertion of HLA-E at the B2M locus efficiently prevents NK cell-mediated depletion of ΔTRAC_CAR_ΔB2M_HLAE_ *in vitro*. **a** Schematic showing the experimental design to investigate the susceptibility of ΔTRAC_CAR_ΔB2M_HLAE_ cells, engineered under suboptimal conditions, to NK cell-dependent depletion *in vitro*. CFSE-labelled target T-cells engineered with sub-optimal condition to obtain a mixture of HLA-ABC(+)/(-) and HLA-ABC(-)/HLA-E(+)/(-) subpopulations, were cultured with or without NK cells for up to 4 days. **b** Representative flow cytometry plots showing the frequency of HLA-ABC(+)/(-) and HLA-E(+)/(-) subpopulations of ΔTRAC_CAR22_ΔB2M and ΔTRAC_CAR22_ΔB2M_HLAE_ T-cells obtained after co-culture. **c** Box plots representing the ratio of HLA-E(+) to HLA-E(-) computed from the remaining HLA-ABC(-) subpopulation of ΔTRAC_CAR_ΔB2M_HLAE_ T-cells cultured with or without NK cells; ΔTRAC_CAR22_ΔB2M_HLAE_ T-cells (left panel); ΔTRAC_CAR123_ΔB2M_HLAE_ T-cells (right panel). **d** Schematic showing the experimental design to investigate the susceptibility of ΔTRAC_CAR_ΔB2M_HLAE_ cells, engineered using optimal conditions, to NK cell-dependent depletion *in vitro*. CFSE-labelled target cells, engineered to obtain a majority of HLA-ABC(-) or HLA-ABC(-) HLA-E(+) subpopulations, were cultured in the presence or absence of NK cells for 3 days. **e** Representative flow cytometry plots showing the frequency of HLA-ABC(+)/(-) and HLA-E(+)/(-) subpopulations of ΔTRAC_CAR22_ΔB2M and ΔTRAC_CAR22_ΔB2M_HLAE_ T-cells obtained after co-culture. **f** Box plots representing the fraction of HLA-ABC(-) cells of ΔTRAC_CAR22_ΔB2M and ΔTRAC_CAR22_ΔB2M_HLAE_ T-cells remaining after co-culture (n = 5 NK donors, n = 1 T cell donor with 2 technical duplicates). **g** Box plot representing the frequencies of NKG2A(+), NKG2C(+) cells detected by flow cytometry in healthy NK cell donors (n = 17). **h** Box plot representing the ratio of NKG2A(+)/NKG2C(+) cell frequencies detected by flow cytometry in healthy donors (n = 17). NKG2^A>C^ and NKG2^C>A^ donors are shown in the light blue and dark blue fields, respectively. In each box plot, the central mark indicates the median, the bottom and top edges of the box indicate the interquartile range (IQR), and the whiskers represent the maximum and minimum data point. Paired T-test were used to compute statistics for the dataset illustrated in **c**, **f** and **g**. *p*-values are indicated on the figures. Source data are provided as a Source Data file.

To further confirm that HLA-E expression efficiently protects HLA-ABC-deficient T-cells from NK cell attack, we next engineered T-cells to obtain optimal HLA-ABC inactivation and HLA-E expression (Fig. 3d). The resulting ΔTRAC_CAR22_ΔB2M_HLAE_, harboring a majority of HLA-ABC(-)/HLA-E(+) subpopulation (∼90%) were then co-cultured with NK cells for 3 days and the remaining HLA-ABC(-) subpopulation was quantified. Consistent with the data described in Fig. 3b-c, the HLA-ABC(-) subpopulation of ΔTRAC_CAR22_ΔB2M control T-cells was significantly depleted (80% depletion compared to untreated control). The extent of this depletion was variable, suggesting that some NK cell donors may be tolerant toward HLA-ABC(-) HLA-E(-) T-cells. In stark contrast, the HLA-ABC(-) subpopulation of ΔTRAC_CAR22_ΔB2M_HLAE_ T-cells remained resistant to NK cell attack, confirming the protective role of HLA-E (Fig. 3e, 3f and supplementary Fig. 3). As expected^30,31^, ΔTRAC_CAR22_ΔB2M_HLAE_ T-cells were not killed by NK cells harvested from donors with more NKG2A(+) NK cells than NKG2C(+) NK cells (NKG2^A>C^ donors, see supplementary figure 3, donor 1, 3, 4 and 5), but from those harboring more NKG2C(+) NK cells than NKG2A(+) NK cells (NKG2^C>A^ donors, supplementary figure 3, donor 2). This trend was confirmed by investigating the level of CD107a/b degranulation of total NK cells, NKG2A(+) NK cells or NKG2C(+) NK cells subpopulations after being co-cultivated with ΔTRAC_CAR_ΔB2M_HLAE_ (Supplementary figure 4). This phenomenon is consistent with the dual specificity of HLA-E for NKG2C and NKG2A, two orthogonal surface-exposed NK receptor that are known to activate and inhibit NK cytotoxic activity, respectively, upon HLA-E engagement^32,33^. Noteworthy, because NKG2^A>C^ donors (n=15 out 17, 88%, Fig. 3g-h) were significantly more prevalent than NKG2^C>A^ donors (n=2 out 17, 12%, Fig. 3g-h), suggesting that ΔTRAC_CAR_ΔB2M_HLAE_ T-cells are widely hypoimmunogenic toward healthy donor NK cells.

### Targeted insertion of HLA-E at the B2M locus prolongs the antitumor activity engineered ΔTRAC_CAR_ΔB2M_HLAE_ in the presence of activated NK cell in vitro

We demonstrated in two independent sets of experiments that targeted insertion of HLA-E at the B2M locus did not affect the antitumor activity of ΔTRAC_CAR_ΔB2M_HLAE_ and allowed them to resist to NK cell attack. We then sought to test if these two properties held true when CAR T-cell were repeatedly co-challenged by tumor and NK cells (Fig. 4). To do so, we set up a serial killing assay where the antitumor activity of the three different engineered versions of CAR T-cell (ΔTRAC_CAR22_, ΔTRAC_CAR22_ΔB2M, ΔTRAC_CAR22_ΔB2M_HLAE_) were challenged over 4 days, by daily addition of RAJI and NK cells (NKG2^A>C^ donor, Fig.4a). Two scenario of NK cell addition were investigated (+NK day 0 or +NK day 1) to mimic the different physiological conditions that may be encountered by CAR T-cell following patients’ preconditioning. Our results showed that while ΔTRAC_CAR_ΔB2M, ΔTRAC_CAR_ΔB2M_HLAE_ showed similar antitumor activity in the absence of NK cells (Fig. 4b left panel), daily addition of NK led to complete or partial abolition of ΔTRAC_CAR_ΔB2M activity (Fig. 4b middle and right panels, respectively). In stark contrast, ΔTRAC_CAR_ΔB2M_HLAE_ remained highly active and behaved similarly to ΔTRAC_CAR_. By design, the NK-dependent drop of ΔTRAC_CAR_ΔB2M activity was correlated to a marked decrease CAR T-cell counts and to an increase of RAJI cell counts (Fig. 4c and 4d). Further analysis of the cell populations remaining at the end of serial killing assay showed that the B2M(-) subpopulation of ΔTRAC_CAR_ΔB2M was selectively depleted by NK cells while remaining constant in the case of ΔTRAC_CAR_ΔB2M_HLAE_. Similar observations were made with different CAR to RAJI and CAR to NK cell ratio (Supplementary figure 5). Taken together, these results indicate that the targeted insertion of HLA-E at the B2M locus prolongs the antitumor activity of engineered ΔTRAC_CAR_ΔB2M_HLAE_ in the presence of cytotoxic levels of activated NK cell.

**Figure 4.**
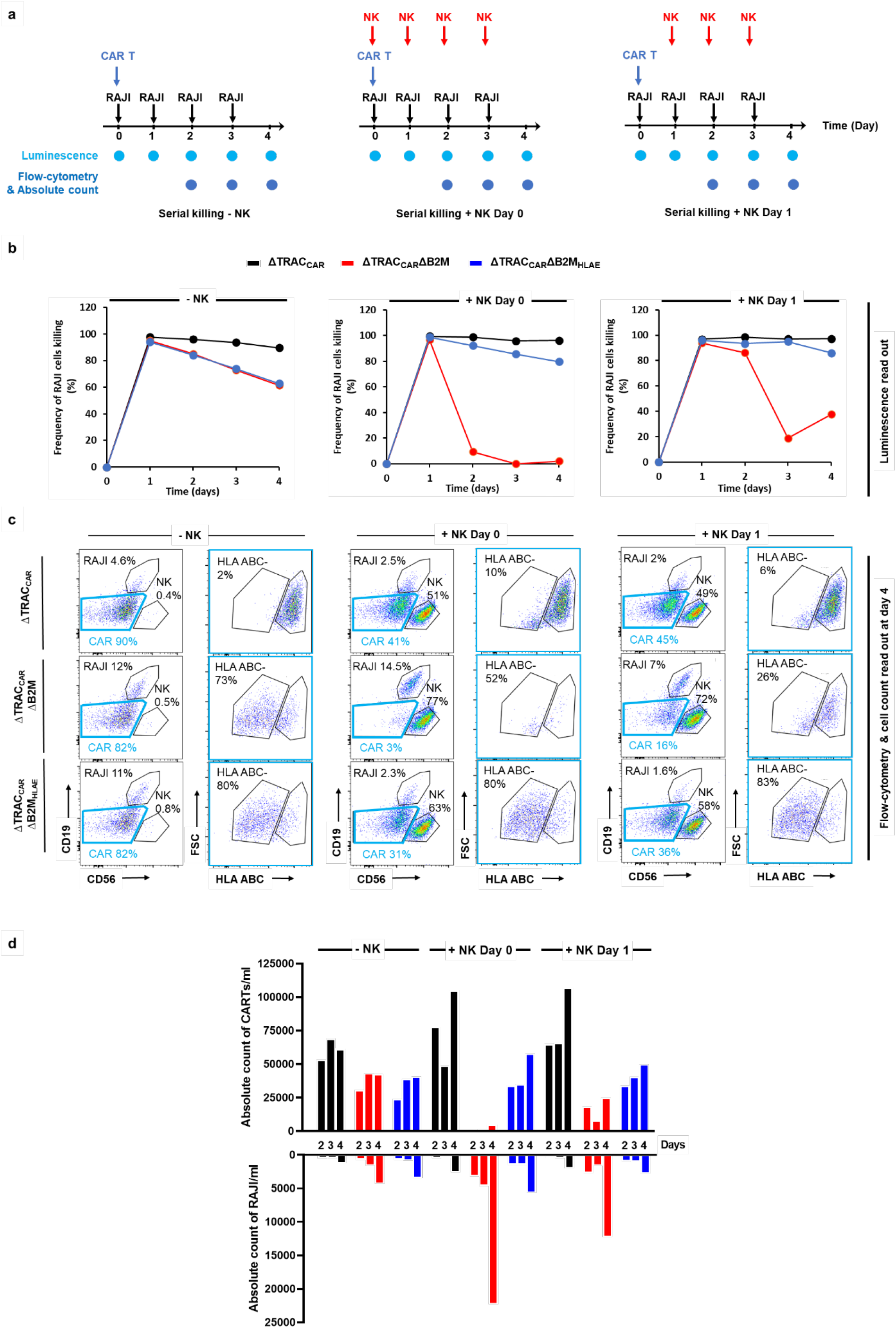
Targeted insertion of HLA-E at the B2M locus enables efficient and prolonged antitumor activity of ΔTRAC_CAR_ΔB2M_HLAE_ in the presence of activated NK cells *in vitro*. **a** Schematic showing the serial killing assay designed to investigate the long term antitumor activity of ΔTRAC_CAR22_ΔB2M_HLAE_ cells toward RAJI-Luc cells in the presence or absence of activated NK cells. The three different serial killing assay scenarios investigated are illustrated (No NK, NK Day 0 and NK Day 1). Black, blue and red arrows indicate addition of RAJI cells, CAR T-cells (ΔTRAC_CAR22_, ΔTRAC_CAR22_ΔB2M and ΔTRAC_CAR22_ΔB2M_HLAE_) and NK cells respectively. Luminescence and flow-cytometry analysis of cell populations are indicated at the different measurment time points. **b** Frequency of RAJI-Luc cells killing observed by luminescence with the three different serial killing scenarios performed with a CAR T-cell to RAJI ratio of 2.5:1 and NK to CAR T-cell ratio of 1:1. Each point represents one experiment performed with a given T-cell donor. **c** Representative flow cytometry plots showing the different cell populations remaining at the end of the serial killing assay in absence of NK cell (left panel) and in the presence of NK cells added at day 0 and at day 1 (middle and right panel, respectively). The gating strategy is indicated by blue boxes. **d** Absolute CAR T-cells and RAJI cells counts obtained by flow cytometry analysis and cell counts in the same condition as in b. Source data are provided as a Source Data file.

### NK cells from AML/ALL patients and healthy donors display similar phenotypical characteristics

Cancer patient NK cells may display different fitness and cytotoxic activity compared to healthy donor NK cells. Thus, their ability to attack and deplete the ΔTRAC_CAR_ΔB2M_HLAE_ scaffold should be evaluated to assess the robustness and potential clinical translatability of the B2M/HLAE engineering embedded in this scaffold. To gauge the magnitude of this attack, we further investigated the phenotype, fitness and cytotoxicity of clinically relevant primary NK cells obtained from several ALL and AML patients. Because NK cell numbers and fitness can be affected by previous treatment and by the nature and stage of each disease, we selected patient samples obtained before and after conventional frontline therapy. We first selected a cohort 23 ALL patients and 27 AML patients treated with induction therapy (7+3 anthracycline, cytarabine), and thoroughly investigated the phenotype of their NK cell subpopulations using mass cytometry (Fig. 5a).

**Figure 5.**
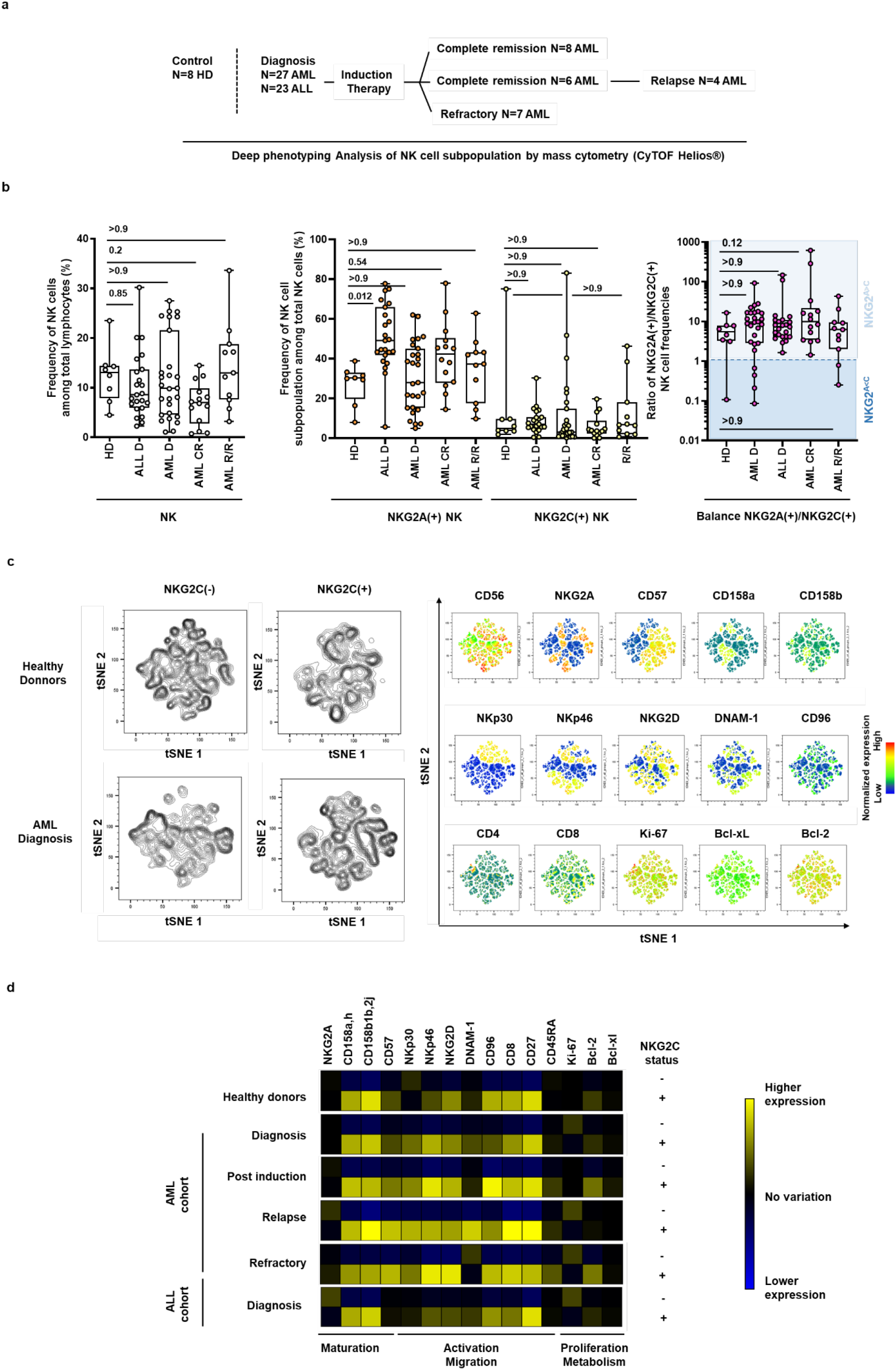
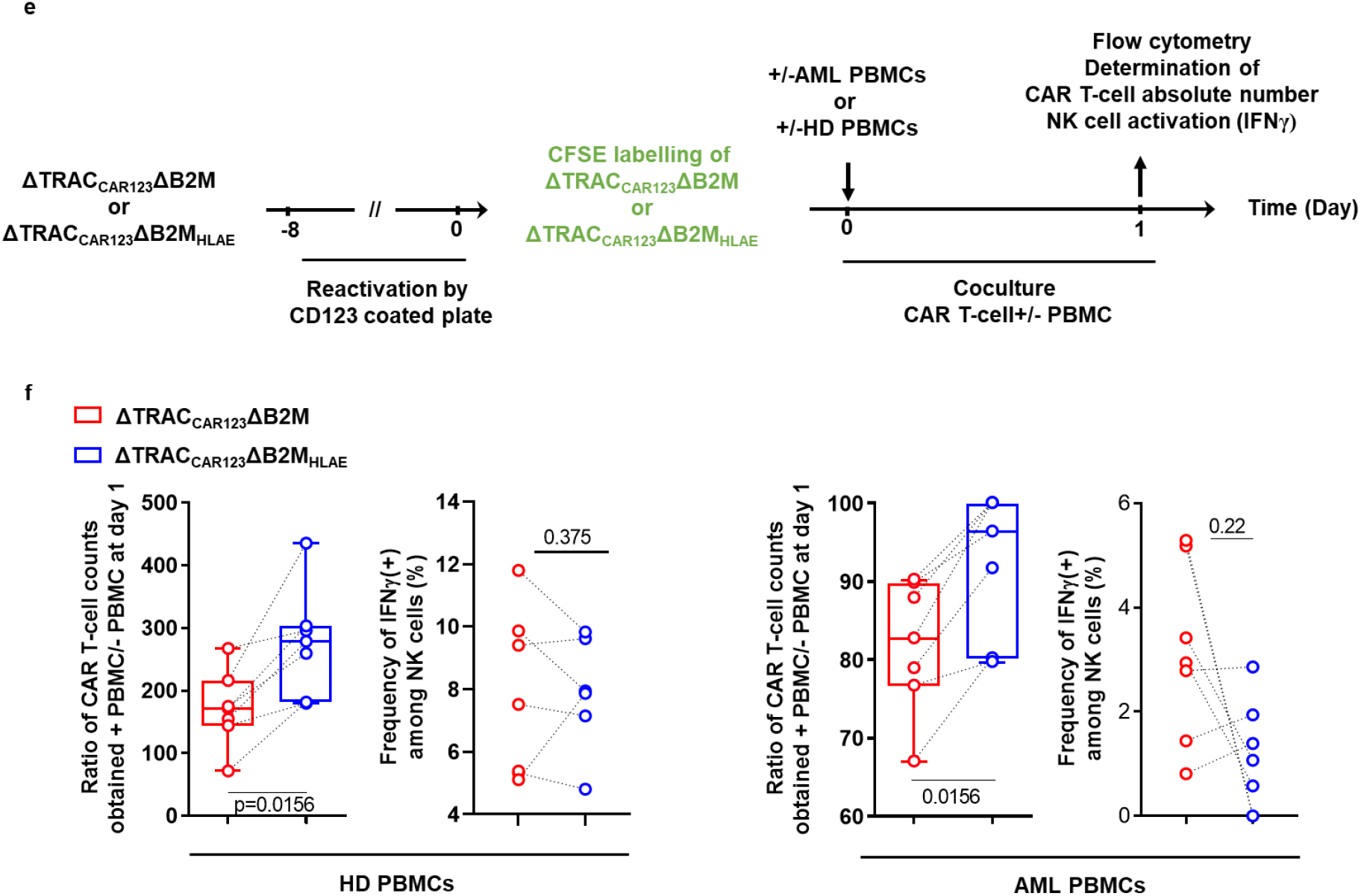
Deep phenotyping and functional characterization of NK cells from healthy donors and AML or ALL patients undergoing standard induction chemotherapy. **a** Total lymphocytes obtained from healthy donors (HD), newly-diagnosed ALL patients and AML patients selected at the time of diagnosis, complete remission, and in relapse/refractory status were characterized using mass cytometry. **b** Frequencies of NK cell subsets among total lymphocytes (left panel), frequencies of NKG2A(+) and NKG2C(+) subpopulations among total NK cell subset (middle panel), and ratio of NKG2A(+) over NKG2C(+) subpopulations (right panel) characterized by mass cytometry. For quantitative comparisons, data were analyzed using a one-way ANOVA test. **c and d** For each group described in **a**, NKG2C(-) and NKG2C(+) NK cells from each individual were exported and concatenated in order to generate consensus files. **c** The optimized parameters for T-distributed stochastic neighbor embedding (opt-SNE) algorithm were used to cluster NK cell populations based on CD56, NKG2A, CD158a,h, CD158b1,b2,j, CD57, NKp30, NKp46, NKG2D, DNAM-1, CD4 and CD8 expression. Density of NK cell clusters are shown for NKG2C(-) and NKG2C(+) NK cells in healthy donors and in AML patients at the time of diagnosis (left panel). Expression of markers of interest defining the different clusters are projected on opt-SNE maps (right panel). **d** The heatmap displays the mean frequencies of NK cell markers in NKG2C(-) and NKG2C(+) NK cells, relative to pooled NKG2C(-) and NKG2C(+) NK cells. **e**, Schematic showing the experimental design to investigate the susceptibility of ΔTRAC_CAR123_ΔB2M and ΔTRAC_CAR123_ΔB2M_HLAE_ cells, engineered using optimal conditions, to NK cell-dependent depletion *in vitro* using PBMC from AML patient at diagnostic. ΔTRAC_CAR123_ΔB2M and ΔTRAC_CAR123_ΔB2M_HLAE_ were thawed and reactivated using CD123 coated plates for 8 days. Reactivated cells were then co-cultivated for 1 day with PBMC from AML patients (n = 7) or healthy donors (n = 7) and then analyzed by flow cytometry to determine the absolute count of the remaining ΔTRAC_CAR123_ΔB2M and ΔTRAC_CAR123_ΔB2M_HLAE_. **f,** Box plots represent the ratio of ΔTRAC_CAR123_ΔB2M and ΔTRAC_CAR123_ΔB2M_HLAE_ counts obtained in the presence of PBMCs over the counts obtained in the absence of PBMCs. Dotted lines indicate data obtained from the same PBMC donor. In each box plot, the central mark indicates the median, the bottom and top edges of the box indicate the interquartile range (IQR), and the whiskers represent the maximum and minimum data point. For quantitative comparisons, data were analyzed using a non-parametric Wilcoxon matched-pairs signed rank test. *p*-values are indicated on the figures.

The frequencies (Fig. 5b, left panel) and absolute counts (Supplementary figure 6) of NK cells were comparable between the different cohorts. One exception could be however noticed for AML patients in complete remission (AML CR), who displayed a significantly lower frequency of total NK cells (Fig. 5b, left panel) and median absolute counts that fell under the normal range observed in healthy donors (Supplementary figure 6b). Frequencies of NKG2A(+) and NKG2C(+) NK cell subpopulations were also comparable between the different cohorts (except for newly diagnosed ALL patients, p value <0.0001) with a majority of patient being NKG2^A>C^ (92% of patients observed across diagnostic, complete remission and relapse/refractory status were NKG2^A>C^; Fig. 5b, middle and right panels). Therefore, these results suggest that HLA-E expression at the surface of ΔTRAC_CAR_ΔB2M_HLAE_ T-cells is likely to inhibit NK cells from most ALL and AML patients, regardless of the stage of their disease.

As described earlier for healthy donors (Fig 3g and Supplementary figure 3b), some NKG2^C>A^ outlier patients could be identified in the different cohorts studied. Indeed, these patients were identified in 4 out of 27 (15%) and 2 out of 12 (17%) of newly diagnosed and relapsed refractory AML patients, respectively (Fig. 5b, right panel). Because we observed that NK cells from NKG2^C>A^ healthy donors depleted ΔTRAC_CAR_ΔB2M_HLAE_ T-cells (Fig. 3 and Supplementary figure 3), we hypothesized that NKG2^C>A^ patient NK cells would exhibit similar behavior.

To predict the cytolytic activity of NKG2^C>A^ patient NK cells toward ΔTRAC_CAR_ΔB2M_HLAE_ T-cells, we further dissected the phenotypic characteristics of their NKG2C(+) NK cells. To do so, we investigated the fitness of NKG2C(+) NK cell subpopulations by quantitatively analyzing their maturation, activation/migration and proliferation markers (Fig. 5c-d). Deep phenotyping analysis performed on several extracellular and intracellular relevant markers, indicated that the NKG2C(+) subpopulation was similar in overall fitness and maturation in AML and ALL patients compared to healthy donors. Indeed, we observed conventional expression of CD56, NKG2A, KIRs, CD57, and of the triggering receptors NKp30, NKp46 and NKG2D (Fig. 5d). These triggering receptors were markedly more prevalent in NKG2C(+) NK cells than in NKG2C(-) NK cells, used here as a control subpopulation. The NKG2C(-) control subpopulation was altered in newly diagnosed AML patients, as evidenced by the lower expression of NKP30, NKP46 and NKG2D activating receptors compared to healthy donors (Fig.4c, left panel; compare the density plots of healthy donor and AML patients). This trend was consistent with a former report^34^, validating the robustness and reproducibility of our dataset. Finally, we observed no difference in the intrinsic fitness and maturation profile of NKG2C(+) cells in both AML and ALL cohorts after chemotherapy, suggesting that NKG2C(+) NK cells lack the classical changes described in individuals undergoing chemotherapy (Fig. 5c)^34^. Altogether, our deep phenotyping results indicate that NKG2^C>A^ outlier patients may be equipped to deplete ΔTRAC_CAR_ΔB2M_HLAE_ T-cells, although they represent a minority among the studied cohorts.

To validate our phenotypical investigation, we assessed the cytolytic activity of NK cells from AML patients against ΔTRAC_CAR123_ΔB2M_HLAE_. To do so, we co-cultivated ΔTRAC_CAR123_ΔB2M_HLAE_ or ΔTRAC_CAR123_ΔB2M with or without human peripheral blood mononuclear cells (PBMCs) from healthy donors and AML patients for 24 hours and determined their absolute number at the end of the culture (Fig. 5e). Our results showed that ΔTRAC_CAR123_ΔB2M_HLAE_ was significantly more enriched than ΔTRAC_CAR123_ΔB2M at the end of the co-culture (Fig. 5f). This HLA-E-driven enrichment was correlated with a decrease of NK cell activation probed by IFNγ release, although the number of PBMCs specimens tested was too low to reach statistical significance. These results confirmed that HLA-E expression at the surface ΔTRAC_CAR123_ΔB2M_HLAE_ could inhibit the missing-self mediated cytolytic activity of NK cells from AML patients. Taken together, these findings confirm that ΔTRAC_CAR_ΔB2M_HLAE_ T-cells are widely hypoimmunogenic toward primary NK cells isolated from multiple donors, including cancer patients.

### ΔTRAC_CAR_ΔB2M_HLAE_ T-cells resist NK cell attacks in the hIL-15 NOG mouse model

To confirm the HLA-E inhibits NK cells in a more complex model, we evaluated the persistence and enrichment of ΔTRAC_CAR123_ΔB2M_HLAE_ in hIL-15 NOG mice engrafted with PBMCs from a healthy allogeneic donor. We first verified that this *in vivo* model supported efficient engraftment and expansion of human NK cells as described earlier^35^. To do so, we intravenously injected hIL-15 NOG mice with either PBMCs or PBMCs depleted of NK cells as negative control (Fig. 6a, Supplementary Figure 7a). Flow cytometric analysis confirmed the successful engraftment of human CD45(+) immune cells and CD56(+) NK cells (Fig 5b and Supplementary Figure 7b). We then confirmed that the mouse cohort injected with PBMCs partially depleted of NK cells displayed significantly lower NK cell engraftment than did the cohort injected with PBMCs (Fig. 6b). As expected, this difference in NK levels did not impact the composition and engraftment of other immune cell compartments (Supplementary Figure 7b). In parallel, we engineered T-cells from a different donor using sub-optimal HLA-E knock-in conditions to generate ΔTRAC_CAR123_ΔB2M_HLAE_ cells with both HLA-ABC(-) HLA-E(-) and HLA-ABC(-) HLA-E(+) subpopulations (in an approximate ratio 1:1; Fig. 6c, i). This allowed us to assess the relative enrichment of HLA-E(+) over HLA-E(-) CAR T-cells in mice engrafted with allogeneic PBMCs. Four days after CAR T-cell engraftment, we observed a significant enrichment of HLA-ABC(-) HLA-E(+) ΔTRAC_CAR123_ΔB2M_HLAE_ cells in the spleen relative to their HLA-ABC(-) HLA-E(-) counterpart (Fig. 6c, d, Supplementary Figure 7d). NK cell activity in this model was corroborated by the clearance of the HLA-ABC(-) HLA-E(-) subpopulation, partially rescued in mice engrafted with NK cells-depleted PBMCs (Fig. 6c, d, Supplementary Figure 7d). Consistent with earlier results, our *in vivo* results confirmed that HLA-E expression protects HLA-ABC(-) CAR T-cells from host NK cells attack.

**Figure 6.**
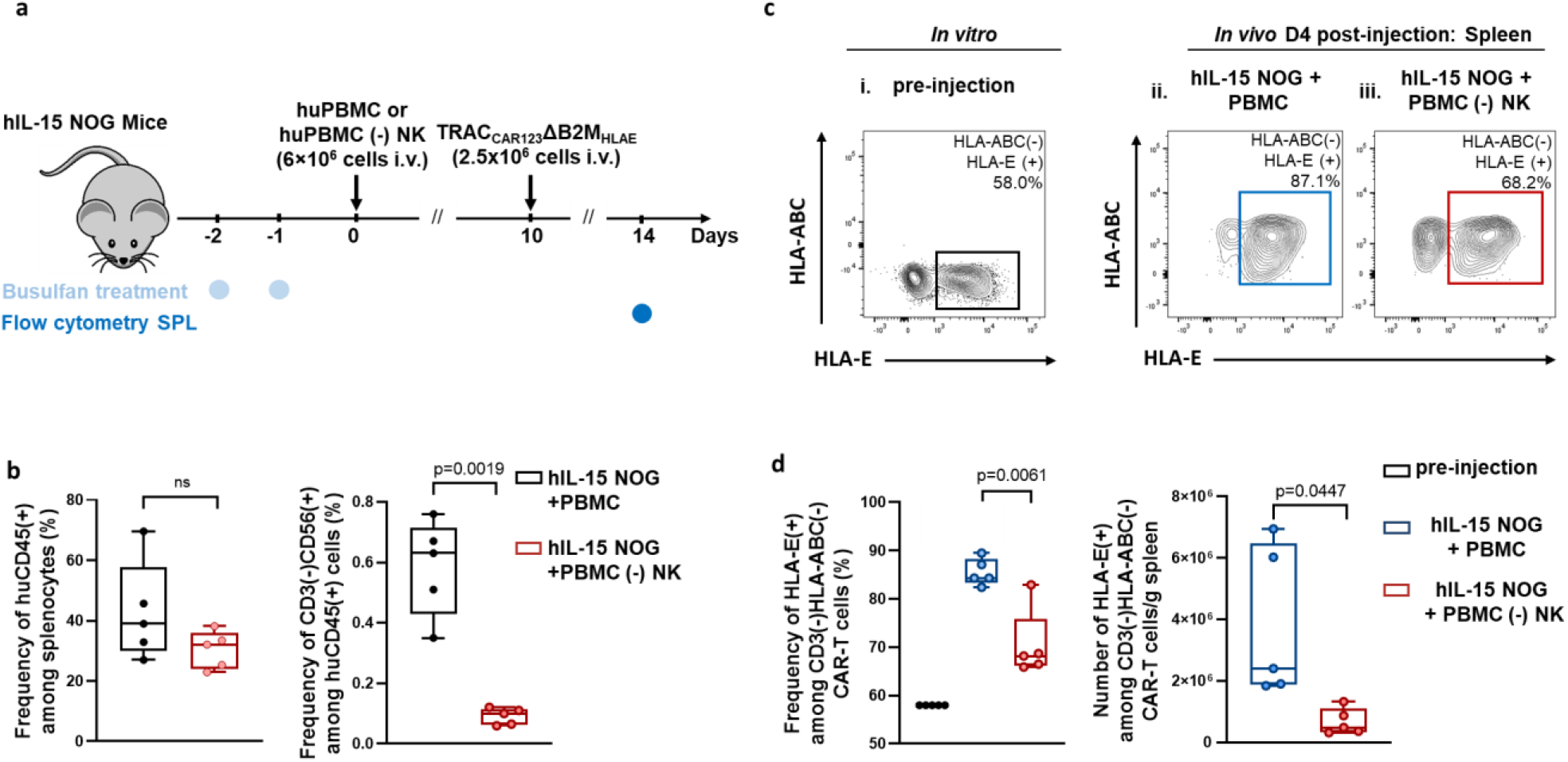
The targeted insertion of HLA-E at the B2M locus allows for efficient engraftment of ΔTRAC_CAR_ΔB2M_HLAE_ in hIL-15 NOG mice adoptively transferred with human NK cells. **a** Strategy for assessing resistance of ΔTRAC_CAR123_ΔB2M_HLAE_ T-cells to NK cells in a xenotransplantation model using hIL-15 NOG mice intravenously injected with human PBMCs. **b** Quantitation of human CD45^+^ immune cell and CD56^+^ NK cell engraftment in the spleen of hIL-15 NOG mice injected with human PBMCs or NK-depleted human PBMCs. **c** Representative flow cytometry plots showing the expression of HLA-E by ΔTRAC_CAR123_ΔB2M_HLAE_ T-cells at the time of injection and four days post intravenous injection in PBMC-engrafted hIL-15 NOG mice. **d** Box plots represent the mean value ± s.d. of **c**, depicting the percentage and absolute number of HLA-E(+) ΔTRAC_CAR123_ΔB2M_HLAE_ T-cells (n=5). On each box plot, the central mark indicates the median, the bottom and top edges of the box indicate the interquartile range (IQR), and the whiskers represent the maximum and minimum data point. Source data are provided as a Source Data file.

## Discussion

The goal of this work was to develop an immune-evasive universal CAR T-cell scaffold that is able to resist to both NK cell and alloreactive T-cell attacks and that would be compatible with adoptive cell transfer in an allogeneic setting. Using a combination of TALEN-mediated gene editing and AAV6-dependent gene insertion, we developed a hypoimmunogenic universal CAR T-cell scaffold that we designated ΔTRAC_CAR_ΔB2M_HLAE_. ΔTRAC_CAR_ΔB2M_HLAE_ is devoid of TCRαβ and HLA Class I expression and endowed with an engineered surface-exposed HLA-E. These three features enable CAR T-cells to prevent GvH reaction and to evade the cytolytic activities from alloresponsive T-cells and NK cells. While we confirmed that HLA-ABC(-) ΔTRAC_CAR_ΔB2M_HLAE_ overcame alloresponsive T-cell attack, we demonstrated their ability to evade NK cells attack by the significant enrichment of HLA-ABC(-) HLA-E(+) ΔTRAC_CAR_ΔB2M_HLAE_ in the presence of primary NK cells from healthy donors and AML patients. As a corollary, this feature enabled to prolong the antitumor activity of CAR T-cells in the presence of supra stoichiometric levels of cytotoxic NK cells *in vitro*, corroborating the very recent results obtained with hypoimmunogenic T-cells derived from iPS cells^36^. We also showed that ΔTRAC_CAR_ΔB2M_HLAE_ T-cells exhibit antitumor activity *in vivo*, similar to that of ΔTRAC_CAR_ T-cells, demonstrating that their additional hypoimmunogenic features do not affect their cytolytic functions. Finally, because NK cells from healthy donors, AML patients and ALL patients display similar phenotypic and fitness characteristics, independent of the stage of their disease, we anticipate that expression of HLA-E is likely to protect ΔTRAC_CAR_ΔB2M_HLAE_ from NK cells in the majority of AML and ALL patients.

Because of the complexity and exquisite efficacy of the human immune system, engineering hypoimmunogenic cellular scaffolds that would enable adoptive cell transfer in an allogeneic setting, has long been a daunting goal. However, the recent development of gene editing tools and a growing understanding of allograft rejection mechanisms have enabled the emergence of different engineering solutions. For instance, hypoimmunogenic cellular scaffolds have been proposed in the field of iPSC and hES transplantation^18–20,37^. One of them consists of overexpressing PD-L1, HLA-G and CD47 at the surface of pluripotent stem cells specifically depleted for HLA-A, HLA-B, HLA-C, and HLA class II ^19,20^. This immune cloaking strategy was elegantly proven to efficiently prevent NK cell-, T cell- and macrophage-dependent depletion of engineered human pluripotent stem cells. These results were confirmed by a similar study showing that the overexpression of CD47 on the surface of induced pluripotent stem cells lacking HLA class I and II surface receptors, allowed them to evade immune rejection^19^. Immune rejection of hESC was also efficiently prevented by the overexpression of PD-L1 and soluble CTLA4 immunoglobulin without relying on the inactivation of HLA class I and II^38^.

While these strategies are appealing and manageable for stem cell engineering, they require multiple iterative gene editing and antibiotic-dependent selection steps that are not well suited for the CAR T-cell manufacturing processes. In addition, the expression of certain cloaking factors including PD-L1, HLA-G, and others, may be detrimental to the cytolytic functions of CAR T-cells, due in part, to these factors’ to engage with inhibitory receptors (PD1 and ILT-2/4^39^) expressed at the surface of T-cells. Furthermore, because they are tailored to maximize the long-term engraftment of pluripotent stem cells, these strategies aim at blocking all possible allograft rejection pathways including the T-cell- (HLA Class I and II/TCR), NK cell-(missing HLA Class-I/NKG2A) and macrophage-dependent (CD47) rejection of engineered cells. This high degree of hypoimmunogenicity may not be required for efficient CAR T-cell antitumor functions because transient CAR T-cell activity correlates with positive therapeutic outcome^40–44^. Furthermore, long-term engraftment of CAR T-cells may be associated with different adverse events including the B-cell aplasia and bacterial infection usually observed in CD19 and CD22 autologous CAR T-cell therapies^45–47^. Thus, rather than engineering CAR T-cells for long term maintenance in allogenic settings, our strategy sought to extend CAR T-cell persistence by inhibiting the most important allograph rejection pathways.

Allograft rejection is mainly driven by CD8(+) T-cell, CD4(+) T-cells, NK cells and, to a lesser extent, by macrophages^48^. However, in the context of CAR T-cell therapy, the relative contribution of these cell types to allograft rejection may vary depending on their absolute numbers and reconstitution kinetics following preconditioning regimen. Indeed, clinical data obtained from three independent patient cohorts (ALL and Large B-cell malignancies) showed significant differences in CD8(+), CD4(+) T-cells and NK cell reconstitution kinetics following autologous CD19 CAR T-cell treatment and preconditioning regimen (Cyclophosphamide and fludarabine)^49–51^. Interestingly, CD8(+) T-cells and NK cells quickly recovered to their initial levels in 3 to 4 weeks, whereas CD4(+) T-cells showed a significantly slower recovery rate and, in some patients, had not returned to their initial level 1 to 2 years after treatment onset. Thus, host CD8(+) T-cells and NK cells are expected to play a key role in controlling the length of the allogeneic CAR T-cell therapeutic window by being the first and primary contributors to rejection.

Although CD8(+) T-cells need to go through a clonal expansion to mount an efficient alloresponse, they may be the first subset to reject allogeneic CAR T-cells. In this way, the inactivation of HLA Class I at the surface of CAR T-cells could efficiently blunt such rejection and offer an initial therapeutic window to eradicate cancer cells. The subsequent reconstitution of host NK cells may complexify this scenario by recognizing and depleting HLA Class I-deficient CAR T-cells without the need for clonal expansion. Embedding an NK inhibitor within CAR T-cells could further extend their persistence. We therefore focused on engineering a CAR T-cell that could evade the cytolytic activities of NK cells and allo-responsive CD8(+) T-cells. To achieve this goal, we aimed to prevent HLA Class I expression in CAR T-cells by inactivating B2M and to endow these cells with a surface-exposed NK inhibitor (HLA-E). We also considered the inactivation of TCRαβ to prevent GvHD, a major risk of allograft transplants. We reasoned that TRAC and B2M genes could be concomitantly inactivated and used as landing pads for transgene insertion (CAR and HLA-E). We forecasted that this strategy would simplify and speed up the CAR T-cell production process and mitigate potential genetic adverse events. The resulting cellular scaffold, ΔTRAC_CAR_ΔB2M_HLAE_, can be efficiently generated via a simultaneous double knock-out by knock-in strategy. The combination of TRAC and B2M TALEN treatment along with an optimized design of AAV6 repair matrices enabled us to obtain a high frequency of TCRαβ(-) HLA-ABC(-) T-cells expressing CAR and HLA-E. This engineering strategy is compatible with standard universal CAR T-cell production processes that already includes TCRαβ(+) T-cell depletion, does not require any HLA-E (+) CAR T-cell enrichment or antibiotic-dependent selection step and thus, appears suitable for clinical product manufacturing.

Co-treatment of T-cells with TRAC and B2M TALEN did not result in significant off-site cleavage activity. This was demonstrated by an unbiased OCA assay and high-throughput DNA sequencing. As expected, this co-treatment induced translocations between Chr14 and Chr15, to an extent similar to the one reported previously with different gene editing tools ^2,26,52,53^. A search of gene fusion databases, including the Atlas of Genetics and Cytogenetics in Oncology and Hematology, did not show any record of pathogenic translocation between the TRAC and B2M loci (14q11 and 15q21.1 respectively), suggesting that serious adverse events associated with such gene fusion are unlikely. However, further preclinical studies should be carried out to assess the potential toxicity of such translocations before moving to clinical studies. In addition, the homology-dependent insertion of AAV6 repair matrices was specific to the TRAC and B2M loci and correlated with robust transgene expressions. Nevertheless, we found rare homology-independent insertions of HLAE_m_ and CAR_m_ at the TRAC and B2M loci, respectively. Because both repair matrices are devoid of promoter, these improper insertions would not be expected to lead to transgene expression unless they inserted in frame with the edited gene. We did not observe CAR expression at the B2M locus (Supplementary Figure 1c, TLA sample 4), but we did find low but detectable expression of HLA-E at the TRAC locus (Supplementary Figure 1c, TLA sample 2). Although we do not envision critical adverse events related to HLA-E expression by the TRAC locus, this dataset points out one of the limitations of this multiplex gene editing strategy that must be considered in the development of other cell therapy products. It is noteworthy that this AAV6-mediated insertion approach was markedly more specific than random transgene insertion mediated by lentivirus particles (Supplementary Figure 1c, TLA sample 8).

We further report that efficient targeted insertion of HLAE_m_ at the B2M locus leads to a robust HLA-E surface expression that inhibits the cytolytic activity of NK cells. Consistent with former reports^17,23^, this outcome results from the structure and identity of the different domains of the engineered HLA-E. The inhibitory potency of HLA-E toward NK cells is intimately linked to the nature of the nonameric peptide it exposes. Its sequence was shown to significantly influence the binding affinity of HLA-E complex to the inhibitory receptor NKG2A and the activating receptor NKG2C, and thus to strongly alter the fine balance controlling NK activation and inhibition^21,22^. We chose to expose at the surface of the HLA-E engineered construct, the nonameric sequence VMAPRTLIL, because it promotes stronger engagement of NKG2A than NKG2C, as opposed to other conventional peptides including VMAPRTLFL (HLA-G-pep)^21–23^. By design, our *in vitro* results showed that our construct efficiently inhibits NKG2A(+) NK cells from multiple healthy donors and from AML patients. However, despite our choice of peptide, HLA-E can still activate NKG2C(+) NK cells, leading to the swift depletion of ΔTRAC_CAR_ΔB2M_HLAE_ T-cells in NKG2^C>A^ donors. For the reasons described earlier, we believe that one way to mitigate this activation would be to identify a non-canonical nonameric peptide displaying orthogonal specificity for NKG2A and substitute it for VMAPRTLIL within the engineered HLA-E construct.

Nevertheless, in a clinical perspective, our results suggest that the HLA-E-mediated protection of ΔTRAC_CAR_ΔB2M_HLAE_ is likely to occur in the vast majority of AML and ALL patients. Indeed, most of these patients are NKG2^A>C^ (like healthy donors) and thus harbor a greater frequency of NKG2A(+) NK cells than NKG2C(+) NK cells. This imbalance, observed at different stages of disease (92% of NKG2^A>C^ patients observed across diagnostic, complete remission and relapse refractory status, Fig. 5b), is expected to promote evasion of ΔTRAC_CAR_ΔB2M_HLAE_ from NK cells and thus, potentially extend their overall persistence, although this needs to be demonstrated in clinical settings. Determining the NKG2A(+)/NKG2C(+) ratio prior to patients injection would be beneficial for fully exploiting the therapeutic potential of ΔTRAC_CAR_ΔB2M_HLAE_.

Expressing HLA-E on the surface of HLA-ABC(-) CAR T-cells is not the only strategy to evade the cytolytic activity of NK cells. Indeed, one alternate approach would be to prevent expression of key T-cell surface receptors involved in NK cell activation. These receptors include MIC-A/B, SLAM family receptors CD48 and CD229, NKP46 ligand, B7H3 and CD155, different receptors known to activate NK cells through the engagement of their cognate receptors CD244, CD229, NKP46, IL20ra and KIR2DL5A, respectively ^37,54–57^. Genetic abrogation or downregulation of these receptors may dampen the activation of NK cells triggered by the absence of MHC Class I. Although further work is needed to evaluate the robustness and clinical translatability of this approach, a recent report demonstrated that overexpression of the Human Herpes Virus-8 ubiquitin ligase E5 in K562 cell lines or T-cells, could mitigate their depletion by NK cell through an hypothetical downregulation of MICA/B^58^. Another alternate strategy would be to endow engineered T-cells with cytolytic activity toward NK cells. This strategy was elegantly explored by Mo *et al*^59^ who recently showed that engineering CAR T-cells with an alloimmune defense receptor (ADR) specific for 4-1BB, enabled them to efficiently blunt the HvG reaction by actively targeting activated NK cells and alloresponsive T-cells.

The ultimate goal of a universal CAR T-cell product is to allow for efficient and specific depletion of cancer cells in allogeneic settings with minimal biological and toxicological effects. Our strategy was designed to mitigate such footprints by allowing CAR T-cells to evade the host immune system passively and locally without relying on active, systemic, and prolonged lymphodepletion. One potential advantage over the previously described strategies^2,9,59^ is to spare endogenous immune effectors and allow them to work in concert with CAR T-cells in the fight against cancer cells. Such collaboration could be especially useful in the context of solid tumor treatments, where endogenous immune effectors, including tissue resident memory cells^60,61^, tumor infiltrating lymphocytes^62^ and other cellular subsets are already equipped and poised to deplete tumor antigen- and neoantigen-expressing cells. The maintenance of functional endogenous immune effectors could be also a key advantage to improve the potency of CAR T-cell therapies by allowing combination therapies with oncolytic viruses^63^, vaccine boosting agents^64^, bispecifc engagers^65^ or other antibody-based immunotherapies^66^. We believe ΔTRAC_CAR_ΔB2M_HLAE_ could allow for multiple relevant combination therapies that will leverage the full potential of our human immune system and improve the therapeutic outcome of adoptive cell therapies in allogeneic settings.

In summary, we report here the development of an immune-evasive universal CAR T-cell scaffold that is deficient for TCRαβ and HLA Class I and endowed with a surface-exposed HLA-E NK inhibitor. These features render it compatible with adoptive cell transfer in allogeneic settings by preventing GvHD and allowing it to evade the cytolytic activities of NK cells and allo-responsive CD8(+) T-cells, the two major actors of HvG rejection. Although it must be demonstrated in clinical settings, these hypoimmunogenic properties could potentially extend the persistence of universal CAR T-cells and therefore, increase their antitumor potency in an immune competent host. Our engineering strategy is efficient and specific, transportable to different CAR constructs and adaptable to conventional CAR T-cell manufacturing processes. We believe this next generation of universal CAR T-cell has the potential to improve the therapeutic outcome of off-the-shelf T-cell therapies and to allow their utilization against multiple malignancies, with the potential to benefit of a broad range of patients.

## Contributions

AW, LP, PD, JV, conceived the study. AW, SD, SJ, BJ, JV, JPF, DL, ISC, AL, TP and AC designed and performed experiments. AG, MD and RG supervised DL and ICS. AC designed and perform the mass cytometry analysis of Healthy donor and AML/ALL patient samples. AL and TP perform functional assays with healthy donors and AML/ALL patient PBMCs. SJ and JPF performed the serial killing assays. EM provided critical biological reagents. RD coordinated selection and access to AML/ALL patient specimens. DO supervised AC, AL, TP and RD. J.V, SD, SJ, AC, AL, TP, DL, AD analyzed experiments. J.V. led the project, coordinated the research and wrote the manuscript in collaboration with all authors.

## Online Methods

### Materials

Cryopreserved human PBMCs were acquired from ALLCELLS (cat# PB006F). PBMCs were cultured in CTS OpTmizer media (obtained from Gibco, cat# A1048501), containing IL-2 (obtained from Miltenyi Biotec, cat# 130-097-748) or IL-7 and IL-15 (obtained from Miltenyi Biotec, cat#130-095-361 and #130-095-764), human serum AB (obtained from Seralab, cat# GEM-100-318), and CTS Immune Cell SR (obtained from Gibco, cat# A2596101). Human T Cell TransAct from Miltenyi Biotec (cat# 130-111-160) was used to activate T-cells. Antibody staining was carried out with antibodies summarized in Supplementary Table 7. Luminescence of tumor cell lines was assessed in vitro using NANO-Glo and oneGlo reagents (Promega, cat# N1110 and cat# E6110, respectively) and *in vivo* using XenoLight D-luciferin (obtained from PerkinElmer, cat#770504).

### Cell lines

MOLM13-nanoLuc-GFP and RAJI-Luc-GFP were engineered out of MOLM13 and RAJI cells (DSMZ, cat# ACC 554 and ATCC, cat# CCL-86, respectively) using an in house rLV encoding NanoLuc_T2A_EGFP construct and AMSbio cat# LVP323-PBS, respectively, using the manufacturers’ protocols.

### Targeted integration of CAR and HLA-E constructs

Targeted insertion of CAR and HLA-E at the TRAC and B2M loci were performed as described previously^24^ with minor variations. Briefly, PBMCs were thawed, washed, resuspended, and cultivated in CTS OpTmizer complete media (reconstituted CTS OpTmizer, 5% human AB serum, 20 ng/mL IL-2). One day later, the cells were activated with Human T Cell TransAct (25 µL of beads/10^6^ CD3 positive cells) and cultivated at a density of 10^6^ cells/mL for 3 days in CTS OpTmizer complete media at 37°C in the presence of 5% CO_2_. The cells were then split into fresh complete media and transduced/transfected the next day according to the following procedure. On the day of transduction-transfection, the cells were washed twice in Cytoporation buffer T (BTX Harvard Apparatus, Holliston, Massachusetts), and resuspended at a final concentration of 28×10^6^ cells/mL in the same solution. The cellular suspension (5×10^6^ cells) was mixed with 5 µg mRNA encoding each TRAC TALEN arm in the presence or absence of 5 µg of mRNA encoding each arm of B2M TALEN in a final volume of 180 µl. The cellular suspension was transfected in 0.4 cm^2^ cuvettes using Pulse Agile technology. The electroporation consisted of two 0.1 mS pulses at 3,000 V/cm followed by four 0.2 mS pulses at 325 V/cm. Immediately after electroporation, T-cells were transferred to a 12-well plate containing 2 mL of prewarmed CTS OpTmizer serum-free media and incubated at 37°C for 15 min. Half of this cellular suspension was concentrated in 250 µL of the same media in the presence or absence of AAV6 particles (MOI=2.5×10^5^ vg/cells) containing the CAR_m_ and/or HLAE_m_ matrices and seeded in 48-well plates. After 2 hours of culture at 30°C, 250 µL of CTS OpTmizer media supplemented by 10% human AB serum and 40 ng/ml IL-2 was added to the cell suspension, and the mix was incubated for overnight under the same culture conditions. The following day, cells were seeded at a density of 10^6^ cells/mL in complete CTS OpTmizer media and cultivated at 37°C in the presence of 5% CO2. On day 6 after thawing, IL-2 was replaced with IL-7 and IL-15 (2800 IU/ml and 44 IU/ml final concentration, respectively). On day 8, the cells were resuspended in fresh complete medium supplemented with IL-7 and IL-15 (2800 IU/ml and 44 IU/ml final concentration, respectively) and 5% CTS Immune Cell SR. The cells were seeded in GREX10 at 0.125×10^6^ cell/ml and cultivated in the same media according to the manufacturer’s guidelines.

### Identification and detection of candidate off-site targeting by Oligo Capture Assay and high-throughput DNA sequencing

Oligo capture assays (OCA)^24^ were used to assess the specificity of B2M and TRAC TALEN activity. Briefly, primary T-cells were co-transfected with mRNA encoding TRAC and B2M TALEN and dsODN and expanded for 6 days. Genomic DNA was recovered, sheared, end-repaired/A-tailed, processed and analyzed by high throughput DNA sequencing as described previously^24^. The resulting sequences were mapped onto the human genome (GRCh38) to identify potential off-site candidates. The frequency of insertion and deletion events (indels) generated at potential off-site candidates were then quantitatively assessed using high-throughput DNA sequencing of candidate off-site-specific PCR amplicons obtained from T-cells co-treated with TRAC and B2M TALEN (without dsODN) and TCRαβ (-) cells enriched by TCRαβ negative purification as described previously^9^. Quantification of indels frequency shows background noise due to the non-negligible error rate of the Illumina sequencing process. This error rate varies depending on the sequencing run and on the sequence itself. To quantitatively assess this error rate, sequencing datasets from 647 TALEN assayed against non-relevant targets (TALEN-transfected) were retrieved from our internal database, as were the corresponding controls performed without TALEN (mock-transfected). The difference in indels frequency between the TALEN-treated samples and the control samples was computed for each dataset. These differences followed an approximately Gaussian distribution with a mean (m) of 0.0095%, and a standard deviation (σ) of 0.04922%. We defined an off-site candidate as confirmed when the difference in indels frequency of the TALEN- and Mock-transfected T-cells was larger than a threshold (T) defined as T = m + 3 σ = 0.16%.

### Detection of translocations generated by simultaneous TRAC and B2M TALEN treatment in the presence or absence of CAR**_m_** and HLAE**_m_** AAV6 matrices

Translocation events between the TRAC and B2M loci were detected via quantitative PCR using the translocation-specific primers described in Supplementary Table 4. Reactions were performed using PowerUp SYBR Green Master Mix (ThermoScicence Cat # A25742) and the Bio-rad CFX qPCR instrument (Bio-Rad) on genomic DNA extracted from each experimental group. Amplification efficiencies and copy numbers were determined using reference control matrices designed to mimic the expected translocation events (Supplementary Table 3). Genomic DNA and control matrices were quantified using PicoGreen dsDNA quantitation assay (Thermo Fisher). Within these matrices, an XhoI restriction site was introduced at break points between B2M and TRAC TALEN target sites to control for potential contaminations of experimental samples from control matrices. The four potential translocations (T1-T4) were quantified in technical quadruplets, using sets of genomic DNA obtained from 2 independent T-cell donors. The two donors were either mock treated (negative control), co-treated with mRNA encoding the TRAC and B2M TALEN (positive control), or co-treated with mRNA encoding the TRAC and B2M TALEN and transduced by AAV6 encoding CAR_m_ and HLAE_m_.

### Targeted locus amplification assay

TLA analysis of engineered T cell was performed by Cergentis (Utrecht, Netherlands) as described^24,25^ using the engineered T-cell groups described in Supplementary Table 5 and Supplementary figure 1f and a primers described in Supplementary Table 6. Two or three specific primer sets were used for each engineered T-cells group. PCR products were purified, and library prepped using the Illumina Nextera flex protocol and sequenced on an Illumina sequencer. Reads were mapped using BWA-SW, version 0.7.15-r1140, settings bwasw -b 7. The NGS reads were aligned to the matrix sequences and host genome. Human genome build hg19 was used as a host reference genome sequence. Integration sites were detected based on a coverage peaks in the genome and on the identification of fusion-reads between the matrices sequence and the host genome as described in ^24^.

### *In vitro* T-cell antitumor activity assay

The antitumor activity of the engineered T-cells was assessed using cytotoxicity assays. Engineered T-cells were mixed with a suspension of 5×10^4^ MOLM13-Nano-Luc tumor cells at an effector to target ratio (E:T) ranging from 20:1 to 0.16:1 in a final volume of 0.1 mL of CTS OpTmizer media supplemented with 5% human AB serum. The mixture was incubated for 24 hours, and cells were lysed using 0.24% Triton-X solution. The luminescence of the remaining viable MOLM13-Nano-luc cells was determined after incubating the cell lysate with NANO-Glo reagent at a 1:1 volume ratio for 3 minutes.

### *In vitro* NK-cell cytotoxicity assay with NK cells purified from healthy donors

After thawing, cryopreserved purified human CD56^+^ NK cells (AllCells) (1×10^6^ cells/mL) in complete medium (NK MACS medium supplemented with 1% NK MACS supplement (Miltenyi Biotec) and 5% human AB serum) were cultured in a 24-well plate (500 μL/well) and incubated overnight at 37°C, 5% CO_2_. On day 1, IL-2 (60 ng/mL) was added to NK cells. After an additional 5 hrs of incubation at 37°C, 5% CO_2_, the NK cells were washed and placed in NK expansion medium (complete medium containing 40 ng/mL IL-2). Target cells (Mock-transduced T-cells, ΔTRAC_CAR_ T-cells, ΔTRAC_CAR_ΔB2M T-cells, and ΔTRAC_CAR_ΔB2M_HLAE_ T-cells), labeled with CFSE (1 mM) according to the manufacturer’s instructions, in NK expansion medium were co-cultured with IL-2 pretreated NK cells in a U-bottom 96-well plate (100 μL/well) at effector to target (5×10^4^ cells) ratios of 4:1 (CD123-specific CAR T-cells) or 1:1 (CD22-specific CAR T-cell) for 84 hrs (CD123-specific CAR T-cell) or 72 hrs (CD22-specific CAR T-cells) at 37°C, 5% CO_2_. Donor NK cells used in assays targeting CD22-specific CAR T-cells were pre-activated with NK Activation/Expansion Kit (Miltenyi Biotec) according to the manufacturer’s instructions. Cells were then analyzed using flow cytometry. Data were expressed as the ratio of HLA-E(+) to HLA-E(-) subpopulations among remaining HLA ABC(-) ΔTRAC_CAR_ΔB2M T-cells (i.e., the frequency of HLA-E(+) T-cells among total HLA ABC(-) T-cells/the frequency of HLA-E(-) T-cells among total HLA ABC(-) T-cells).

### *In vitro* expansion and activation of NK cells used in the serial killing assay

To perform an *in vitro* serial killing assay in the presence of NK cells (see below), a large-scale activation and expansion of NK cells was set up according to the following method. After thawing, cryopreserved purified human CD56^+^ NK cells, negatively selected (AllCells) (5×10^6^ cells) were resuspended at 1×10^6^ cells/mL in expansion NK MACS medium (NK MACS medium supplemented with 1% NK MACS supplement (Miltenyi Biotec), 5% human AB serum, IL-2 (500 IU/mL), and IL-15 (140 IU/mL)). Cells were then cultured in a 24-well plate (700 μL/well) and incubated at 37°C, 5% CO_2_ undisturbed for the first 5-6 days. At day 5 or 6, 300 μL expansion NK MACS medium was added without disturbing the cells. At day 7, a fresh expansion NK MACS medium was added to cells to dilute to a final concentration of 5×10^5^ cells/mL and cells were cultured in a 6-well plate (2.5 mL/well). Starting at day 10, a fresh expansion NK MACS medium was added every 2 days to dilute cells to a final concentration of 5×10^5^ cells/mL and cells were cultured in T75 flasks (>7 mL/flask) or T175 flasks (>30 mL/flask). Expanded cells were utilized in serial killing assays starting at day 13 or 14.

### *In vitro* serial killing assay in the presence of activated NK cells

To assess the persistence of ΔTRAC_CAR_ΔB2M_HLAE_ T-cells *in vitro*, a serial killing assay was performed. ΔTRAC_CAR22_ T cells, ΔTRAC_CAR22_ΔB2M T cells, and ΔTRAC_CAR22_ΔB2M_HLAE_ T cells were co-cultured with RAJI-luc tumor cells (1×10^5^) at CAR T to RAJI ratio = 5:1 and 2.5:1 in the presence or absence of *in vitro*-expanded and activated NK cells at a CAR T cell to NK cell ratio of 1:1 and 0.5:1 in a total volume of 500 μL of Xvivo-15 media supplemented with 5% AB serum in a 48-well plate. The cell mixture was incubated for 24 h before determining the luminescence of 25 μL of cell suspension using 25 μL of ONE-Glo reagent (Promega). The cell mixture was then spun down, the supernatant was discarded and replaced by 250 μL of fresh complete Xvivo-15 media containing 1×10^5^ RAJI-Luc cells, additional 250 μL of media was added to each well with or without NK cells depending on the experimental group. The resulting cell mixture was incubated for 24 h. This protocol was repeated for 4 days. Additional 20 μL from the cell culture was used for flow cytometry analysis at days 2, 3 and 4. Flow panel included the following anti-human antibodies: CD22EC-mFc (Lakepharma, 1/300), APC-Goat Anti-Mouse IgG (1/100, Jackson, cat#115135164), PE-TCRab (1/100, Miltenyi, cat# 130113539), PECy7-HLA-E (1/100,Biolegend, cat# 342608), VioBlue-HLA-ABC (1/100, Miltenyi, cat# 130120435), PercPCy5.5-CD56 (1/100, Biolegend, cat# 362526), FITC-CD19 (1/50, BDbioscience, cat# 555412). Samples were acquired using the NovoCyte Penteon flow cytometer (Agilent), which enables direct volumetric absolute count without the need for reference counting beads.

### Flow cytometry analysis

Cells in U-bottom 96-well plate were spun down and washed with PBS (150 μL/well) at 300 x g for 2 minutes. Prior to surface staining, cells were stained with Fixable Viability Dye eFluor 450 or eFluor 780 (eBiosciences) according to the manufacturer’s instructions. The cells were then stained with antibodies diluted in FACS buffer (4% FBS + 5mM EDTA + 0.05% azide in PBS) (20 μL/well) at room temperature for at least 15 minutes in the dark at 4°C. Cells were washed with PBS (150 μL/well) at 300 x g for 2 minutes and resuspended in fix buffer (4% paraformaldehyde in PBS) (100 μL/well). Sample collection was performed on a FACSCanto II cytometer (BD) or NovoCyte Penteon flow cytometer (Agilent) and data were analyzed using FlowJo V.10.6.1 (Treestar).

### Mouse models

NOD.Cg-*Prkdc^scid^ Il2rg^tm1Wjl^*/SzJ (NSG) mice and NOD.Cg-*Prkdc^scid^ Il2rg^tm1Sug^* Tg(CMV-IL2/IL15)1-1Jic/JicTac (hIL-15 NOG) mice were obtained from The Jackson Laboratory and Taconic Biosciences, respectively. Experiments were conducted in accordance with regulations and established guidelines and were reviewed and approved by the Cellectis Institutional Animal Care and Use Committee (IACUC) as well as by the Animal Ethical Committee at Mispro-Biotech (New York, NY).

### *In vivo* antitumor activity of ΔTRAC_CAR_ΔB2M_HLAE_

To assess the antitumor activity of ΔTRAC_CAR123_ΔB2M_HLAE_ *in vivo*, 7-week-old NSG mice were adoptively transferred at day 0 with MOLM13-Luc-GFP leukemia cell line (2.5×10^5^ cells/mouse in 100 μL of PBS via i.v. injection). At day 7, the mice randomly received a single-dose treatment of mock-transduced T-cells, ΔTRAC_CAR_T-cells, ΔTRAC_CAR_ΔB2M T-cells, ΔTRAC_CAR_ΔB2M_HLAE_ T-cells or no T-cells; experimental groups received a total of 7×10^6^ viable CAR(+) cells/mouse in 100 μL of PBS via intravenous injection. The mice were then monitored for health, weighed at least twice weekly, and followed to measure survival. Disease progression was monitored on a weekly basis by *in vivo* bioluminescence imaging starting at day 15. Mice were intraperitoneally injected using a XenoLight D-luciferin (PerkinElmer, 200 μL/mouse, 15 mg/mL stock solution) prior to data acquisition. All live imaging was performed on Spectrum-CT apparatus (Perkin Elmer) and Living Image software (Caliper). The endpoint was defined as the presence of ≥ 20% weight loss, hind limb paralysis, labored respiration, or inability to eat/drink. Average radiance was determined on dorsal and ventral settings, averaged and plotted as a function of time to depict tumor growth.

### *In vivo* enrichment of ΔTRAC_CAR_ΔB2M_HLAE_ in hIL-15 NOG mice engrafted with human PBMCs

To assess the persistence of ΔTRAC_CAR_ΔB2M_HLAE_ T-cells *in vivo*, cryopreserved human PBMCs acquired from ALLCELLS (cat # PB006F) were thawed as described above. NK cell depletion in freshly thawed PBMCs was performed by two successive rounds of CD56 positive selection using a NK Cell Isolation kit (Miltenyi Biotec, # 130-092-657) at twice the recommended reagent volumes. Immediately after thawing and NK depletion, cells were adoptively transferred to 7-weeks-old hIL-15 NOG mice (6×10^6^ cells/mouse in 100 µl PBS via tail vein injection). At day 12, mice within the two cohorts were randomly injected intravenously with ΔTRAC_CAR_ΔB2M_HLAE_ T-cells (2.5×10^6^ cells/mouse in 100 µl PBS). On day 16, spleens harvested from humanely euthanized mice were processed to single-cell suspensions. Cells were suspended in 2% FBS/PBS, passed through a 70 µm strainer and treated with RBC lysis buffer (eBioscience). Residual cells were labeled with monoclonal antibodies in 2% FBS/PBS buffer for 30 minutes at 4 degrees, washed with 2%FBS/PBS and subsequently fixed in 4% PFA. Sample collection was performed on FACSCanto II cytometer (BD Biosciences) and data were analyzed using FlowJo v10.6.1 (Treestar).

### Mass cytometry analysis of NK cells from AML, ALL and healthy donors

PBMCs were processed as previously described with slight modifications ^34^. PBMCs were thawed and washed with RPMI 1640 medium supplemented with 10% fetal calf serum (FCS) and incubated in RPMI 1640 with 2% FCS and 1/10000 Pierce® Universal Nuclease 5kU (Thermo Fisher Scientific, Waltham, MA, USA) at 37°C with 5% CO_2_ for 30 min. Cells were incubated with cisplatin 0.1 M to stain dead cells. Non-specific epitopes were blocked using 0.5 mg/mL Human Fc Block (BD Biosciences, San Jose, CA, USA). PBMCs were stained for 45 min at 4°C with a mix of extracellular antibodies (see Supplemental Table 7) and barcoded with the Cell-ID™ 20-Plex Pd *Barcoding Kit* (Fluidigm, San Francisco, CA, USA) according to the manufacturer’s recommendations. Cells were washed with Maxpar® cell staining buffer (CSM) (Fluidigm) and samples were combined and stained with metal-labeled anti-phycoerythrin secondary antibodies for 30 min at 4°C. After centrifugation, cells were washed with CSM and permeabilized with Foxp3 Staining Buffer (eBioscience, San Diego, CA, USA) for 40 min at 4°C. Intracellular nonspecific epitopes were blocked using 0.5 mg/mL Human Fc Block for 40 min at 4°C and then incubated with a mix of intracellular antibodies for 40 min at 4°C in Foxp3 Staining Buffer (Supplemental Table 7). Cells were then washed and labeled overnight with 125 nM iridium intercalator (Fluidigm) in Cytofix (BD Biosciences). Finally, cells were diluted in EQTM four element calibration beads (Fluidigm) and analyzed using a mass cytometer (Helios®, Fluidigm).

### Algorithm-based high-dimensional analysis

NK cells were manually defined as CD13-CD33-CD34-CD45+CD3-CD19-CD56+ and exported using FlowJo V10.6.2. Consensus files were generated for NKG2C(+) NK cells and NKG2C(-) NK cells from healthy volunteers and AML or ALL patients with a fixed number of NK cells. Data were arcsinh-transformed with a cofactor of 5. NK cell populations were automatically defined using the optimized parameters for T-distributed stochastic neighbor embedding (opt-SNE) algorithm (Belkina Nat Com 2019). The frequency of each NK cell subpopulation (among total lymphocytes as well as lymphocytosis (G/L) was used to determine the absolute counts of NK cell subsets using total cell number per volume unit obtained for each patient.

### Functional characterization of the cytolytic activity of AML patients, healthy donor PBMCs toward ΔTRAC_CAR_ΔB2M_HLAE_

Seven AML patients from Institut Paoli Calmettes (Marseille, France) entered this study after informed consent, obtained from all participants in accordance with the Declaration of Helsinki. Peripheral blood was collected at time of diagnosis and the mononuclear cells (<30% of blasts) were isolated by density gradient centrifugation (Lymphoprep; AbCys) and cryopreserved in RPMI 1640 supplemented with 10% heat-inactivated FCS (Eurobio) containing 10% of DMSO (Sigma-Aldrich). PBMCs from healthy donors (HD) were obtained from blood samples provided by the Etablissement Français du Sang (EFS, Marseille, France), after isolation by density gradient centrifugation (Lymphoprep; AbCys).

In vitro cytotoxicity assays were performed with PBMCs from HD donors or AML patients and engineered CAR T cells according to the following procedure. Cryopreserved anti CD123 CAR T-cells (ΔTRAC_CAR123_ΔB2M_HLAE_ or ΔTRAC_CAR123_ΔB2M_HLAE_), engineered according to the process delineated earlier, were thawed and reactivated on CD123-mFC coated plate during 4 days and then cultured in CTS OpTmizer media supplemented with 5% human AB serum and 40 ng/ml IL-2 for 4 additional days before cytotoxicity assay. Reactivated CAR T-cells were labeled with Celltrace Violet dye according to the manufacturer’s instructions. PBMCs effectors and labelled CAR T-cell targets (Effector:Target ratio 10:1) were co-incubated for 24 hours at 37°C in a final volume of 0.2 mL of CTS OpTmizer media supplemented with 5% human AB serum and 40 ng/ml IL-2. Cells were then collected, washed twice, and stained for surface markers (CD45, CD3, CD56, HLA-E), and viability dye (LIVE/DEAD fixable Near-IR). Regarding intracellular stainings, cells were fixed and permeabilized (BD Cytofix/Cytoperm™) according to the manufacturer’s instructions and stained with anti-IFN-γ antibodies. Cells were finally re-suspended in PBS and analyzed on a BD FACS LSRII. The antibodies and reagents used for cytometry are listed in Supplementary table 8.

## References

1. Kamiya, T., Wong, D., Png, Y. T. & Campana, D. A novel method to generate T-cell receptor-deficient chimeric antigen receptor T cells. Blood Adv. (2018) doi:10.1182/bloodadvances.2017012823.

2. Poirot, L. et al. Multiplex Genome-Edited T-cell Manufacturing Platform for ‘Off-the-Shelf’ Adoptive T-cell Immunotherapies. Cancer Res 75, 3853–3864 (2015).

3. Torikai, H. et al. A foundation for universal T-cell based immunotherapy: T cells engineered to express a CD19-specific chimeric-antigen-receptor and eliminate expression of endogenous TCR. Blood 119, 5697–5705 (2012).

4. Bunse, M. et al. RNAi-mediated TCR Knockdown Prevents Autoimmunity in Mice Caused by Mixed TCR Dimers Following TCR Gene Transfer. Mol. Ther. (2014) doi:10.1038/mt.2014.142.

5. Bot, A. et al. Cyclophosphamide and Fludarabine Conditioning Chemotherapy Induces a Key Homeostatic Cytokine Profile in Patients Prior to CAR T Cell Therapy. Blood (2015) doi:10.1182/blood.v126.23.4426.4426.

6. Gattinoni, L. et al. Removal of homeostatic cytokine sinks by lymphodepletion enhances the efficacy of adoptively transferred tumor-specific CD8+ T cells. J Exp Med 202, 907–912 (2005).

7. Yu, C. et al. Co-infusion of high-dose haploidentical donor cells and CD19-targeted CART cells achieves complete remission, successful donor engraftment and significant CART amplification in advanced ALL. Ther. Adv. Med. Oncol. (2020) doi:10.1177/1758835920927605.

8. Jin, X. et al. HLA-matched and HLA-haploidentical allogeneic CD19-directed chimeric antigen receptor T-cell infusions are feasible in relapsed or refractory B-cell acute lymphoblastic leukemia before hematopoietic stem cell transplantation. Leukemia 34, 909–913 (2020).

9. Valton, J. et al. A Multidrug-resistant Engineered CAR T Cell for Allogeneic Combination Immunotherapy. Mol Ther 23, 1507–1518 (2015).

10. Benjamin, R. et al. Preliminary Data on Safety, Cellular Kinetics and Anti-Leukemic Activity of UCART19, an Allogeneic Anti-CD19 CAR T-Cell Product, in a Pool of Adult and Pediatric Patients with High-Risk CD19+ Relapsed/Refractory B-Cell Acute Lymphoblastic Leukemia. Blood (2018) doi:10.1182/blood-2018-99-111356.

11. Qasim, W. et al. Preliminary Results of UCART19, an Allogeneic Anti-CD19 CAR T-Cell Product in a First-in-Human Trial (PALL) in Pediatric Patients with CD19+ Relapsed/Refractory B-Cell Acute Lymphoblastic Leukemia. Am. Soc. Hematol. Annu. Meet. (2017).

12. Yuki Kagoya, Tingxi Guo, Brian Yeung, Kayoko Saso, Mark Anczurowski, Chung-Hsi Wang, Kenji Murata, Kenji Sugata, Hiroshi Saijo, Yukiko Matsunaga, Yota Ohashi, M. O. B. and N. H. Genetic Ablation of HLA Class I, Class II, and the T Cell Receptor Enables Allogeneic T Cells to Be Used for Adoptive T Cell Therapy. Cancer Immunol Res (2020).

13. Ren, J. et al. Multiplex Genome Editing to Generate Universal CAR T Cells Resistant to PD1 Inhibition. Clin Cancer Res 23, 2255–2266 (2017).

14. Lee, J. et al. Abrogation of HLA surface expression using CRISPR/Cas9 genome editing: a step toward universal T cell therapy. Sci. Rep. (2020) doi:10.1038/s41598-020-74772-9.

15. Liu, X. et al. CRISPR-Cas9-mediated multiplex gene editing in CAR-T cells. Cell Research (2017) doi:10.1038/cr.2016.142.

16. Kagoya, Y. et al. Genetic ablation of HLA class I, class II, and the T cell receptor enables allogeneic T cells to be used for adoptive T cell therapy. Cancer Immunol. Res. (2020) doi:10.1158/2326-6066.cir-18-0508.

17. Crew, M. D., Cannon, M. J., Phanavanh, B. & Garcia-Borges, C. N. An HLA-E single chain trimer inhibits human NK cell reactivity towards porcine cells. Mol. Immunol. (2005) doi:10.1016/j.molimm.2004.11.013.

18. Zhao, W. et al. Strategies for Genetically Engineering Hypoimmunogenic Universal Pluripotent Stem Cells. iScience (2020) doi:10.1016/j.isci.2020.101162.

19. Deuse, T. et al. Hypoimmunogenic derivatives of induced pluripotent stem cells evade immune rejection in fully immunocompetent allogeneic recipients. Nat. Biotechnol. (2019) doi:10.1038/s41587-019-0016-3.

20. Han, X. et al. Generation of hypoimmunogenic human pluripotent stem cells. Proc. Natl. Acad. Sci. U. S. A. (2019) doi:10.1073/pnas.1902566116.

21. Hammer, Q. et al. Peptide-specific recognition of human cytomegalovirus strains controls adaptive natural killer cells article. Nat. Immunol. (2018) doi:10.1038/s41590-018-0082-6.

22. Rölle, A., Meyer, M., Calderazzo, S., Jäger, D. & Momburg, F. Distinct HLA-E Peptide Complexes Modify Antibody-Driven Effector Functions of Adaptive NK Cells. Cell Rep. (2018) doi:10.1016/j.celrep.2018.07.069.

23. Gornalusse, G. G. et al. HLA-E-expressing pluripotent stem cells escape allogeneic responses and lysis by NK cells. Nat. Biotechnol. (2017) doi:10.1038/nbt.3860.

24. Sachdeva, M. et al. Repurposing endogenous immune pathways to tailor and control chimeric antigen receptor T cell functionality. Nat. Commun. (2019) doi:10.1038/s41467-019-13088-3.

25. de Vree, P. J. et al. Targeted sequencing by proximity ligation for comprehensive variant detection and local haplotyping. Nat Biotechnol 32, 1019–1025 (2014).

26. Bothmer, A. et al. Detection and Modulation of DNA Translocations during Multi-Gene Genome Editing in T Cells. Cris. J. (2020) doi:10.1089/crispr.2019.0074.

27. Schober, K. et al. Orthotopic replacement of T-cell receptor alpha- and beta-chains with preservation of near-physiological T-cell function. Nat Biomed Eng (2019) doi:10.1038/s41551-019-0409-010.1038/s41551-019-0409-0 [pii].

28. Roth, T. L. et al. Reprogramming human T cell function and specificity with non-viral genome targeting. Nature 559, 405–409 (2018).

29. Pereira, B. I. et al. Senescent cells evade immune clearance via HLA-E-mediated NK and CD8+ T cell inhibition. Nat. Commun. (2019) doi:10.1038/s41467-019-10335-5.

30. Lanier, L. L., Corliss, B., Wu, J. & Phillips, J. H. Association of DAP12 with activating CD94/NKG2C NK cell receptors. Immunity (1998) doi:10.1016/S1074-7613(00)80574-9.

31. Lee, N. et al. HLA-E is a major ligand for the natural killer inhibitory receptor CD94/NKG2A. Proc. Natl. Acad. Sci. U. S. A. (1998) doi:10.1073/pnas.95.9.5199.

32. Houchins, J. P., Lanier, L. L., Niemi, E. C., Phillips, J. H. & Ryan, J. C. Natural killer cell cytolytic activity is inhibited by NKG2-A and activated by NKG2-C. J. Immunol. (1997).

33. Braud, V. M. et al. HLA-E binds to natural killer cell receptors CD94/NKG2A, B and C. Nature (1998) doi:10.1038/35869.

34. Chretien, A. S. et al. Natural killer defective maturation is associated with adverse clinical outcome in patients with acute myeloid leukemia. Front. Immunol. (2017) doi:10.3389/fimmu.2017.00573.

35. Katano, I. et al. Long-term maintenance of peripheral blood derived human NK cells in a novel human IL-15-transgenic NOG mouse. Sci. Rep. (2017) doi:10.1038/s41598-017-17442-7.

36. Wang, B. et al. Generation of hypoimmunogenic T cells from genetically engineered allogeneic human induced pluripotent stem cells. *Nat*. Biomed. Eng. 5, (2021).

37. Chimienti, R. et al. Engineering of NK activating receptor ligands enhances immune compatibility of MHC-I−/− iPSC-derived β cells for cell therapy of type 1 diabetes. Cytotherapy (2020) doi:10.1016/j.jcyt.2020.03.479.

38. Rong, Z. et al. An effective approach to prevent immune rejection of human ESC-derived allografts. Cell Stem Cell (2014) doi:10.1016/j.stem.2013.11.014.

39. Naji, A., Durrbach, A., Carosella, E. D. & Rouas-Freiss, N. Soluble HLA-G and HLA-G1 Expressing Antigen-Presenting Cells Inhibit T-Cell Alloproliferation through ILT-2/ILT-4/FasL-Mediated Pathways. Hum. Immunol. (2007) doi:10.1016/j.humimm.2006.10.017.

40. Van Der Stegen, S. J. C., Hamieh, M. & Sadelain, M. The pharmacology of second-generation chimeric antigen receptors. Nature Reviews Drug Discovery (2015) doi:10.1038/nrd4597.

41. Davila, M. L. et al. Efficacy and toxicity management of 19-28z CAR T cell therapy in B cell acute lymphoblastic leukemia. Sci Transl Med 6, 224ra25 (2014).

42. Maude, S. L. et al. Chimeric antigen receptor T cells for sustained remissions in leukemia. N Engl J Med 371, 1507–1517 (2014).

43. Milone, M. C. & Bhoj, V. G. The Pharmacology of T Cell Therapies. Molecular Therapy - Methods and Clinical Development (2018) doi:10.1016/j.omtm.2018.01.010.

44. Park, J. H. et al. Long-Term Follow-up of CD19 CAR Therapy in Acute Lymphoblastic Leukemia. N Engl J Med 378, 449–459 (2018).

45. Park, J. H., Geyer, M. B. & Brentjens, R. J. CD19-targeted CAR T-cell therapeutics for hematologic malignancies: interpreting clinical outcomes to date. Blood 127, 3312– 3320 (2016).

46. Fry, T. J. et al. CD22-CAR T Cells Induce Remissions in CD19-CAR Naïve and Resistant B-ALL. Nat. Med. (2018) doi:10.1038/nm.4441.

47. Wudhikarn, K. et al. Infection during the first year in patients treated with CD19 CAR T cells for diffuse large B cell lymphoma. Blood Cancer J. (2020) doi:10.1038/s41408-020-00346-7.

48. Lanza, R., Russell, D. W. & Nagy, A. Engineering universal cells that evade immune detection. Nature Reviews Immunology (2019) doi:10.1038/s41577-019-0200-1.

49. Strati, P. et al. Hematopoietic recovery and immune reconstitution after axicabtagene ciloleucel in patients with large B-cell lymphoma. Haematologica (2020) doi:10.3324/haematol.2020.254045.

50. Wang, Y. et al. Kinetics of Immune Reconstitution after CD19 CAR-T Cell Therapy in ALL Patients. Blood 134, 1301 (2019).

51. Wang, Y. et al. Kinetics of immune reconstitution after anti-CD19 chimeric antigen receptor T cell therapy in relapsed or refractory acute lymphoblastic leukemia patients. Int. J. Lab. Hematol. (2020) doi:10.1111/ijlh.13375.

52. Webber, B. R. et al. Highly efficient multiplex human T cell engineering without double-strand breaks using Cas9 base editors. Nat. Commun. (2019) doi:10.1038/s41467-019-13007-6.

53. Qasim, W. et al. Molecular remission of infant B-ALL after infusion of universal TALEN gene-edited CAR T cells. Sci Transl Med 9, (2017).

54. Husain, B. et al. A platform for extracellular interactome discovery identifies novel functional binding partners for the immune receptors B7-H3/CD276 and PVR/CD155. Mol. Cell. Proteomics (2019) doi:10.1074/mcp.TIR119.001433.

55. Pallmer, K. & Oxenius, A. Recognition and regulation of T cells by NK cells. Frontiers in Immunology (2016) doi:10.3389/fimmu.2016.00251.

56. McArdel, S. L., Terhorst, C. & Sharpe, A. H. Roles of CD48 in regulating immunity and tolerance. Clinical Immunology (2016) doi:10.1016/j.clim.2016.01.008.

57. Romero, X. et al. CD229 (Ly9) Lymphocyte Cell Surface Receptor Interacts Homophilically through Its N-Terminal Domain and Relocalizes to the Immunological Synapse. J. Immunol. (2005) doi:10.4049/jimmunol.174.11.7033.

58. Wang, X. et al. Engineering tolerance toward allogeneic CAR-T cells by regulation of MHC surface expression with Human Herpes Virus-8 proteins. Mol. Ther. (2020) doi:10.1016/j.ymthe.2020.10.019.

59. Mo, F. et al. Engineered off-the-shelf therapeutic T cells resist host immune rejection. Nat. Biotechnol. (2020) doi:10.1038/s41587-020-0601-5.

60. Malik, B. T. et al. Resident memory T cells in the skin mediate durable immunity to melanoma. Sci. Immunol. (2017) doi:10.1126/sciimmunol.aam6346.

61. Enamorado, M. et al. Enhanced anti-tumour immunity requires the interplay between resident and circulating memory CD8+ T cells. Nat. Commun. (2017) doi:10.1038/ncomms16073.

62. Zhang, Y. & Zhang, Z. The history and advances in cancer immunotherapy: understanding the characteristics of tumor-infiltrating immune cells and their therapeutic implications. Cellular and Molecular Immunology (2020) doi:10.1038/s41423-020-0488-6.

63. Park, A. K. et al. Effective combination immunotherapy using oncolytic viruses to deliver CAR targets to solid tumors. Sci. Transl. Med. (2020) doi:10.1126/SCITRANSLMED.AAZ1863.

64. Ma, L. et al. Enhanced CAR–T cell activity against solid tumors by vaccine boosting through the chimeric receptor. Science (80-. ). (2019) doi:10.1126/science.aav8692.

65. Suurs, F. V., Lub-de Hooge, M. N., de Vries, E. G. E. & de Groot, D. J. A. A review of bispecific antibodies and antibody constructs in oncology and clinical challenges. Pharmacology and Therapeutics (2019) doi:10.1016/j.pharmthera.2019.04.006.

66. Waldman, A. D., Fritz, J. M. & Lenardo, M. J. A guide to cancer immunotherapy: from T cell basic science to clinical practice. Nature Reviews Immunology (2020) doi:10.1038/s41577-020-0306-5.

