## Supplementary material for "Endowing Universal CAR T-cell with Immune-Evasive Properties using TALEN-Gene Editing": Supplemantary material

---

[‡] Co-first Authors

### Supplemental figures

**a**

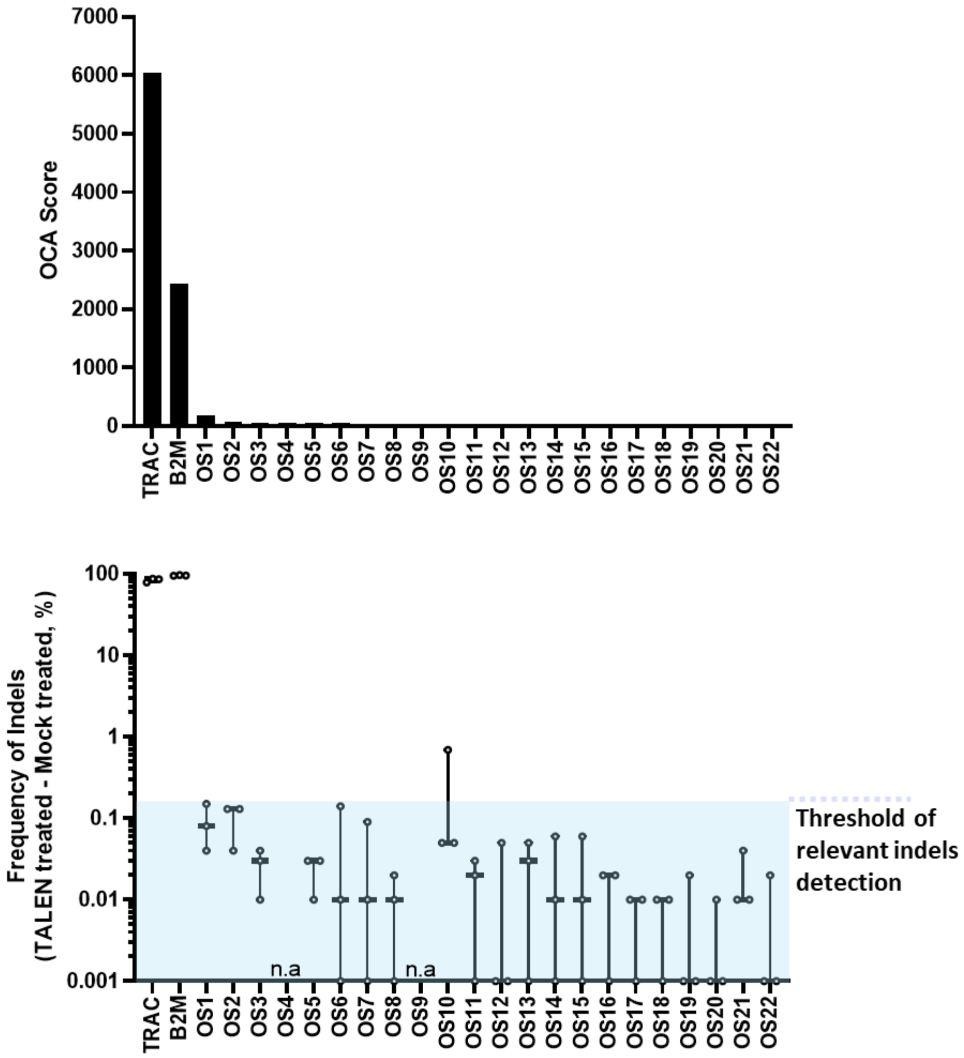

**b**

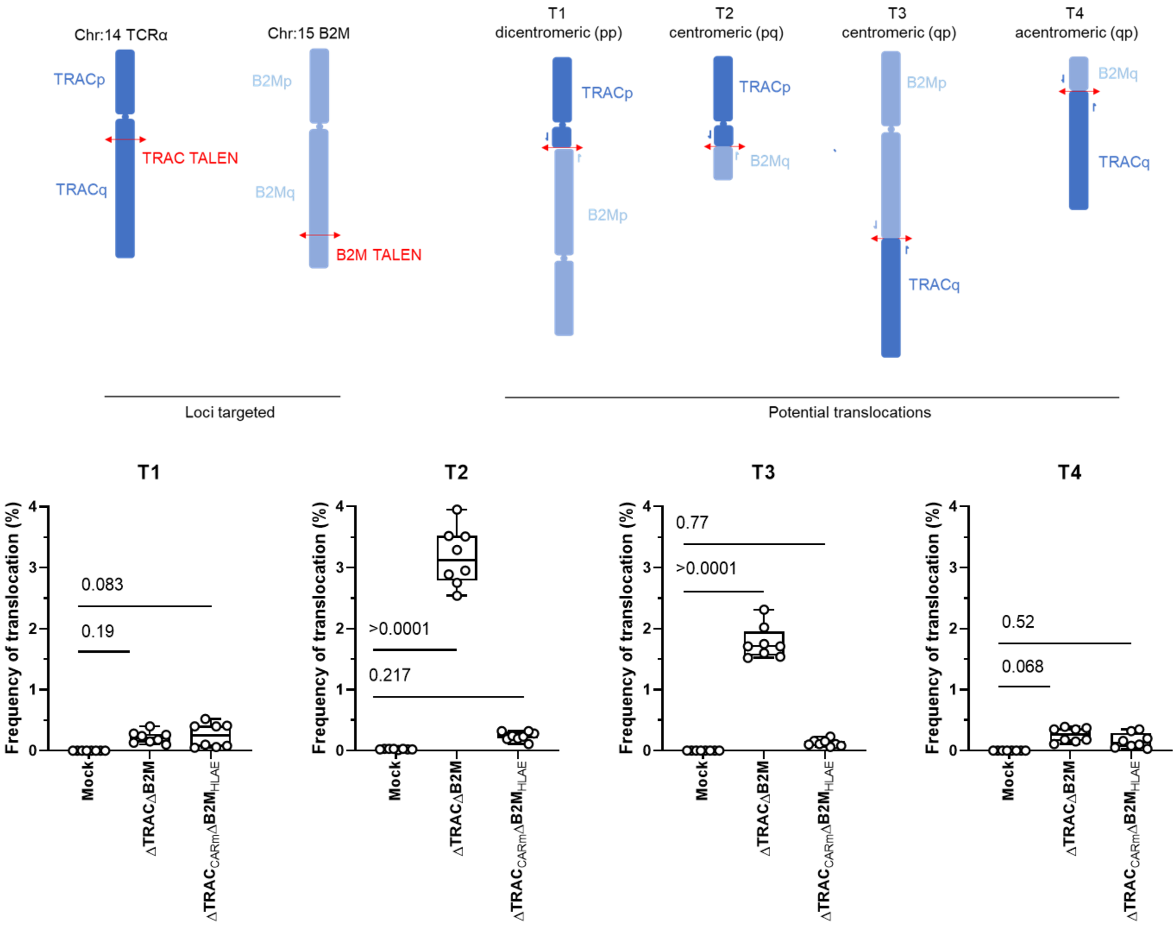
Chr:14 TCRα
TRACp
TRAC TALEN
TRACq
Chr:15 B2M
B2Mp
B2Mq
B2M TALEN
T1
dicentromeric (pp)
TRACp
B2Mp
T2
centromeric (pq)
TRACp
B2Mq
T3
centromeric (qp)
B2Mp
TRACq
T4
acentromeric (qp)
B2Mq
TRACq
Loci targeted
Potential translocations
T1
T2
T3
T4
Frequency of translocation (%)
0.083
0.19
>0.0001
0.217
0.77
>0.0001
0.52
0.068
Mock
ΔTRACΔB2M
ΔTRACCARmΔB2MHLAE

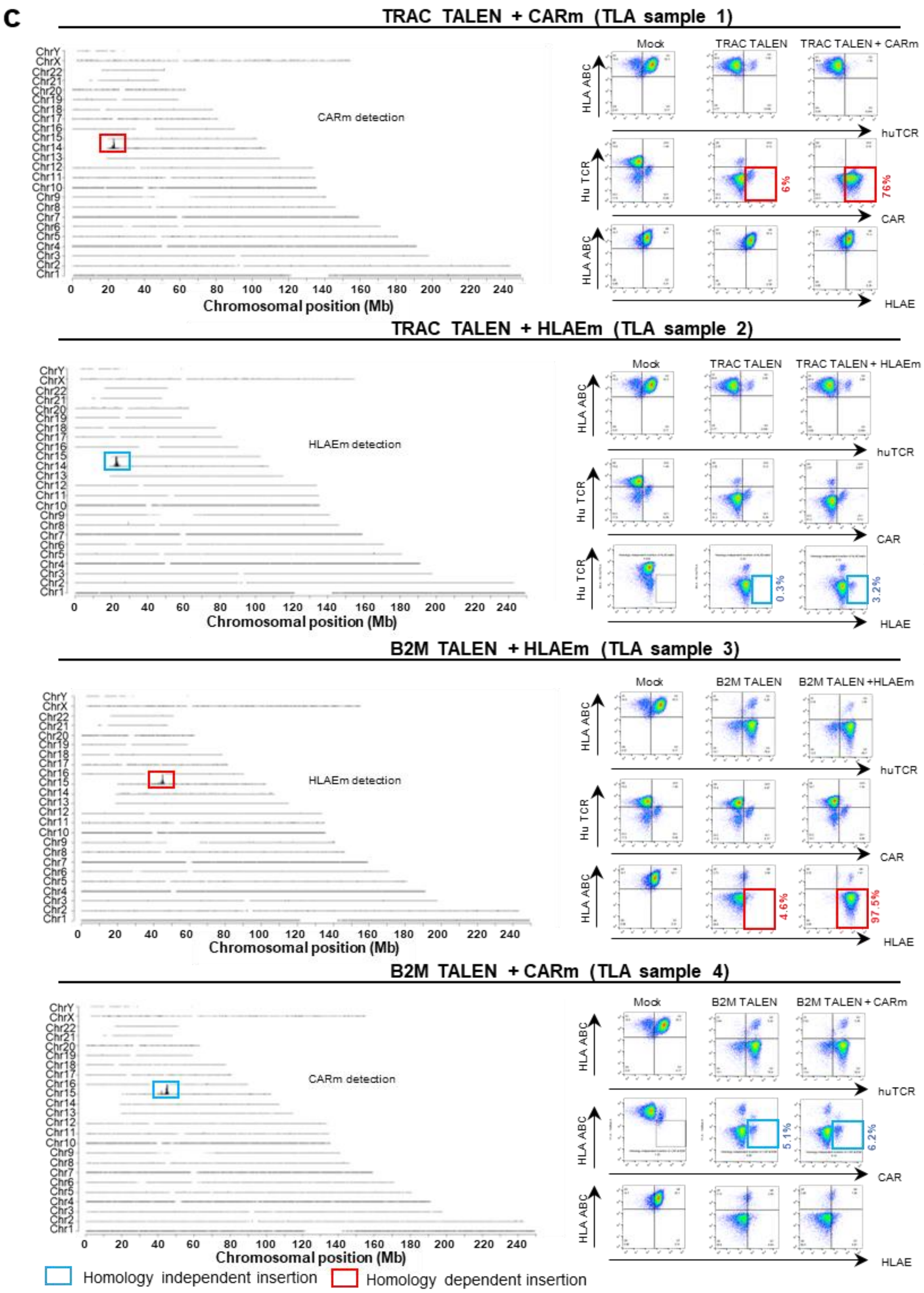
c
TRAC TALEN + CARm (TLA sample 1)
CARm detection
Chromosomal position (Mb)
Mock
TRAC TALEN
TRAC TALEN + CARm
HLA ABC
huTCR
Hu TCR
CAR
6%
76%
HLAE
TRAC TALEN + HLAEm (TLA sample 2)
HLAEm detection
TRAC TALEN + HLAEm
0.3%
3.2%
B2M TALEN + HLAEm (TLA sample 3)
B2M TALEN
B2M TALEN +HLAEm
4.6%
97.5%
B2M TALEN + CARm (TLA sample 4)
B2M TALEN + CARm
5.1%
6.2%
Homology independent insertion
Homology dependent insertion

d

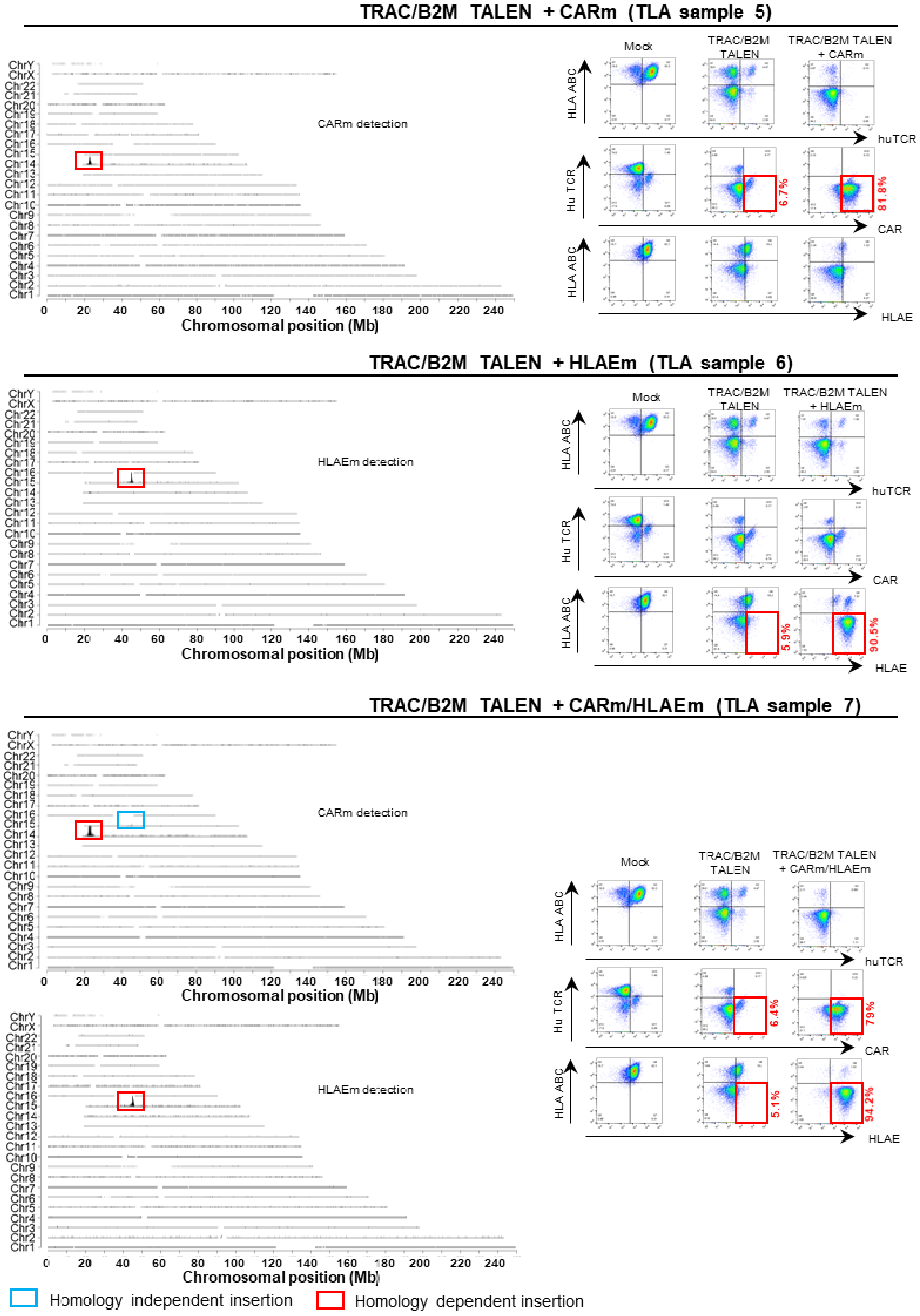
TRAC/B2M TALEN + CARm (TLA sample 5)
CARm detection
Chromosomal position (Mb)
Mock
TRAC/B2M TALEN
TRAC/B2M TALEN + CARm
HLA ABC
huTCR
Hu TCR
CAR
HLAE
6.7%
81.8%
TRAC/B2M TALEN + HLAEm (TLA sample 6)
HLAEm detection
Chromosomal position (Mb)
Mock
TRAC/B2M TALEN
TRAC/B2M TALEN + HLAEm
5.9%
90.5%
TRAC/B2M TALEN + CARm/HLAEm (TLA sample 7)
CARm detection
HLAEm detection
Chromosomal position (Mb)
Mock
TRAC/B2M TALEN
TRAC/B2M TALEN + CARm/HLAEm
6.4%
79%
5.1%
94.2%
Homology independent insertion
Homology dependent insertion

**e**

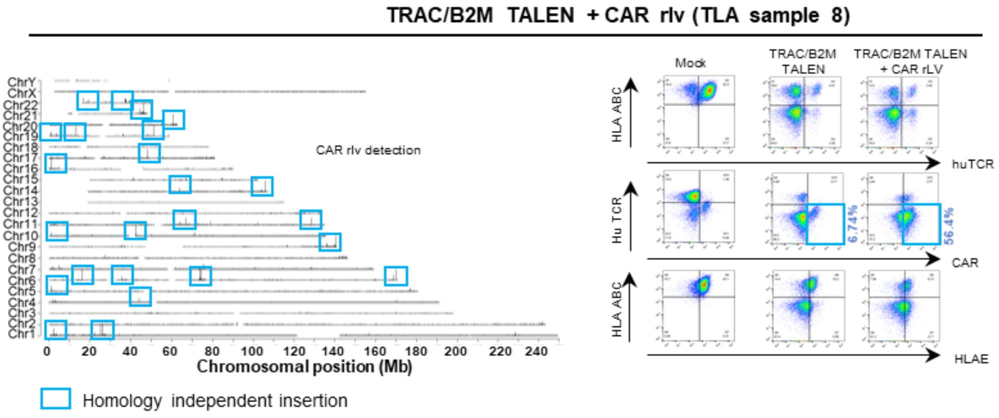


**f**

| | Experiental layout | | | |
|---|---|---|---|---|
| **Sample name** | **TALEN Modifications** | | **AAV6 matrix insertion** | |
| **Sample 1** | TRAC KO | - | CARm123 | - |
| **Sample 2** | TRAC KO | - | - | HLAEm |
| **Sample 3** | - | B2M KO | - | HLAEm |
| **Sample 4** | - | B2M KO | CARm123 | - |
| **Sample 5** | TRAC KO | B2M KO | CARm123 | - |
| **Sample 6** | TRAC KO | B2M KO | - | HLAEm |
| **Sample 7** | TRAC KO | B2M KO | CARm123 | HLAEm |
| **Sample 8** | TRAC KO | B2M KO | rlv CAR-2A-HLA-E | - |

**Supplementary figure 1. Specificity of TALEN-mediated TRAC and B2M double knock out and $CAR_m$/$HLAE_m$ targeted insertion.** **a** Top panel, Oligo Capture Assay (OCA) identifying on-site and candidate off-site TALEN activity in T-cells (n=2 donors) co-transfected with mRNA encoding TRAC and B2M TALEN. OCA score obtained for the two onsites (TRAC and B2M) as well as the 22 first candidate off-sites, are indicated. **a** Bottom panel, frequencies of indels obtained by high-throughput DNA sequencing of TRAC and B2M TALEN on-sites and of the 22 first candidate off-sites identified by OCA, except for OS4 and OS9, that could not be PCR-amplified (n.a). Frequencies of insertion and deletions (indels) obtained in TALEN treated T-cells subtracted from those obtained in mock treated T-cells are indicated (n=3 donors). The blue frame indicates the data points falling below the threshold (T) of significant indels detection (T=0.16%). **b**, qPCR analysis of canonical T1, T2, T3 and T4 translocation expected to occur between the TRAC and B2M loci targeted by their respective TALEN. **b top panel**, schemes illustrating the TRAC and B2M loci on chromosomes 14 and 15, respectively and the potential translocation generated after co-treatment of T-cells with the TRAC and B2M TALEN . **b bottom panel**, frequencies of translocation events detected by qPCR are shown for

ΔTRAC$_{CAR123}$ΔB2M$_{HLAE}$ and relevant controls T-cells (negative control, mock treated T-cell and positive control, TRAC and B2M TALEN treated T-cell, ΔTRACΔB2M) engineered out of 2 donors (n=4 technical replicates). In each box plot, the central mark indicates the median, the bottom and top edges of the box indicate the interquartile range (IQR), and the whiskers represent the maximum and minimum data point. Two-way ANOVA was used to compute statistics, and p values are indicated. **c**, **d, e left panels**, Targeted locus amplification (TLA) results. **c**, TLA results obtained from T-cells transfected with either TRAC or B2M TALEN and transduced with CAR$_m$ or HLAE$_m$ AAV6 particles. **d**, TLA results obtained from T-cells co-transfected with TRAC and B2M TALEN and transduced with CAR$_m$ or HLAE$_m$ AAV6 or co-transduced with CAR$_m$ and HLAE$_m$ AVV6 particles. **e**, TLA results obtained from T-cells co-transfected with TRAC and B2M TALEN and transduced with rlv particles encoding a CAR123-2A-HLAE construct. Homology dependent- (specific targeted integration) and homology independent- (non-specific integration) integration of AAV6 matrices are indicated in the red and blue boxes, respectively. **c**, **d, e right panels**, Flow cytometry analysis illustrating the frequency of TCR, HLA ABC, CAR and HLA-E expression within samples 1-8 analyzed by TLA and those used as relevant controls. Specific targeted expression and non-specific expression of CAR or HLA-E are indicated in the flow plots as red and blue boxes, respectively. **f,** Experimental layout to generate engineered CAR T-cells analyzed by TLA. Source data are provided as a Source Data file.

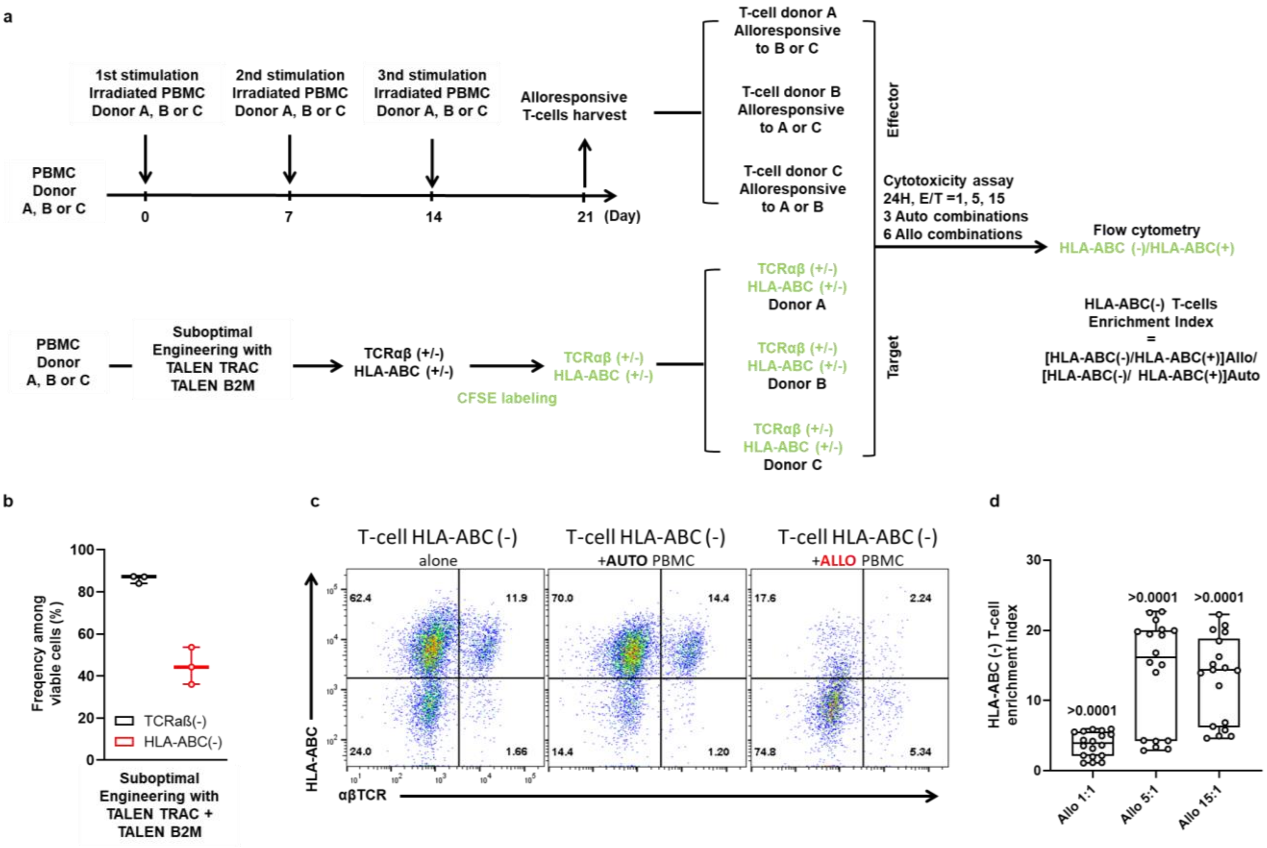


**Supplementary figure 2. Depletion of HLA-ABC from the surface of T-cell prevents their depletion by alloresponsive T-cells. a** Schematic illustrating the generation of alloresponsive T-cells and subsequent Mix lymphocyte reaction (MLR) performed with TCRαβ/B2M double knock out T-cells. PBMC from donors A, B and C were thawed and mixed at a 1:1 ratio with irradiated PBMC from donor A, B or C at day 0. Cells were co-cultivated for 7 days before being re-stimulated with irradiated PBMC from donor A, B or C at day 7. The same procedure was reiterated one more time before harvesting alloresponsive T-cells at day 21. In parallel, TCRαβ/B2M double knock out T-cells were generated from donor A, B and C (target), labeled with CFSE dye and eventually mixed with alloresponsive or autologous T-cells (effector) at a target to effector ratio of 1:1, 1:5 and 1:15. Cells were recovered after 24 hours of co-culture and analyzed by flow cytometry to determine the frequency of CFSE(+) HLA-ABC(+) and CFSE(+) HLA-ABC(-) T-cell targets remaining. An HLA-ABC(-) enrichment index was determined to quantify the resistance of CFSE(+) HLA-ABC(-) engineered T-cell to alloresponsive T-cells. **b** Efficiency of TALEN-mediated depletion of TCRαβ and HLA-ABC surface expression. Each point represents the frequency of TCRαβ and HLA-ABC knock-out obtained in a given donor. **c** representative flow cytometry plots showing the evolution of TCRαβ(+/-)HLA-ABC(+/-) T-cell targets when co-cultured in the presence of autologous PBMC (Auto PBMC) or in the presence of allogeneic PBMC (Allo PBMC) for 24 hours. **d** Box plots representing the HLA-ABC (-) enrichment index obtained when engineered TCRαβ(+/-)HLA-ABC(+/-) T-cell targets were co-cultured with autologous and alloresponsive T-cells at different ratios (3 technical replicates, with 3 donors for T-cell effector and 3 donors for engineered T-cell targets). *p*-value obtained from an unpaired t-test performed to assess the statistical significance of the HLA-ABC(-) enrichment index, are documented on the top of each box plot. On each box plot, the central mark indicates the median, the bottom and top edges of the box indicate the interquartile range (IQR) and the whiskers represent the maximum and minimum data point. Source data are provided as a Source Data file.

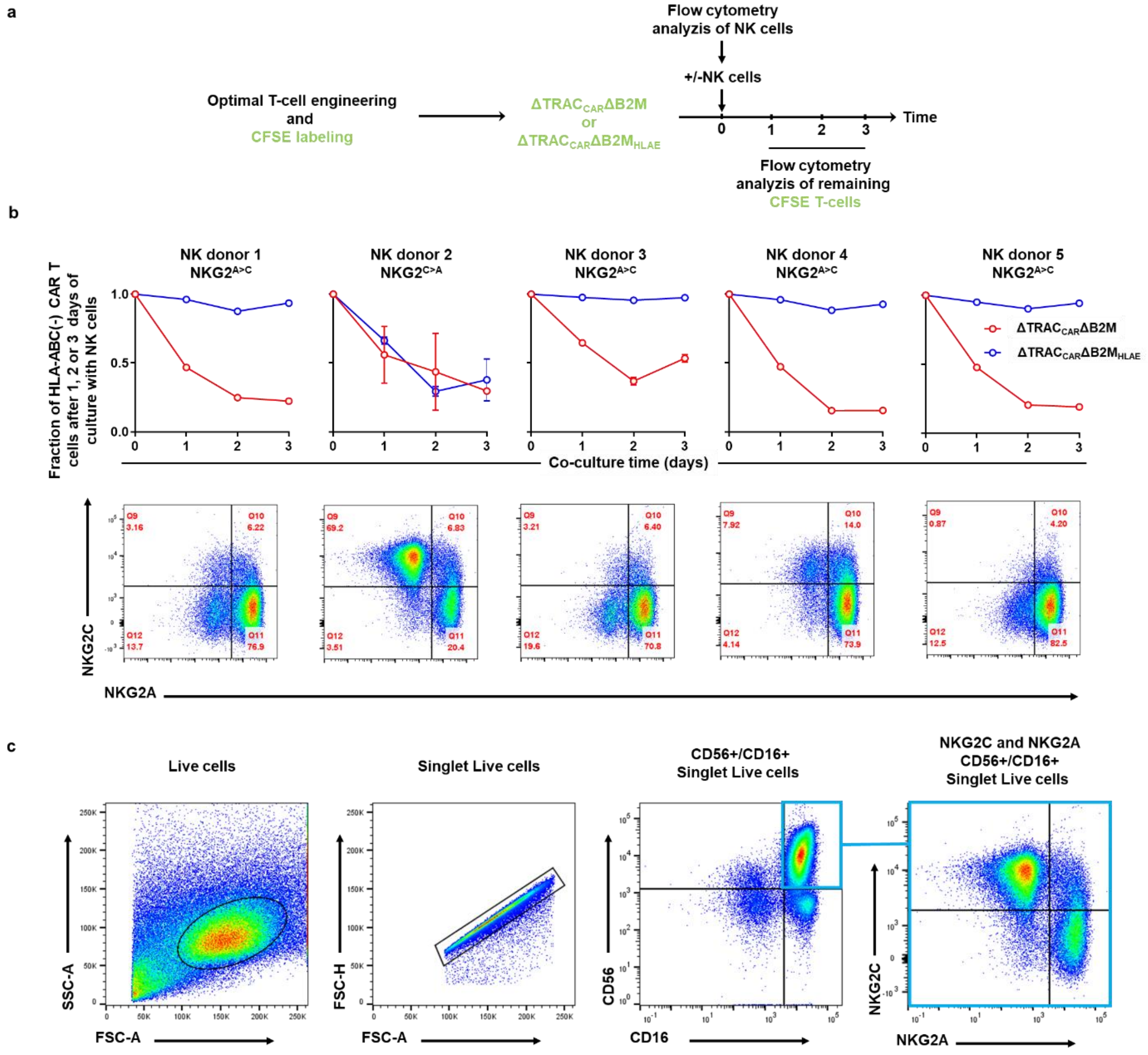

**Supplementary figure 3. Targeted insertion of HLA-E at the B2M locus efficiently prevents NK cell-mediated depletion of $\Delta TRAC_{CAR}\Delta B2M_{HLAE}$ cells *in vitro*. a** Schematic of the experimental design to investigate the susceptibility of $\Delta TRAC_{CAR}\Delta B2M_{HLAE}$ cells, engineered with optimal conditions, to NK cell-dependent depletion *in vitro*. CFSE labelled-target cells, engineered to obtain a majority of HLA-ABC (-) or HLA-ABC (-) HLA-E(+) subpopulations, were cultured for 3 days in the presence or absence of NK cells coming from 5 different donors. **b** top panel Plots showing the evolution of the fraction of HLA-ABC(-) cells of $\Delta TRAC_{CAR22}\Delta B2M$ and $\Delta TRAC_{CAR22}\Delta B2M_{HLAE}$ T-cells from 0, 1, 2 or 3 days of co-culture (n=5 NK donors, n=1 T cell donor with 2 technical duplicates). **b bottom panel**, Flow cytometry plots showing the frequency of the NKG2A(+), NKG2C(+) subpopulation among viable CD56(+), CD16(+) double-positive NK cells used to perform the NK cytotoxicity kinetics illustrated in the top panel (NK from healthy donors, n = 5). **c** Gating strategy used to obtain the flow cytometry plot of **3b**. Source data are provided as a Source Data file.

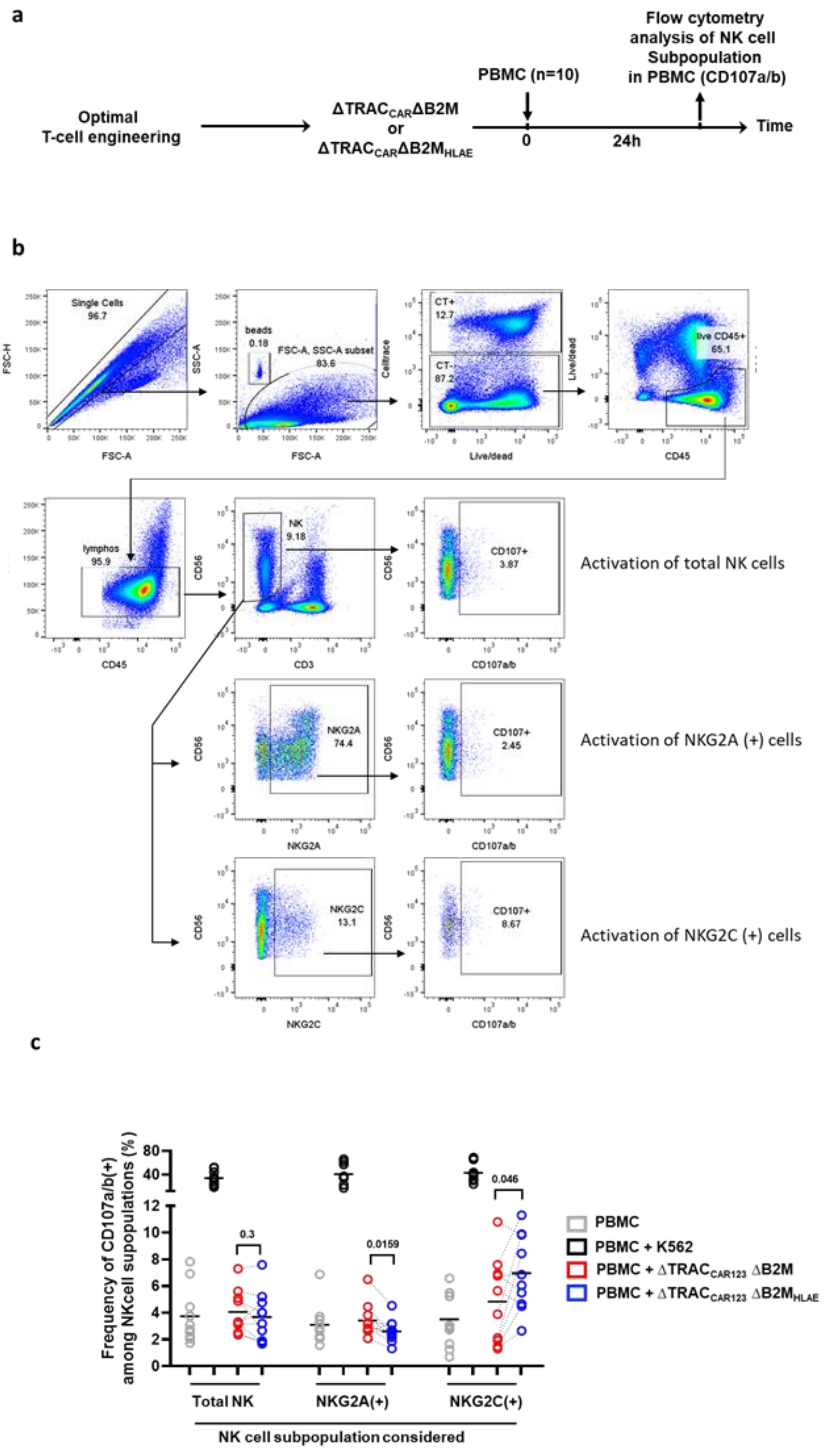
a
Optimal
T-cell engineering
ΔTRAC_CARΔB2M
or
ΔTRAC_CARΔB2M_HLAE
PBMC (n=10)
Flow cytometry
analysis of NK cell
Subpopulation
in PBMC (CD107a/b)
0
24h
Time
b
Single Cells
96.7
FSC-H
FSC-A
beads
0.18
FSC-A, SSC-A subset
83.6
SSC-A
CT+
12.7
CT-
87.2
Celltrace
Live/dead
live CD45+
65.1
CD45
lymphos
95.9
NK
9.18
CD56
CD3
CD107+
3.87
CD107a/b
Activation of total NK cells
NKG2A
74.4
CD107+
2.45
Activation of NKG2A (+) cells
NKG2C
13.1
CD107+
8.67
Activation of NKG2C (+) cells
c
Frequency of CD107a/b(+)
among NKcell supopulations (%)
0.3
0.0159
0.046
Total NK
NKG2A(+)
NKG2C(+)
NK cell subpopulation considered
PBMC
PBMC + K562
PBMC + ΔTRAC_CAR123 ΔB2M
PBMC + ΔTRAC_CAR123 ΔB2M_HLAE

**Supplementary figure 4**. **Investigation of NK cells activation/degranulation in the presence of engineered CAR T-cell ΔTRAC$_{CAR123}$ΔB2M and ΔTRAC$_{CAR123}$ΔB2M$_{HLAE}$.** 0.15$^{E}$6 ΔTRAC$_{CAR123}$ΔB2M or ΔTRAC$_{CAR123}$ΔB2M$_{HLAE}$ were mixed with 1.5$^{E}$6 PBMCs sourced from 10 different healthy donors and co-cultured for 24 hours. Cell mixture was then analyzed by flow cytometry to determine the level of activation of NK cell, NKG2A(+) and NKG2C(+) NK cell subpopulations. K562 is used as a positive control of NK activation. **a**, Scheme representing the experimental protocol. **b**, Flow cytometry gating strategy used to decipher NK cell, NKG2A(+) and NKG2C(+) NK cell populations and determine their level of CD107a/b degranulation. **c**, Summary plot illustrating the frequency of activation of NK cell, NKG2A(+) and NKG2C(+) NK cell subpopulations (CD107a/b degranulation) obtained after co-culture with K562, ΔTRAC$_{CAR123}$ΔB2M or with ΔTRAC$_{CAR123}$ΔB2M$_{HLAE}$. Each point represents one experiment performed with a given PBMC batch obtained from a healthy donor (n=10) and the black bar indicate the mean of CD107a/b frequency obtained for each experimental group. Dotted lines indicate results obtained with the same PBMC donor and ΔTRAC$_{CAR123}$ΔB2M or ΔTRAC$_{CAR123}$ΔB2M$_{HLAE}$. The *p*-value obtained from a paired t-test is indicated on the figure.

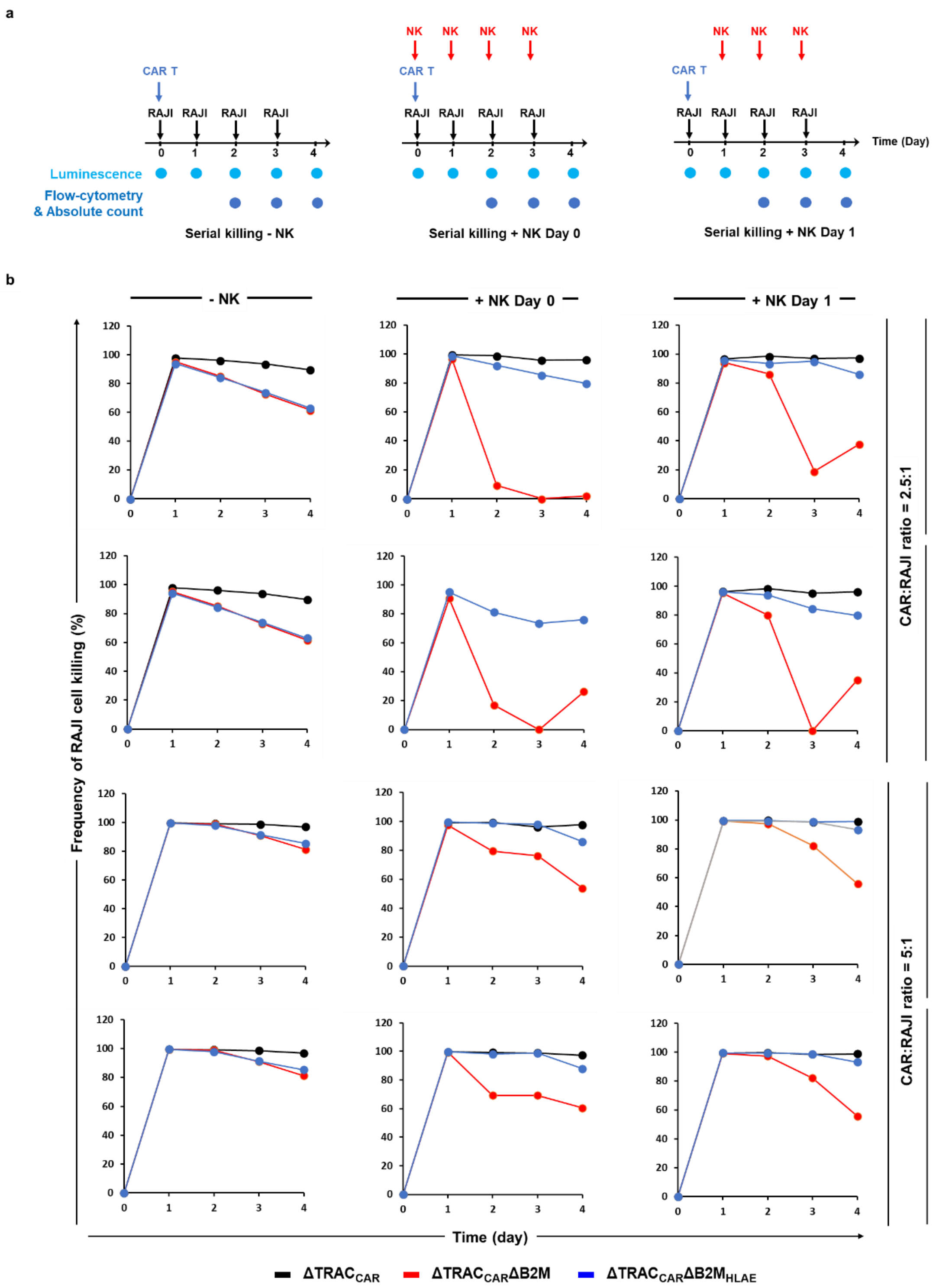

a
CAR T
RAJI RAJI RAJI RAJI
Luminescence
Flow-cytometry & Absolute count
Serial killing - NK
NK NK NK NK
Serial killing + NK Day 0
NK NK NK
Serial killing + NK Day 1
Time (Day)
b
- NK
+ NK Day 0
+ NK Day 1
Frequency of RAJI cell killing (%)
Time (day)
CAR:RAJI ratio = 2.5:1
CAR:RAJI ratio = 5:1
(CAR:NK ratio = 1:1)
(CAR:NK ratio = 1:2)
ΔTRAC_CAR ΔTRAC_CARΔB2M ΔTRAC_CARΔB2M_HLAE

c

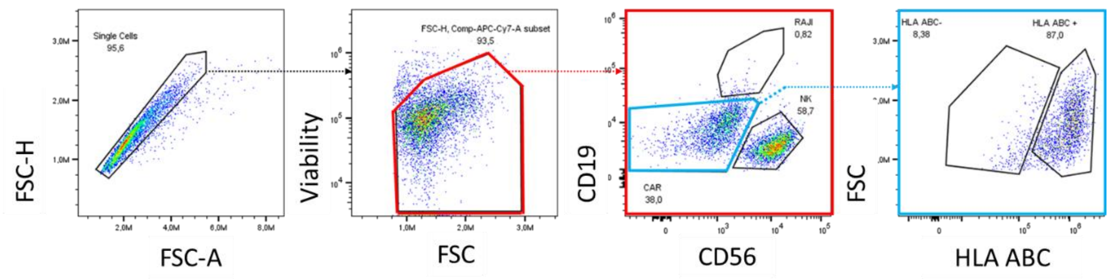
Single Cells
95,6
FSC-H
FSC-A
FSC-H, Comp-APC-Cy7-A subset
93,5
Viability
FSC
RAJI
0,82
NK
58,7
CAR
38,0
CD19
CD56
HLA ABC-
8,38
HLA ABC +
87,0
FSC
HLA ABC

d

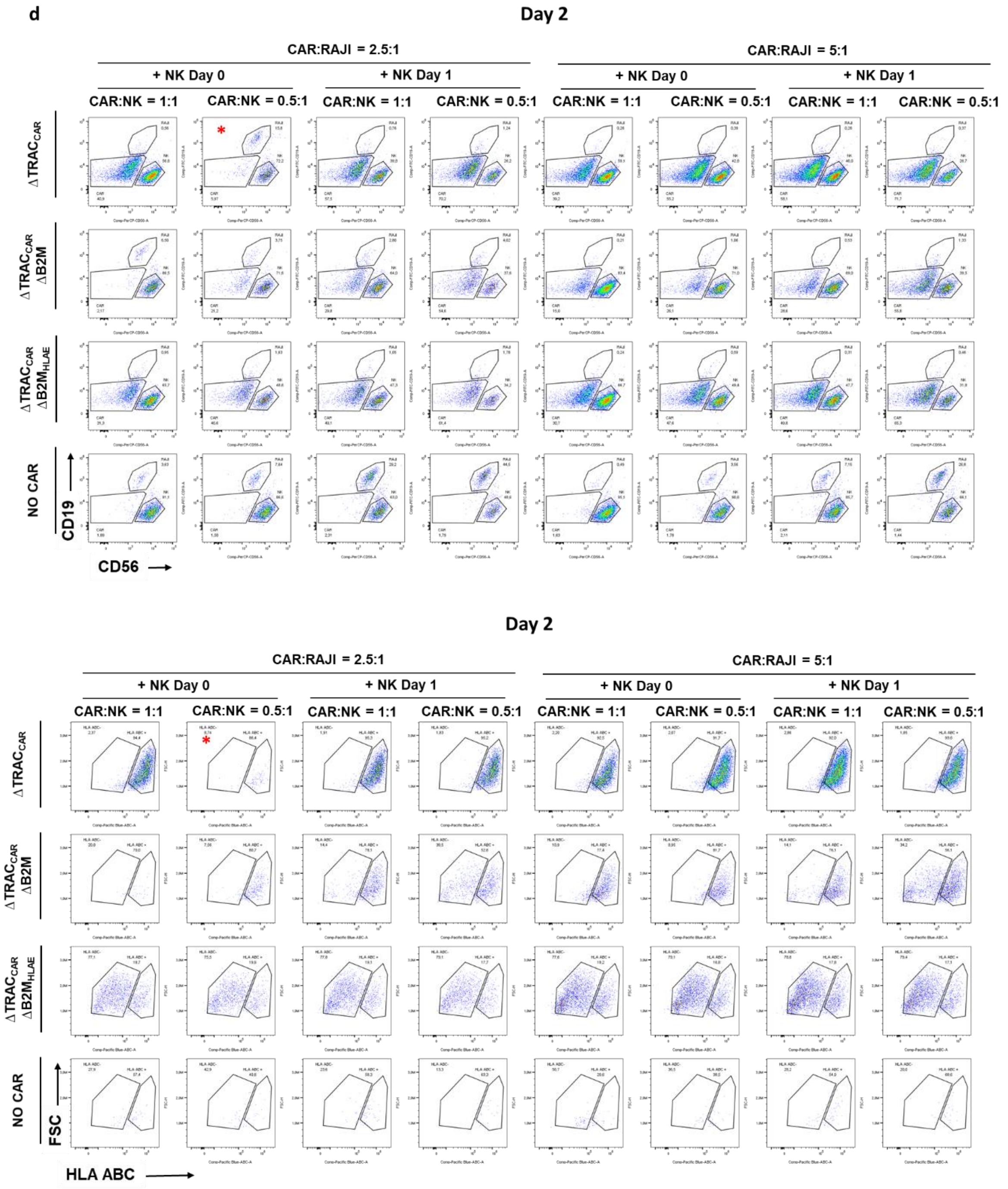
Day 2
CAR:RAJI = 2.5:1
CAR:RAJI = 5:1
+ NK Day 0
+ NK Day 1
CAR:NK = 1:1
CAR:NK = 0.5:1
ΔTRACCAR
ΔTRACCAR ΔB2M
ΔTRACCAR ΔB2MHLAE
NO CAR
CD19
CD56
Day 2
CAR:RAJI = 2.5:1
CAR:RAJI = 5:1
+ NK Day 0
+ NK Day 1
CAR:NK = 1:1
CAR:NK = 0.5:1
ΔTRACCAR
ΔTRACCAR ΔB2M
ΔTRACCAR ΔB2MHLAE
NO CAR
FSC
HLA ABC

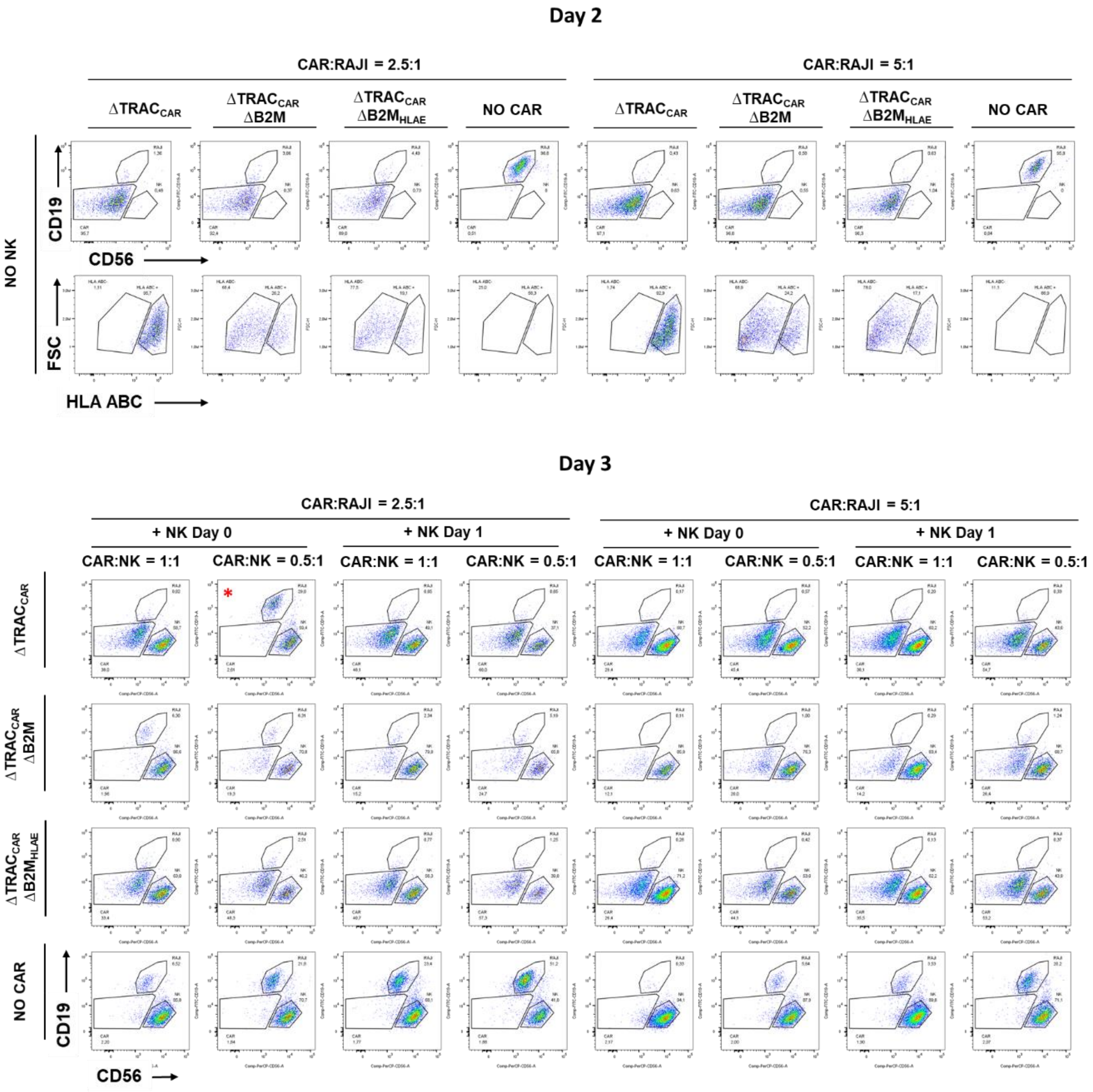
Day 2
CAR:RAJI = 2.5:1
CAR:RAJI = 5:1
ΔTRAC$_{CAR}$
ΔTRAC$_{CAR}$ ΔB2M
ΔTRAC$_{CAR}$ ΔB2M$_{HLAE}$
NO CAR
NO NK
CD19
CD56
FSC
HLA ABC
Day 3
CAR:RAJI = 2.5:1
CAR:RAJI = 5:1
+ NK Day 0
+ NK Day 1
CAR:NK = 1:1
CAR:NK = 0.5:1
ΔTRAC$_{CAR}$
ΔTRAC$_{CAR}$ ΔB2M
ΔTRAC$_{CAR}$ ΔB2M$_{HLAE}$
NO CAR
CD19
CD56

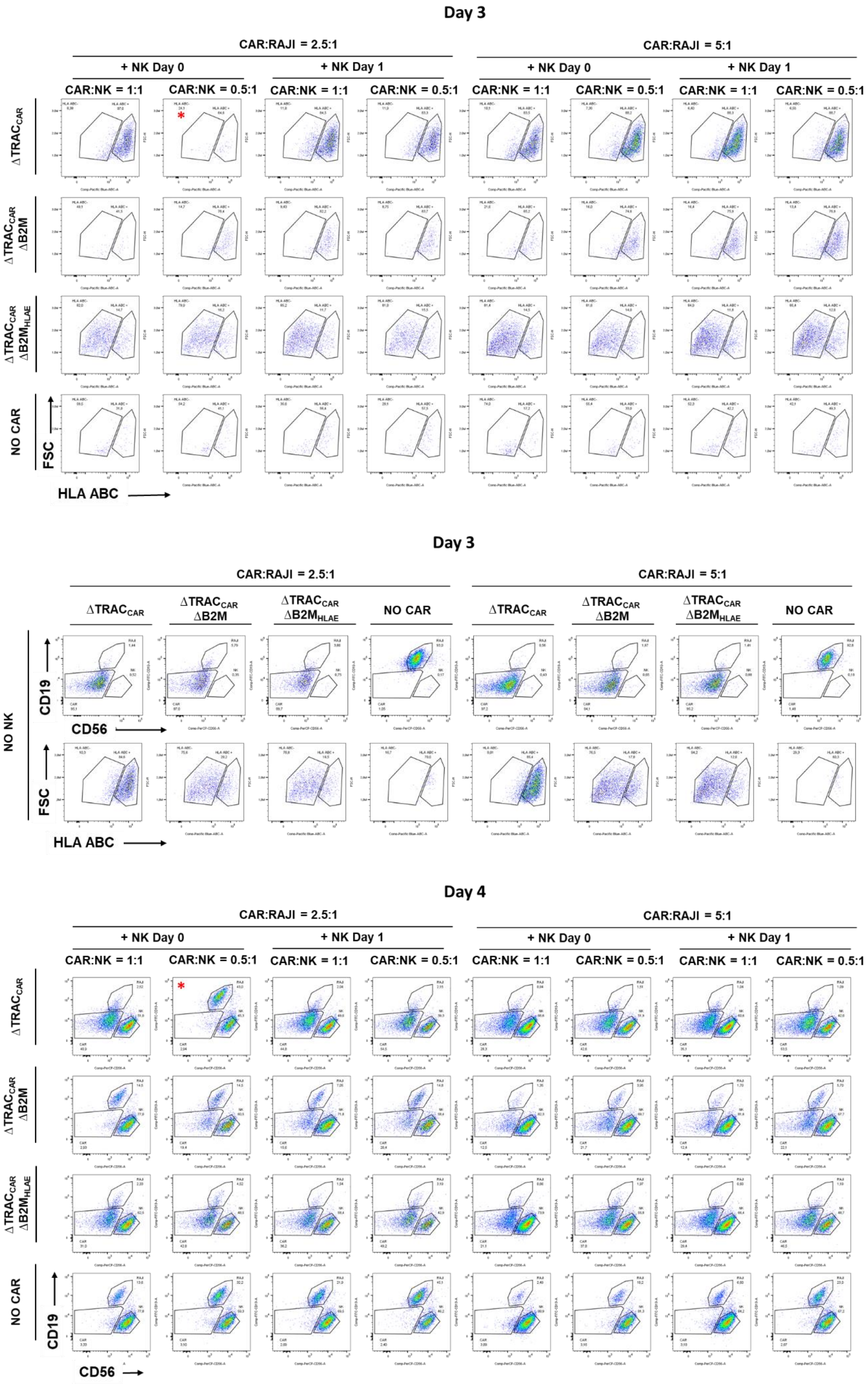
Day 3
CAR:RAJI = 2.5:1
CAR:RAJI = 5:1
+ NK Day 0
+ NK Day 1
CAR:NK = 1:1
CAR:NK = 0.5:1
ΔTRAC_CAR
ΔTRAC_CAR ΔB2M
ΔTRAC_CAR ΔB2M_HLAE
NO CAR
FSC
HLA ABC
Day 3
CAR:RAJI = 2.5:1
CAR:RAJI = 5:1
ΔTRAC_CAR
ΔTRAC_CAR ΔB2M
ΔTRAC_CAR ΔB2M_HLAE
NO CAR
NO NK
CD19
CD56
FSC
HLA ABC
Day 4
CAR:RAJI = 2.5:1
CAR:RAJI = 5:1
+ NK Day 0
+ NK Day 1
CAR:NK = 1:1
CAR:NK = 0.5:1
ΔTRAC_CAR
ΔTRAC_CAR ΔB2M
ΔTRAC_CAR ΔB2M_HLAE
NO CAR
CD19
CD56

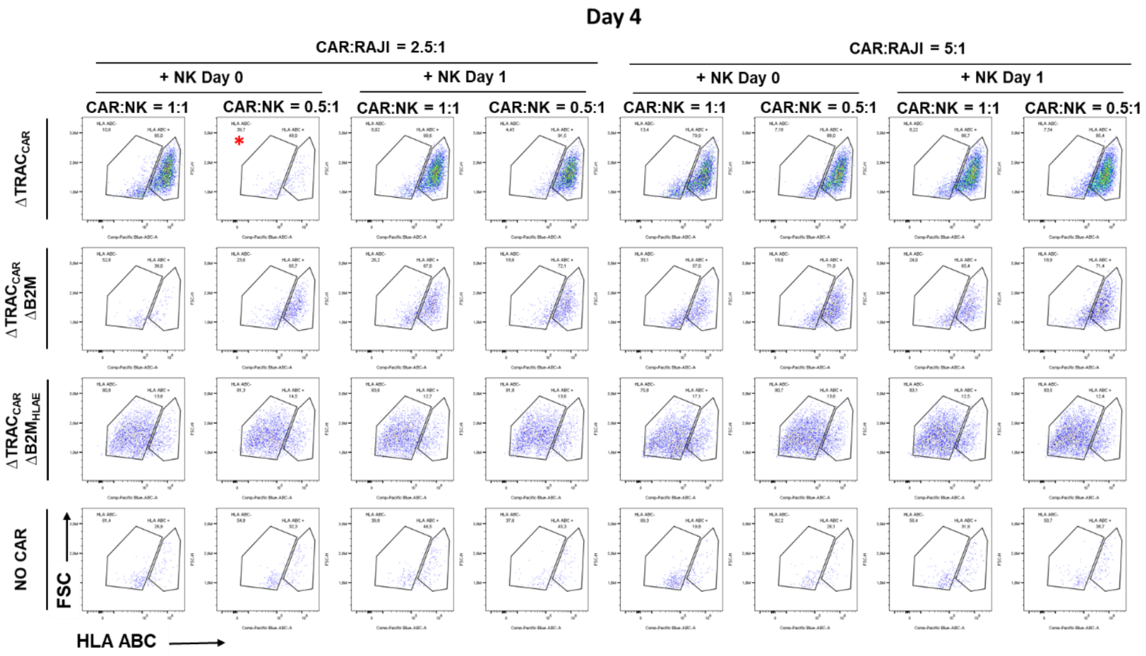
Day 4
CAR:RAJI = 2.5:1
CAR:RAJI = 5:1
+ NK Day 0
+ NK Day 1
CAR:NK = 1:1
CAR:NK = 0.5:1
ΔTRAC$_{CAR}$
ΔTRAC$_{CAR}$ ΔB2M
ΔTRAC$_{CAR}$ ΔB2M$_{HLAE}$
NO CAR
FSC
HLA ABC

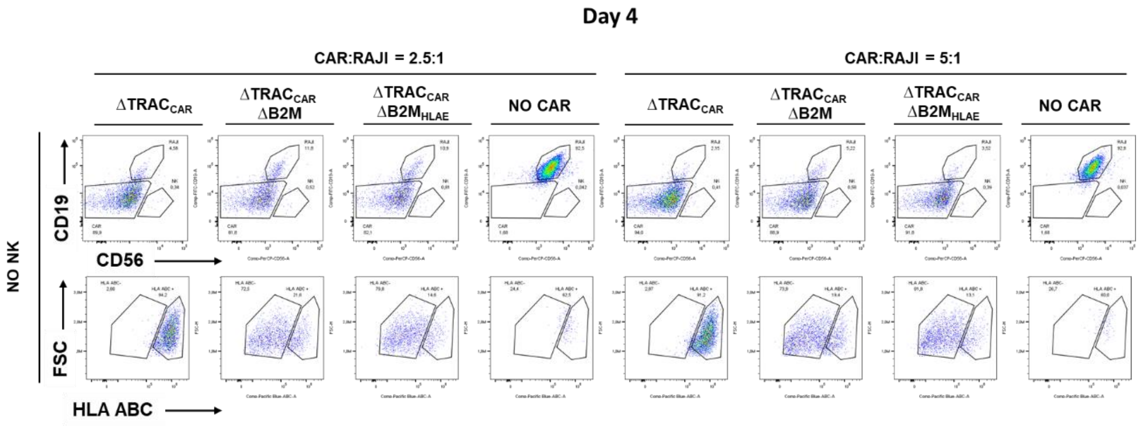
Day 4
CAR:RAJI = 2.5:1
CAR:RAJI = 5:1
ΔTRAC$_{CAR}$
ΔTRAC$_{CAR}$ ΔB2M
ΔTRAC$_{CAR}$ ΔB2M$_{HLAE}$
NO CAR
NO NK
CD19
CD56
FSC
HLA ABC

**d**

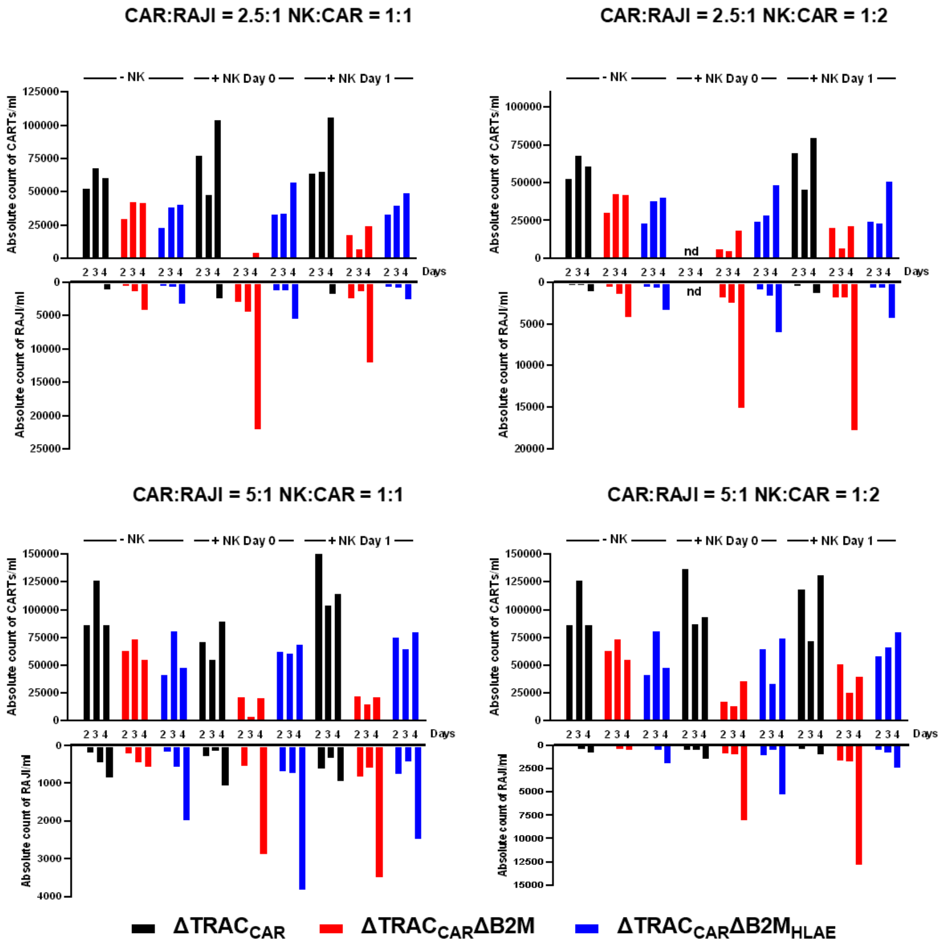


**Supplementary figure 5. Targeted insertion of HLA-E at the B2M locus enables efficient and prolonged antitumor activity of ΔTRAC$_{CAR}$ΔB2M$_{HLAE}$ in the presence of activated NK cells *in vitro*. a** Schematic showing the serial killing assay designed to investigate the long term antitumor activity of ΔTRAC$_{CAR22}$ΔB2M$_{HLAE}$ cells toward RAJI-Luc cells in the presence or absence of activated NK cells. The three different serial killing assay scenarios investigated are illustrated (No NK, NK Day 0 and NK Day 1). Black, blue and red arrows indicate addition of RAJI cells, CAR T-cells (ΔTRAC$_{CAR22}$, ΔTRAC$_{CAR22}$ΔB2M and ΔTRAC$_{CAR22}$ΔB2M$_{HLAE}$) and NK cells respectively. Luminescence and flow cytometry analysis of cell populations are indicated at the different measurment time points. **b** Frequency of RAJI-Luc cells killing observed by luminescence with the three different serial killing scenarios performed with a CAR T-cell to RAJI ratio of 2.5:1 and 5:1, in the absence or in the presence of NK cells with a NK to CAR T-cell ratio of 1:1 and 0.5:1. Each point represents one experiment performed with a given T-cell donor. **c** Absolute CAR T-cells and RAJI cells counts obtained by flow cytometry analysis and cell counts in the same condition as in **b**. Source data are provided as a Source Data file.

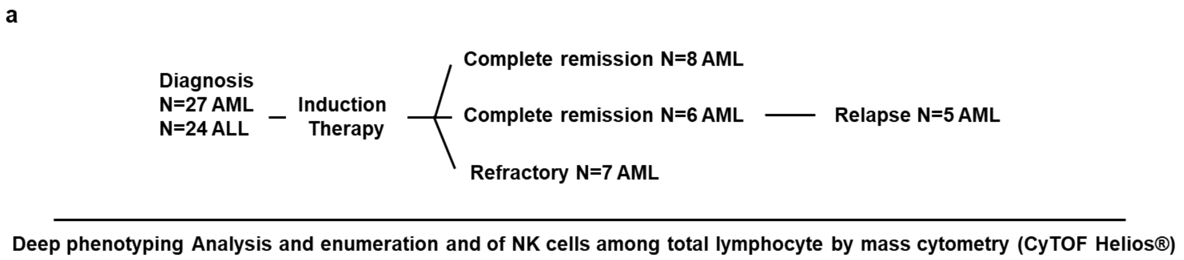


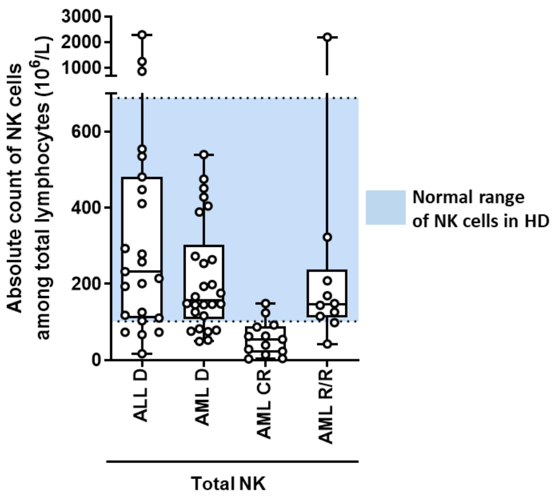


**Supplementary figure 6. Absolute counts of NK cells in AML and ALL patients undergoing standard induction chemotherapy. a** Total lymphocytes from patients with newly-diagnosed ALL and AML as well as AML patients in complete remission and in relapse/refractory disease were characterized by mass cytometry. **b** Absolute counts of NK cells among total lymphocyte obtained for AML and ALL patients at different disease stages. Absolute counts of NK cells were calculated using the frequency of NK cells per total lymphocytes obtained by mass cytometry and the absolute counts of total lymphocytes determined at the time of blood collection. The blue field represents the normal range of absolute counts of NK cell obtained in healthy donors (HD) by flow cytometry [66].

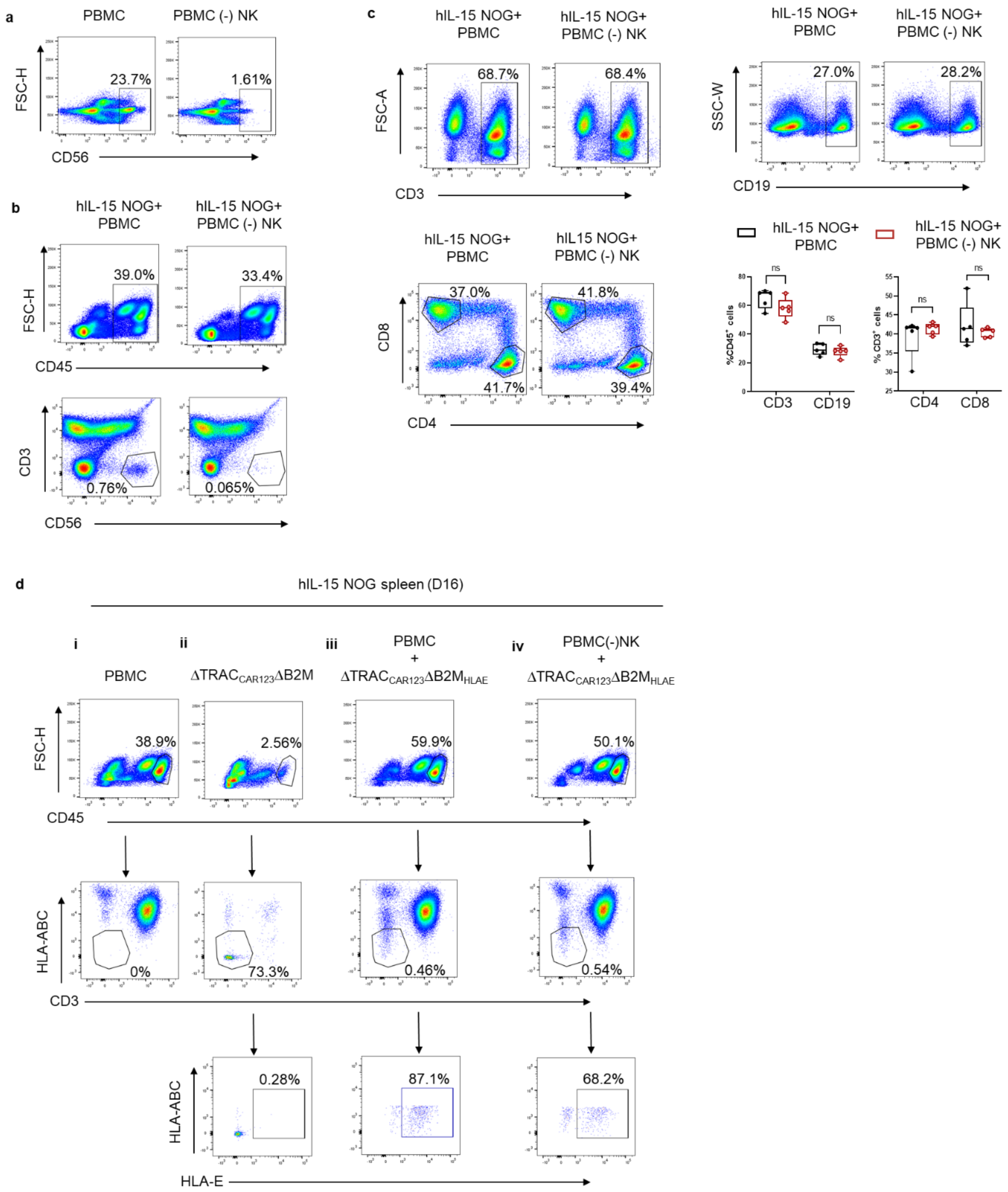


**Supplementary figure 7. Targeted insertion of HLA-E at the B2M locus efficiently prevents NK cell-mediated depletion of ΔTRAC$_{\mathrm{CAR}}$ΔB2M$_{\mathrm{HLAE}}$ cells *in vivo*. a**, Flow cytometry plots depicting the CD56(+) fraction of PBMCs injected into hIL-15 NOG mice with or without CD56(+) cell depletion. Live single cells gated on CD45(+) were analyzed for CD56(+) population in control PBMC samples and NK cell-depleted PBMCs prior to adoptive transfer. **b**, Representative flow cytometry plots depicting CD45(+) immune cell and CD3(-)CD56(+) NK cell engraftment in the spleen of hIL-15 NOG mice, 16 days post adoptive transfer. **c**,

Representative flow cytometry analysis of different immune compartments in the spleen of hIL-15 NOG mice, 16 days post injection of either whole PBMCs or NK cell-depleted PBMCs. **c, bottom right,** Quantitation of the mean value ± s.d. from **c** (n=5). In each box plot, he central mark indicates the median, the bottom and top edges of the box indicate the interquartile range (IQR), and the whiskers represent the maximum and minimum data point. Source data are provided as a Source Data file. **d**, Representative flow cytometry plots depicting gating strategy for identifying injected $\Delta TRAC_{CAR123}\Delta B2M_{HLAE}$ T-cells in the spleen of hIL-15 NOG mice, 4 days post injection. Live single $CD45^{+}$ cells from the spleens of control hIL-15 NOG mice injected with $\Delta TRAC_{CAR123}\Delta B2M$ T-cells alone (i) were gated on CD3(-)HLA-ABC(-) cells-these are the engineered CAR-T cells injected, as evidenced by their absence in the spleen of hIL-15 NOG mice injected with allogeneic PBMCs alone (ii). Illustrated gates were then used to analyze $\Delta TRAC_{CAR123}\Delta B2M$ T-cells for HLA-E expression in samples (iii) and (iv).

### Supplemental Tables

**Supplementary Table 1**. DNA sequences of $CAR_m$, $HLAE_m$ and rlv CAR-2A-HLAE used to package AAV6 and rlv.

| Construct name | DNA sequence and color coding |
|---|---|
| **$CAR_{mCD123}$** | TRAC left homology arm_T2A_Signal peptide_Scfv_CD8 hinge and CD20 mimotopes_Transmembrane domain_41BB_ITAM_Bgh polyA_TRAC Right homology arm<br><br>AAGTAGCCCTGCATTTCAGGTTTCCTTGAGTGGCAGGCCAGGCCTGGCCGTGAACG<br>TTCACTGAAATCATGGCCTCTTGGCCAAGATTGATAGCTTGTGCCTGTCCCTGAGTC<br>CCAGTCCATCACGAGCAGCTGGTTTCTAAGATGCTATTTCCCGTATAAAGCATGAGA<br>CCGTGACTTGCCAGCCCCACAGAGCCCCGCCCTTGTCCATCACTGGCATCTGGACTC<br>CAGCCTGGGTTGGGGCAAAGAGGGAAATGAGATCATGTCCTAACCCTGATCCTCTT<br>GTCCCACAGATATCCAGTCCGGTGAGGGCAGAGGAAGTCTTCTAACATGCGGTGAC<br>GTGGAGGAGAATCCGGGCCCCGGATCCGCTCTGCCCGTCACCGCTCTGCTGCTGCC<br>ACTGGCCCTGCTGCTGCACGCCGCCAGACCCGAAGTCAAACTGGTGGAGTCTGGGG<br>GAGGACTGGTGCAGCCAGGAGGCTCACTGAGCCTGTCCTGCGCCGCTTCCGGCTTC<br>ACCTTCACCGACTACTATATGTCTTGGGTCCGCCAGCCACCTGGGAAGGCTCTGGAG<br>TGGCTGGCACTGATCCGGAGCAAAGCAGATGGATACACCACAGAATATTCTGCCAG<br>TGTGAAGGGCCGCTTCACACTGTCCCGAGACGATTCACAGAGCATTCTGTACCTGCA<br>GATGAACGCTCTGAGACCTGAGGACTCTGCCACTTACTATTGCGCTAGGGATGCAG<br>CCTACTATTCTTACTATAGTCCAGAAGGCGCCATGGACTACTGGGGGCAGGGAACA<br>AGCGTGACTGTCAGCTCCGGAGGAGGAGGATCCGGAGGAGGAGGATCTGGAGGA<br>GGAGGAAGTATGGCTGACTATAAGGATATCGTGATGACCCAGAGCCACAAGTTCAT<br>GTCCACATCTGTGGGCGACCGCGTCAACATTACCTGTAAGGCCTCACAGAATGTGG<br>ATAGCGCCGTCGCTTGGTACCAGCAGAAGCCTGGACAGAGCCCAAAAGCACTGATC<br>TATAGTGCCTCATACCGGTATAGTGGCGTGCCAGACAGATTCACAGGCAGGGGGTC<br>AGGAACTGATTTTACCCTGACAATTTCTAGTGTGCAGGCCGAGGATCTGGCTGTCTA<br>CTATTGCCAGCAGTACTATTCCACCCCCTGGACATTCGGGGGAGGCACAAAGCTGG<br>AAATCAAACGAGGAAGCGGAGGGGGAGGCAGCTGCCCCTACAGCAACCCCAGCCT<br>GTGCAGCGGAGGCGGCGGCAGCTGTCCTTATAGTAATCCAAGCCTGTGTAGCGGC<br>GGAGGGGGTAGCACAACCACACCAGCACCAAGACCACCTACCCCTGCACCAACAAT<br>CGCAAGCCAGCCACTGTCCCTGAGGCCTGAGGCCTGCAGACCAGCAGCAGGAGGA<br>GCAGTGCACACCAGGGGCCTGGATTTTGCCTGCGACATCTATATCTGGGCACCACT<br>GGCCGGAACATGTGGCGTGCTGCTGCTGTCACTGGTCATTACACTGTACTGTAAGC<br>GAGGCCGGAAGAAACTGCTGTATATTTTCAAACAGCCCTTTATGAGACCTGTGCAG<br>ACTACCCAGGAGGAAGACGGCTGCAGCTGTAGGTTCCCCGAGGAAGAGGAAGGCG<br>GGTGTGAGCTGAGGGTCAAGTTTAGCCGCTCCGCAGATGCCCCTGCTTACCAGCAG<br>GGGCAGAATCAGCTGTATAACGAGCTGAATCTGGGACGGAGAGAGGAATACGACG<br>TGCTGGATAAAAGGCGCGGGAGAGACCCCGAAATGGGAGGCAAGCCACGACGGA<br>AAAACCCCCAGGAGGGCCTGTACAATGAACTGCAGAAGGACAAAATGGCAGAGGC<br>CTATAGTGAAATCGGGATGAAGGGAGAGAGAAGGCGCGGCAAAGGGCACGATGG<br>CCTGTACCAGGGGCTGTCTACTGCCACCAAGGACACCTATGATGCTCTGCATATGCA<br>GGCACTGCCTCCAAGGTGATCTAGACTCGAGGTTTAAACCCGCTGATCAGCCTCGAC<br>TGTGCCTTCTAGTTGCCAGCCATCTGTTGTTTGCCCCTCCCCCGTGCCTTCCTTGACCC<br>TGGAAGGTGCCACTCCCACTGTCCTTTCCTAATAAAATGAGGAAATTGCATCGCATT<br>GTCTGAGTAGGTGTCATTCTATTCTGGGGGGTGGGGTGGGGCAGGACAGCAAGGG<br>GGAGGATTGGGAAGACAATAGCAGGCATGCTGGGGATGCGGTGGGCTCTATGACT |

| | |
|---|---|
| | AGTGGCGAATTCCCGTGTACCAGCTGAGAGACTCTAAATCCAGTGACAAGTCTGTCT<br>GCCTATTCACCGATTTTGATTCTCAAACAAATGTGTCACAAAGTAAGGATTCTGATGT<br>GTATATCACAGACAAAACTGTGCTAGACATGAGGTCTATGGACTTCAAGAGCAACA<br>GTGCTGTGGCCTGGAGCAACAAATCTGACTTTGCATGTGCAAACGCCTTCAACAACA<br>GCATTATTCCAGAAGACACCTTCTTCCCCAGCCCAGGTAAGGGCAGCTTTGGTGCCT<br>TCGCAGGCTGTTTCCTTGCTTCAGGAA |
| **HLAE$_m$** | B2M left homology arm_B2M signal sequence_HLA-G peptide_GS Linker_B2M (codon optimized)_GS linker_HLA-E_Bgh polyA B2M Right homology arms<br><br>CGCGCACCCCAGATCGGAGGGCGCCGATGTACAGACAGCAAACTCACCCAGTCTAG<br>TGCATGCCTTCTTAAACATCACGAGACTCTAAGAAAAGGAAACTGAAAACGGGAAA<br>GTCCCTCTCTCTAACCTGGCACTGCGTCGCTGGCTTGGAGACAGGTGACGGTCCCTG<br>CGGGCCTTGTCCTGATTGGCTGGGCACGCGTTTAATATAAGTGGAGGCGTCGCGCT<br>GGCGGGCATTCCTGAAGCTGACAGCATTCGGGCCGAGATGTCTCGCTCCGTGGCCT<br>TAGCTGTGCTCGCGCTACTCTCTCTTAGCGGCCTCGAAGCTGTTATGGCTCCGCGGA<br>CTTTAATTTTAGGTGGTGGCGGATCCGGTGGTGGCGGTTCTGGTGGTGGCGGCTCC<br>ATCCAGCGTACGCCCAAAATTCAAGTCTACAGCCGACATCCTGCAGAGAACGGCAA<br>ATCTAATTTCCTGAACTGCTATGTATCAGGCTTTCACCCTAGCGATATAGAAGTGGA<br>CCTGCTGAAAAACGGAGAGAGGATAGAAAAGGTCGAACACAGCGACCTCTCCTTTT<br>CCAAGGACTGGAGCTTTTATCTTCTGTATTATACTGAATTTACACCCACGGAAAAAG<br>ATGAGTATGCGTGCCGAGTAAACCACGTCACGCTGTCACAGCCCAAAATAGTAAAA<br>TGGGATCGCGACATGGGTGGTGGCGGTTCTGGTGGTGGCGGTAGTGGCGGCGGA<br>GGAAGCGGTGGTGGCGGTTCCGGATCTCACTCCTTGAAGTATTTCCACACTTCCGTG<br>TCCCGGCCCGGCCGCGGGGAGCCCCGCTTCATCTCTGTGGGCTACGTGGACGACAC<br>CCAGTTCGTGCGCTTCGACAACGACGCCGCGAGTCCGAGGATGGTGCCGCGGGCG<br>CCGTGGATGGAGCAGGAGGGGTCAGAGTATTGGGACCGGGAGACACGGAGCGCC<br>AGGGACACCGCACAGATTTTCCGAGTGAACCTGCGGACGCTGCGCGGCTACTACAA<br>TCAGAGCGAGGCCGGGTCTCACACCCTGCAGTGGATGCATGGCTGCGAGCTGGGG<br>CCCGACAGGCGCTTCCTCCGCGGGTATGAACAGTTCGCCTACGACGGCAAGGATTA<br>TCTCACCCTGAATGAGGACCTGCGCTCCTGGACCGCGGTGGACACGGCGGCTCAGA<br>TCTCCGAGCAAAAGTCAAATGATGCCTCTGAGGCGGAGCACCAGAGAGCCTACCTG<br>GAAGACACATGCGTGGAGTGGCTCCACAAATACCTGGAGAAGGGGAAGGAGACGC<br>TGCTTCACCTGGAGCCCCCAAAGACACACGTGACTCACCACCCCATCTCTGACCATG<br>AGGCCACCCTGAGGTGCTGGGCTCTGGGCTTCTACCCTGCGGAGATCACACTGACC<br>TGGCAGCAGGATGGGGAGGGCCATACCCAGGACACGGAGCTCGTGGAGACCAGG<br>CCTGCAGGGGATGGAACCTTCCAGAAGTGGGCAGCTGTGGTGGTGCCTTCTGGAG<br>AGGAGCAGAGATACACGTGCCATGTGCAGCATGAGGGGCTACCCGAGCCCGTCAC<br>CCTGAGATGGAAGCCGGCTTCCCAGCCCACCATCCCCATCGTGGGCATCATTGCTGG<br>CCTGGTTCTCCTTGGATCTGTGGTCTCTGGAGCTGTGGTTGCTGCTGTGATATGGAG<br>GAAGAAGAGCTCAGGTGGAAAAGGAGGGAGCTACTATAAGGCTGAGTGGAGCGA<br>CAGTGCCCAGGGGTCTGAGTCTCACAGCTTGTAACTGTGCCTTCTAGTTGCCAGCCA<br>TCTGTTGTTTGCCCCTCCCCCGTGCCTTCCTTGACCCTGGAAGGTGCCACTCCCACTG<br>TCCTTTCCTAATAAAATGAGGAAATTGCATCGCATTGTCTGAGTAGGTGTCATTCTAT<br>TCTGGGGGGTGGGGTGGGGCAGGACAGCAAGGGGGAGGATTGGGAAGACAATA<br>GCAGGCATGCTGGGGATGCGGTGGGCTCTATGTCTCTTTCTGGCCTGGAGGCTATC<br>CAGCGTGAGTCTCTCCTACCCTCCCGCTCTGGTCCTTCCTCTCCCGCTCTGCACCCTCT<br>GTGGCCCTCGCTGTGCTCTCTCGCTCCGTGACTTCCCTTCTCCAAGTTCTCCTTGGTG<br>GCCCGCCGTGGGGCTAGTCCAGGGCTGGATCTCGGGGAAGCGGCGGGGTGGCCTG<br>GGAGTGGGGAAGGGGGTGCGCACCCGGGACGCGCGCTACTTGCCCCTTTCGGCGG<br>GGAGCAGGGGAGACCTTTGGCCTACGGCGACGGGAGGGTCGGGACAAAG |

| | |
|---|---|
| **CAR-2A-HLA-E** | EF1alpha promoter_Signal peptide_ScFv_CD8 hinge and mimotopes_Transmembrane domain_41BB_P2A_B2M signal sequence_HLAG peptide_GS linker_B2M (codon optimized)_GS linker_HLA-E<br><br>CGTGAGGCTCCGGTGCCCGTCAGTGGGCAGAGCGCACATCGCCCACAGTCCCCGAG<br>AAGTTGGGGGGAGGGGTCGGCAATTGAACCGGTGCCTAGAGAAGGTGGCGCGGG<br>GTAAACTGGGAAAGTGATGTCGTGTACTGGCTCCGCCTTTTTCCCGAGGGTGGGGG<br>AGAACCGTATATAAGTGCAGTAGTCGCCGTGAACGTTCTTTTTCGCAACGGGTTTGC<br>CGCCAGAACACAGGTAAGTGCCGTGTGTGGTTCCCGCGGGCCTGGCCTCTTTACGG<br>GTTATGGCCCTTGCGTGCCTTGAATTACTTCCACGCCCCTGGCTGCAGTACGTGATTC<br>TTGATCCCGAGCTTCGGGTTGGAAGTGGGTGGGAGAGTTCGAGGCCTTGCGCTTAA<br>GGAGCCCCTTCGCCTCGTGCTTGAGTTGAGGCCTGGCCTGGGCGCTGGGGCCGCCG<br>CGTGCGAATCTGGTGGCACCTTCGCGCCTGTCTCGCTGCTTTCGATAAGTCTCTAGC<br>CATTTAAAATTTTTGATGACCTGCTGCGACGCTTTTTTTCTGGCAAGATAGTCTTGTA<br>AATGCGGGCCAAGATCTGCACACTGGTATTTCGGTTTTTGGGGCCGCGGGCGGCGA<br>CGGGGCCCGTGCGTCCCAGCGCACATGTTCGGCGAGGCGGGGCCTGCGAGCGCGG<br>CCACCGAGAATCGGACGGGGGTAGTCTCAAGCTGGCCGGCCTGCTCTGGTGCCTGG<br>CCTCGCGCCGCCGTGTATCGCCCCGCCCTGGGCGGCAAGGCTGGCCCGGTCGGCAC<br>CAGTTGCGTGAGCGGAAAGATGGCCGCTTCCCGGCCCTGCTGCAGGGAGCTCAAAA<br>TGGAGGACGCGGCGCTCGGGAGAGCGGGCGGGTGAGTCACCCACACAAAGGAAA<br>AGGGCCTTTCCGTCCTCAGCCGTCGCTTCATGTGACTCCACGGAGTACCGGGCGCCG<br>TCCAGGCACCTCGATTAGTTCTCGAGCTTTTGGAGTACGTCGTCTTTAGGTTGGGGG<br>GAGGGGTTTTATGCGATGGAGTTTCCCCACACTGAGTGGGTGGAGACTGAAGTTAG<br>GCCAGCTTGGCACTTGATGTAATTCTCCTTGGAATTTGCCCTTTTTGAGTTTGGATCT<br>TGGTTCATTCTCAAGCCTCAGACAGTGGTTCAAAGTTTTTTTCTTCCATTTCAGGTGT<br>CGTGAGTCGACGCCACCATGGCTCTGCCCGTCACCGCTCTGCTGCTGCCACTGGCCC<br>TGCTGCTGCACGCCGCCAGACCCGAAGTCAAACTGGTGGAGTCTGGGGGAGGACT<br>GGTGCAGCCAGGAGGCTCACTGAGCCTGTCCTGCGCCGCTTCCGGCTTCACCTTCAC<br>CGACTACTATATGTCTTGGGTCCGCCAGCCACCTGGGAAGGCTCTGGAGTGGCTGG<br>CACTGATCCGGAGCAAAGCAGATGGATACACCACAGAATATTCTGCCAGTGTGAAG<br>GGCCGCTTCACACTGTCCCGAGACGATTCACAGAGCATTCTGTACCTGCAGATGAAC<br>GCTCTGAGACCTGAGGACTCTGCCACTTACTATTGCGCTAGGGATGCAGCCTACTAT<br>TCTTACTATAGTCCAGAAGGCGCCATGGACTACTGGGGGCAGGGAACAAGCGTGAC<br>TGTCAGCTCCGGAGGAGGAGGATCCGGAGGAGGAGGATCTGGAGGAGGAGGAAG<br>TATGGCTGACTATAAGGATATCGTGATGACCCAGAGCCACAAGTTCATGTCCACATC<br>TGTGGGCGACCGCGTCAACATTACCTGTAAGGCCTCACAGAATGTGGATAGCGCCG<br>TCGCTTGGTACCAGCAGAAGCCTGGACAGAGCCCAAAAGCACTGATCTATAGTGCC<br>TCATACCGGTATAGTGGCGTGCCAGACAGATTCACAGGCAGGGGGTCAGGAACTG<br>ATTTTACCCTGACAATTTCTAGTGTGCAGGCCGAGGATCTGGCTGTCTACTATTGCC<br>AGCAGTACTATTCCACCCCCTGGACATTCGGGGGAGGCACAAAGCTGGAAATCAAA<br>CGAGGAAGCGGAGGGGGAGGCAGCTGCCCCTACAGCAACCCCAGCCTGTGCAGCG<br>GAGGCGGCGGCAGCTGTCCTTATAGTAATCCAAGCCTGTGTAGCGGCGGAGGGGG<br>TAGCACAACCACACCAGCACCAAGACCACCTACCCCTGCACCAACAATCGCAAGCCA<br>GCCACTGTCCCTGAGGCCTGAGGCCTGCAGACCAGCAGCAGGAGGAGCAGTGCAC<br>ACCAGGGGCCTGGATTTTGCCTGCGACATCTATATCTGGGCACCACTGGCCGGAAC<br>ATGTGGCGTGCTGCTGCTGTCACTGGTCATTACACTGTACTGTAAGCGAGGCCGGA<br>AGAAACTGCTGTATATTTTCAAACAGCCCTTTATGAGACCTGTGCAGACTACCCAGG<br>AGGAAGACGGCTGCAGCTGTAGGTTCCCCGAGGAAGAGGAAGGCGGGTGTGAGC<br>TGAGGGTCAAGTTTAGCCGCTCCGCAGATGCCCCTGCTTACCAGCAGGGGCAGAAT<br>CAGCTGTATAACGAGCTGAATCTGGGACGGAGAGAGGAATACGACGTGCTGGATA |

| | |
|---|---|
| | AAAGGCGCGGGAGAGACCCCGAAATGGGAGGCAAGCCACGACGGAAAAACCCCC<br>AGGAGGGCCTGTACAATGAACTGCAGAAGGACAAAATGGCAGAGGCCTATAGTGA<br>AATCGGGATGAAGGGAGAGAGAAGGCGCGGCAAAGGGCACGATGGCCTGTACCA<br>GGGGCTGTCTACTGCCACCAAGGACACCTATGATGCTCTGCATATGCAGGCACTGCC<br>TCCAAGGGGAAGCGGAGCTACTAACTTCAGCCTGCTGAAGCAGGCTGGAGACGTG<br>GAGGAGAACCCTGGACCTATGTCTCGCTCCGTGGCCTTAGCTGTGCTCGCGCTACTC<br>TCTCTTAGCGGCCTCGAAGCTGTTATGGCTCCGCGGACTTTAATTTTAGGTGGTGGC<br>GGATCCGGTGGTGGCGGTTCTGGTGGTGGCGGCTCCATCCAGCGTACGCCCAAAAT<br>TCAAGTCTACAGCCGACATCCTGCAGAGAACGGCAAATCTAATTTCCTGAACTGCTA<br>TGTATCAGGCTTTCACCCTAGCGATATAGAAGTGGACCTGCTGAAAAACGGAGAGA<br>GGATAGAAAAGGTCGAACACAGCGACCTCTCCTTTTCCAAGGACTGGAGCTTTTATC<br>TTCTGTATTATACTGAATTTACACCCACGGAAAAAGATGAGTATGCGTGCCGAGTAA<br>ACCACGTCACGCTGTCACAGCCCAAAATAGTAAAATGGGATCGCGACATGGGTGGT<br>GGCGGTTCTGGTGGTGGCGGTAGTGGCGGCGGAGGAAGCGGTGGTGGCGGTTCC<br>GGATCTCACTCCTTGAAGTATTTCCACACTTCCGTGTCCCGGCCCGGCCGCGGGGAG<br>CCCCGCTTCATCTCTGTGGGCTACGTGGACGACACCCAGTTCGTGCGCTTCGACAAC<br>GACGCCGCGAGTCCGAGGATGGTGCCGCGGGCGCCGTGGATGGAGCAGGAGGGG<br>TCAGAGTATTGGGACCGGGAGACACGGAGCGCCAGGGACACCGCACAGATTTTCC<br>GAGTGAACCTGCGGACGCTGCGCGGCTACTACAATCAGAGCGAGGCCGGGTCTCA<br>CACCCTGCAGTGGATGCATGGCTGCGAGCTGGGGCCAGACAGGCGCTTCCTCCGCG<br>GGTATGAACAGTTCGCCTACGACGGCAAGGATTATCTCACCCTGAATGAGGACCTG<br>CGCTCCTGGACCGCGGTGGACACGGCGGCTCAGATCTCCGAGCAAAAGTCAAATGA<br>TGCCTCTGAGGCGGAGCACCAGAGAGCCTACCTGGAAGACACATGCGTGGAGTGG<br>CTCCACAAATACCTGGAGAAGGGGAAGGAGACGCTGCTTCACCTGGAGCCCCCAAA<br>GACACACGTGACTCACCACCCCATCTCTGACCATGAGGCCACCCTGAGGTGCTGGG<br>CTCTGGGCTTCTACCCTGCGGAGATCACACTGACCTGGCAGCAGGATGGGGAGGGC<br>CATACCCAGGACACGGAGCTCGTGGAGACCAGGCCTGCAGGGGATGGAACCTTCC<br>AGAAGTGGGCAGCTGTGGTGGTGCCTTCTGGAGAGGAGCAGAGATACACGTGCCA<br>TGTGCAGCATGAGGGGCTACCCGAGCCCGTCACCCTGAGATGGAAGCCGGCTTCCC<br>AGCCCACCATCCCCATCGTGGGCATCATTGCTGGCCTGGTTCTCCTTGGATCTGTGG<br>TCTCTGGAGCTGTGGTTGCTGCTGTGATATGGAGGAAGAAGAGCTCAGGTGGAAA<br>AGGAGGGAGCTACTATAAGGCTGAGTGGAGCGACAGTGCCCAGGGGTCTGAGTCT<br>CACAGCTTGTGA |

**Supplementary Table 2**. List of candidate off-site targets identified by OCA analysis. The position (Genome GRCh38/hg38 version) and sequences of candidate off-sites targeted identified by OCA as well as their position within genes are documented.

| Alias | Location | Position GRCh38/hg38 version | Putative hit sequence | Gene (cleavage position) |
|---|---|---|---|---|
| OS1 | chr16 | 17344058-17344112 | TTGCTCCCCAGAGATATTCGGCAATGTCCAGAGACATTTGTGAGACAA | **XYLT1 (Intron)** |
| OS2 | chr14 | 74807773-74807816 | TTGTCAACCAGATAACTGAAAGGAACAGAAGTGTGGAGGCTATCTA | **YLPM1 (Intron)** |
| OS3 | chr1 | 52056997-52057032 | ATAGTTGTCCCCTCGGGAGTGGTAACAATAGTTACCAGTAACTGTGTATCTTCTGTGA | **BTF3L4 (Intron)** |
| OS4 | chr19 | 34927737-34927771 | ATATTGCACAGAAATAGGACCCAGCTTTTTCTTAAGGTTCTTCA | **ZNF30 (Intron)** |
| OS5 | chr10 | 123100518-123100546 | TTCTCCCACAGAACTGGACCACATACTTCTGGGTATTAGTTGAGA | **None** |
| OS6 | chr9 | 136864517-136864560 | TAGTAACACAGAGATGACACATCTCATGGAAGCCTGCAGGCTCTCCA | **EDF1 (Intron)** |
| OS7 | chr17 | 28610946-28610967 | TCTCAGAAGGGACCTCTCTGTTCTAGATGGGGAGGAGATTTGGGGGACCA | **SGK494 (Intron)** |
| OS8 | chr12 | 57535277-57535310 | TAGTACCAAAGAACTGAGGACAGCTAGAGGGCTGGAGGCTCTCCA | **DCTN2 (Intron)** |
| OS9 | chr20 | 58542177-58542452 | ATGTGCGACTGACATTTGGGGCTGAGTCACTGGGGTGGTGGGGACAGATCTTTGGGGCTG | **APCDD1L-AS1 (Intron)** |
| OS10 | chr5 | 168430996-168431008 | TGAATAGCCTCCTCCATAGCTCACTGGTACTCATGCAGGGGCATCTGTGGGATAA | **WWC1 (Intron)** |
| OS11 | chr9 | 34921696-34921731 | GTGTTCCACAGATATACATGGTTTAACATGTGGTATCTGTGGGATAT | **YWHAZP6** |
| OS12 | chr15 | 64422781-64422808 | TAATCCCACAGAAATAAAGTAGGTGGTTTTGCCACATTTGTGGGACCA | **TRIP4 (Intron)** |
| OS13 | chr12 | 121654150-121654180 | TTGACACAAGGGTCAGTGTGGGACAAGAGTGTGGGGTGTGGCCTGTCTCTGTGGGACCA | **MORN3 (Intron)** |
| OS14 | chr1 | 110162385-110162395 | TCTCAACAGATATATGCTAGAGCTGAAGGGGGTCTTGGAGGCCATCCA | **SLC6A17 (Intron)** |
| OS15 | chr8 | 66414344-66414367 | TTATCCTATAAATGTTTTCTAAACTATATGTGTGTGTACCTTCTTATA | **LOC102724687 (Intron)** |
| OS16 | chr7 | 99697290-99697295 | TCTCTGTAGGAAAACCCATTAAGGCTGCTGCTCCCTATGGCAGGCCTGCTTGCAACTCA | **CYP3A51P** |
| OS17 | chr1 | 155549665-155549686 | TACAAGATGATACACATAGGTTATATCCAAGTAGTATACCATTTTATA | **ASH1L (Intron)** |
| OS18 | chr10 | 102067803-102067835 | TGACTACCCTTCAGGCATCAGAACACTCAGGGCCTGGAGGCTTGCTT | **HPS6 (exon non coding sequence)** |
| OS19 | chr7 | 139889800-139889821 | TTGTCCCCCAAAGACCCCACCTTCTAATACTGTCATCTGGGGGTTTA | **TBXAS1 (Intron)** |
| OS20 | chr8 | 73665696-73665726 | TCACTGCTGGTACAAATACAGTCATCCCTAGGTTATCTGTGGGGAAT | **STAU2 (Intron)** |
| OS21 | chr1 | 1398382-1398442 | ATGTTTGTCCCCAGCCACGGCGTCCTGACGGCCGCCAAAGGCTTGAGGGCTATTCA | **CCNL2 (CDS_exon)** |
| OS22 | chr5 | 103834143-103834147 | TGGATATCTCCCAGTAGGGTGTGGAAGGGTTTTTAGAGTAGGGCAA | **LOC105379107 (intron)** |

**Supplementary Table 3**. Sequences of DNA matrices to quantity B2M/TRAC TALEN dependent translocations. A XhoI restriction site (underlined) was added at break points between the B2M and the TRAC sites (bold green and bold blue, respectively) to control for potential contaminations of experimental samples from control matrices

| Name of the translocation | DNA Sequence of matrices used to quantity B2M/TRAC TALEN dependent transloctions. XhoI restriction site (underligned) was added at break points between B2M and TRAC sites (bold green and bold blue, respectively) to control for potential contaminations of experimental samples from control matrices |
|---|---|
| T1 | TATTAAATAAAAGAATAAGCAGTATTATTAAGTAGCCCTGCATTTCAGGTTTCCTTGAGTGGCAGGCCAGGCCTGGCGTGAACGTTCACTGAAATCATGGCCTCTTGGCCAAGATTGATAGCTTGTGCCTGTCCCTGAGTCCCAGTCCATCACGAGCAGCTGGTTTCTAAGATGCTATTTCCCGTATAAAGCATGAGACCGTGACTTGCCAGCCCCACAGAGCCCCGCCCTTGTCCATCACTGGCATCTGGACTCCAGCCTGGGTTGGGGCAAAGAGGGAAATGAGATCATGTCCTAACC**CTGATCCTCTTGTCCCACAGATATAGAACCC**CTCGAG**GAGAGTAGCGCGAGCACAGCTAA**GGCCACGGAGCGAGACATCTCGGCCCGAATGCTGTCAGCTTCAGGAATGCCCGCCAGCGCGACGCCTCCACTTATATTAAACGCGTGCCCAGCCAATCAGGACAAGGCCCGCAGGGACCGTCACCTGTCTCCAAGCCAGCGACGCAGTGCCAGGTTAGAGAGAGGGACTTTCCCGTTTTCAGTTTCCTTTTCTTAGAGTCTCGTGATGTTTAAGAAGGCATGCACTAGACTGGGTGAGTTTGCTGTCTGTACATCGGCGCCCTCCGATCTGGGGTGCGCGCCCCAGCTTGGGACACCGGGCGCTCA |
| T2 | TATTAAATAAAAGAATAAGCAGTATTATTAAGTAGCCCTGCATTTCAGGTTTCCTTGAGTGGCAGGCCAGGCCTGGCGTGAACGTTCACTGAAATCATGGCCTCTTGGCCAAGATTGATAGCTTGTGCCTGTCCCTGAGTCCCAGTCCATCACGAGCAGCTGGTTTCTAAGATGCTATTTCCCGTATAAAGCATGAGACCGTGACTTGCCAGCCCCACAGAGCCCCGCCCTTGTCCATCACTGGCATCTGGACTCCAGCCTGGGTTGGGGCAAAGAGGGAAATGAGATCATGTCCTAACC**CTGATCCTCTTGTCCCACAGATATAGAACCC**CTCGAG**CTTTCTGGCCTGGAGGCTATCCA**GCGTGAGTCTCTCCTACCCTCCCGCTCTGGTCCTTCCTCTCCCGCTCTGCACCCTCTGTGGCCCTCGCTGTGCTCTCTCGCTCCGTGACTTCCCTTCTCCAAGTTCTCCTTGGTGGCCCGCCGTGGGGCTAGTCCAGGGCTGGATCTCGGGGAAGCGGCGGGGTGGCCTGGGAGTGGGGAAGGGGGTGCGCACCCGGGACGCGCGCTACTTGCCCCTTTCGGCGGGGAGCAGGGGAGACCTTTGGCCTACGGCGACGGGAGGGTCGGGACAAAGTTTAGGGCGTCGATAAGCGTCAGAGCGCC |
| T3 | CTCTGGGCAGAACCTGGCCATTCCTGAAGCAAGGAAACAGCCTGCGAAGGCACCAAAGCTGCCCTTACCTGGGCTGGGGAAGAAGGTGTCTTCTGGAATAATGCTGTTGTTGAAGGCGTTTGCACATGCAAAGTCAGATTTGTTGCTCCAGGCCACAGCACTGTTGCTCTTGAAGTCCATAGACCTCATGTCTAGCACAGTTTTGTCTGTGATATACACATCAGAATCCTTACTTTGTGACACATTTGTTTGAGAATCAAAATCGGTGAATAGGCAGACAGACTTGTCACTGGATTTAGAG**TCTCTCAGCTGGTACACGGCAGGGTC**CTCGAG**GAGAGTAGCGCGAGCACAGCTAA**GGCCACGGAGCGAGACATCTCGGCCCGAATGCTGTCAGCTTCAGGAATGCCCGCCAGCGCGACGCCTCCACTTATATTAAACGCGTGCCCAGCCAATCAGGACAAGGCCCGCAGGGACCGTCACCTGTCTCCAAGCCAGCGACGCAGTGCCAGGTTAGAGAGAGGGACTTTCCCGTTTTCAGTTTCCTTTTCTTAGAGTCTCGTGATGTTTAAGAAGGCATGCACTAGACTGGGTGAGTTTGCTGTCTGTACATCGGCGCCCTCCGATCTGGGGTGCGCGCCCCAGCTTGGGACACCGGGCGCTCA |
| T4 | CTCTGGGCAGAACCTGGCCATTCCTGAAGCAAGGAAACAGCCTGCGAAGGCACCAAAGCTGCCCTTACCTGGGCTGGGGAAGAAGGTGTCTTCTGGAATAATGCTGTTGTTGAAGGCGTTTGCACATGCAAAGTCAGATTTGTTGCTCCAGGCCACAGCACTGTTGCTCTTGAAGTCCATAGACCTCATGTCTAGCACAGTTTTGTCTGTGATATACACATCAGAATCCTTACTTTGTGACACATTTGTTTGAGAATCAAAATCGGTGAATAGGCAGACAGACTTGTCACTGGATTTAGAG**TCTCTCAGCTGGTACACGGCAGGGTC**CTCGAG**CTTTCTGGCCTGGAGGCTATCCA**GCGTGAGTCTCTCCTACCCTCCCGCTCTGGTCCTTCCTCTCCCGCTCTGCACCCTCTGTGGCCCTCGCTGTGCTCTCTCGCTCCGTGACTTCCCTTCTCCAAGTTCTCCTTGGTGGCCCGCCGTGGGGCTAGTCCAGGGCTGGATCTCGGGGAAGCGGCGGGGTGGCCTGGGAGTGGGGAAGGGGGTGCGCACCCGGGACGCGCGCTACTTGCCCCTTTCGGCGGGGAGCAGGGGAGACCTTTGGCCTACGGCGACGGGAGGGTCGGGACAAAGTTTAGGGCGTCGATAAGCGTCAGAGCGCC |

**Supplementary Table 4**. List of Translocation specific-oligonucleotides used to quantify potential translocations (T1, T2, T3 and T4) generated by the co-treatment of T-cell with TRAC and B2M TALEN.

| Primer name | Sequence | target |
|---|---|---|
| In-T1R-T3F | GAGACCGTGACTTGCCAGCC | T1 |
| In_B2Mp | TATAAGTGGAGGCGTCGCGC | |
| In-T1R-T3F | GAGACCGTGACTTGCCAGCC | T2 |
| Ex_B2Mq | GAGATCCAGCCCTGGACTAG | |
| Ex-T2R-T4F | AGACCTCATGTCTAGCACAG | T3 |
| Ex_B2Mp | TTGTCCTGATTGGCTGGGCA | |
| Ex-T2R-T4F | AGACCTCATGTCTAGCACAG | T4 |
| Ex_B2Mq | GAGATCCAGCCCTGGACTAG | |

**Supplementary Table 5**. Experimental layout used to investigate the specificity of $CAR_m$ and $HLAE_m$ targeted insertion in T-cells by TLA analysis.

| | **Experiental layout** | | | | **TLA analysis** | | |
|---|---|---|---|---|---|---|---|
| **Sample name** | **TALEN Modifications** | | **Insertion** | | **Locus specific primer set** | | **Construct specific primer set** |
| **Sample 1** | TRAC KO | - | CARm123 | - | Set 1, 2 | - | Set 3, 4 |
| **Sample 2** | TRAC KO | - | - | HLAEm | Set 1, 2 | - | - |
| **Sample 3** | - | B2M KO | - | HLAEm | - | Set 7, 8 | - |
| **Sample 4** | - | B2M KO | CARm123 | - | - | Set 7, 8 | Set 3, 4 |
| **Sample 5** | TRAC KO | B2M KO | CARm123 | - | Set 1, 2 | Set 7, 8 | Set 3, 4 |
| **Sample 6** | TRAC KO | B2M KO | - | HLAEm | Set 1, 2 | Set 7, 8 | - |
| **Sample 7** | TRAC KO | B2M KO | CARm123 | HLAEm | Set 1, 2 | Set 7, 8 | Set 3, 4 |
| **Sample 8** | TRAC KO | B2M KO | rlv CAR-2A-HLA-E | - | Set 1, 2 | Set 7, 8 | Set 9, 10 |

**Supplementary Table 6**. List of primer sets used for TLA analysis of engineered CAR T-cells.

| **Primer set** | **Name/View point** | **Direction** | **Binding position** | **Sequence** |
|---|---|---|---|---|
| 1 | TRAC Locus 1 | Reverse | chr14:23,014,817 | GTGTCAGACTGGAGAAGATC |
| | | Forward | chr14:23,015,055 | CTTCCCAGCAAAGGAACTAT |
| 2 | TRAC Locus 2 | Reverse | chr14:23,018,088 | CTCTTTGCTTTCTCATCCCT |
| | | Forward | chr14:23,018,269 | GGACTCTAGAATGAAGCCAG |
| 3 | CARm123 (Scfv) | Reverse | CARm123 | TGACGGGCAGAGCGGATC |
| | | Forward | CARm123 | ACCTGAGGACTCTGCCACTT |
| 4 | CARm123 (ITAM) | Reverse | CARm123 | AGTGTAATGACCAGTGACAG |
| | | Forward | CARm123 | GGAAGACAATAGCAGGCAT |
| 5 | HLAEm (Locus 1) | Reverse | HLAEm | GAACCGCCACCACCCATG |
| | | Forward | HLAEm | CTGCGCGGCTACTACAAT |
| 6 | HLAEm (Locus 2) | Reverse | HLAEm | GCTTCCATCTCAGGGTGA |
| | | Forward | HLAEm | GGAAGACAATAGCAGGCAT |
| 7 | B2M Locus 1 | Reverse | chr13:45,002,707 | TTAGTGAGAGTTTGGACTGC |
| | | Forward | chr13:45,003,442 | CCTGAAGTCCTAGAATGAGC |
| 8 | B2M Locus 2 | Reverse | chr13:45,008,173 | ATGTATTTGTGCAAGTGCTG |
| | | Forward | chr13:45,008,336 | TGGATTGGTATCTGAGGCTA |
| 9 | rlv CAR-2A-HLA-E (EF1a) | Reverse | CAR-2A-HLA-E | ACTAATCGAGGTGCCTGG |
| | | Forward | CAR-2A-HLA-E | CCATTTCAGGTGTCGTGA |
| 10 | rlv CAR-2A-HLA-E (HLA-E) | Reverse | CAR-2A-HLA-E | GAACCGCCACCACCCATG |
| | | Forward | CAR-2A-HLA-E | CTGCGCGGCTACTACAAT |

**Supplementary Table 7.** Antibodies used to assess expression of relevant markers on the surface of engineered T-cells

| Antibody | Dilution | Fluorophore | Manufacturer | Catalogue # |
|---|---|---|---|---|
| CD56 | 1:50 | APC | Miltenyi | 130-113-310 |
| NKG2A | 1:50 | PE-Vio700 | Miltenyi | 130-113-567 |
| NKG2C | 1:50 | PE | Miltenyi | 130-119-776 |
| HLA ABC | 1:50 | VioBlue | Miltenyi | 130-120-435 |
| HLA E | 1:50 | APC | Miltenyi | 130-117-402 |
| TCR α/β | 1:25 | PE-Vio770 | Miltenyi | 130-109-922 |
| CD3 | 1:50 | PE | Miltenyi | 130-113-139 |
| CD4 | 1:50 | VioBlue | Miltenyi | 130-113-258 |
| CD8 | 1:50 | APC | Miltenyi | 130-110-679 |
| Viability | 1:1000 | e780 | eBiosciences | 65-0865-14 |
| Rituximab | 1:10 | FITC | R&D systems | FAB9575G |

**Supplementary Table 8.** Antibodies and reagents used to assess the cytolytic activity of NK cells present in healthy donors or AML patients' PBMCs and to assess the extent of engineered $\Delta TRAC_{CAR}\Delta B2M_{HLAE}$ target enrichment with our without PBMC effectors.

| Antibodies and Reagents | Dilution | Fluorophore | Manufacturer | Catalogue # |
|---|---|---|---|---|
| Anti-Human CD56 | 1:15 | BV605 | BD Biosciences | 562780 |
| Anti-Human CD45 | 1:12.5 | BV786 | BD Biosciences | 563716 |
| Anti-Human CD3 | 1:25 | PerCP-Vio700 | Miltenyi | 130-113-141 |
| Anti-Human HLA-E | 1:40 | Pe-Vio615 | Miltenyi | 130-117-403 |
| Anti-Human IFN-γ | 1:30 | AF700 | BD Biosciences | 557995 |
| LIVE/DEAD Fixable | 1:100 | Near -IR | Invitrogen | L10119 |
| Celltrace | 1:2500 | Violet dye | Life Technologies | C34557 |
| Golgistop | 1:300 | NA | BD Biosciences | 51-2092KZ |
| Countbright absolute counting beads | 1:100 | APC | Life Technologies | C36950 |
| BD Cytofix/Cytoperm | NA | NA | BD Biosciences | 554714 |
